# Artificial Embodied Circuits Uncover Neural Architectures of Vertebrate Visuomotor Behaviors

**DOI:** 10.1101/2024.12.19.629427

**Authors:** Xiangxiao Liu, Matthew D. Loring, Luca Zunino, Kaitlyn E. Fouke, François A. Longchamp, Alexandre Bernardino, Auke J. Ijspeert, Eva A. Naumann

## Abstract

All brains evolve within specific sensory and physical environments^1^. Traditionally, neuroscience has focused on studying neural circuits in isolation, yet holistic characterization of their function requires integrative brain-body testing^2,3^. To investigate the neural and biomechanical mechanisms of sensorimotor transformations, we constructed realistic neuromechanical simulations (*simZFish*) of the larval zebrafish optomotor response, a visual stabilization behavior^4,5^. By computationally reproducing the body, physical body-water interactions, visual environments, and experimentally derived neural architectures, we closely replicated the behavior of real zebrafish^6^. Through systematic manipulation of physiological and circuit features, impossible in biological experiments, we demonstrate how embodiment shapes neural circuit architecture and behavior. When challenged with novel visual stimuli, *simZFish* predicted neuronal response types, which we identified via calcium imaging in the brain of real zebrafish and used to update the *simZFish* neural network. In virtual rivers, *simZFish* performed rheotaxis by using current-induced optic flow as navigational cues, compensating for the simulated water flow. Finally, a physical robot (*ZBot*) validated the role of embodied sensorimotor circuits in maintaining position in a real river with complex fluid dynamics and visual environments. Together, by iterating between simulations, behavioral observations, neural imaging, and robotic testing, we demonstrate the power of an integrative approach to investigating sensorimotor processing.

**Research Highlights:**

- Developed *simZFish*, an open-source neuromechanical simulation modeling the zebrafish optomotor response (OMR).
- Demonstrated that *simZFish* visuomotor neural circuits are sufficient to maintain position in virtual water currents without additional sensory modalities.
- Investigated how variations in eye geometry influence neural circuit functionality and behavior.
- Conducted optic flow analysis of simulated visual input to identify retinal connectivity requirements, uncovering the factors that shape pretectal neurons’ preference for the lower posterior visual field.
- *simZFish* predicted new neural response types confirmed via real zebrafish calcium imaging.
- Validated *simZFish* circuits with a physical robot, *ZBot,* performing rheotaxis in a natural river with rich visual and complex flow dynamics compared to idealized lab and simulation experiments.

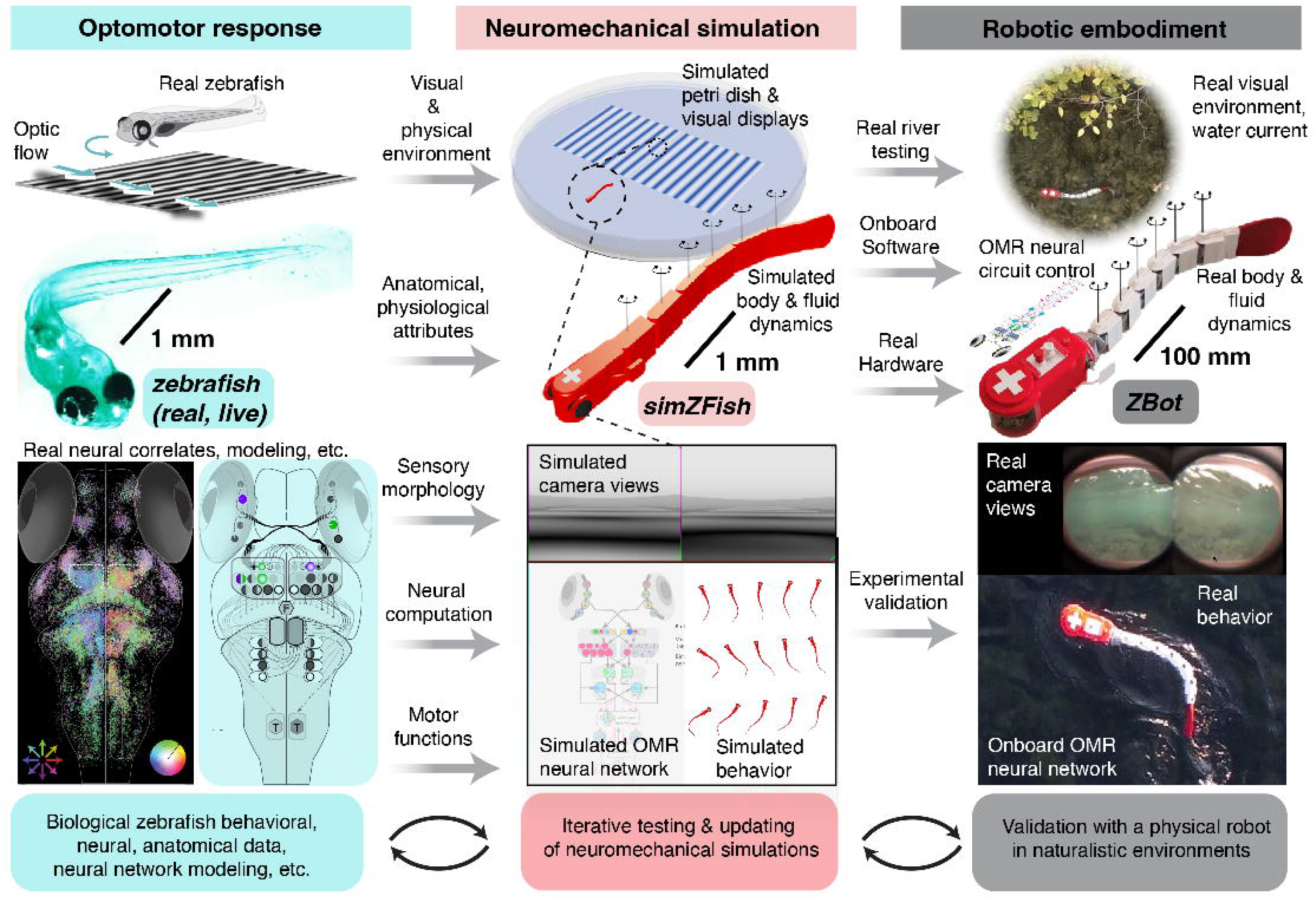

**Graphical abstract:** Combining data-driven neuromechanical simulations and robotic testing, we established an integrative framework for investigating embodied neural circuit functions. Leveraging the behavioral, neurobiological, and theoretical foundations of the visually guided optomotor response in larval zebrafish (blue, left), we developed a virtual zebrafish, *simZFish* (red, middle), to replicate realistic hydrodynamic, anatomical, sensory, neural, and behavioral aspects, permitting the investigation of emergent properties driven by embodiment. Validating these findings using a physical, zebrafish-inspired robot, *ZBot* (grey, right), in naturalistic environments informs further experiments, hypotheses, and robotic design.

## Introduction

The brain is tightly intertwined with the physical attributes of its environment, e.g., hydro or aerodynamics, and its body, e.g., the size and sensory properties^7^. Despite decades of research into sensorimotor systems^8^, we still lack a holistic understanding of how physical body features and environmental factors shape neural computations during behavior^9^, as these systems cannot be fully understood without considering the body in which they evolved^1^. Yet, our insights into how these physical interactions affect sensory processing remain limited due to the challenges of manipulating the physical aspects of animal bodies, controlling sensory environments, and recording neural activity during behavior^10^. These challenges hinder our ability to achieve a comprehensive understanding of the neural architecture requirements and sensorimotor computations in naturalistic settings. To address these challenges, neuromechanical simulations, i.e., numerical simulations of the nervous system, the body, and the environment, as well as real, physical robots, provide powerful tools to integrate multiple components and explore the effects of the environment on neural function and sensorimotor behaviors^2,3,11,12^. Integration of these methods offers untapped opportunities to understand the relevance of embodiment for neural architectures^13–15^, enabling the testing of biologically inspired artificial neural circuits within physical bodies ^15^.

Here, we leverage neuromechanical simulations to investigate the neural control of natural visually guided stabilization behaviors, which are essential for survival and have been identified in many species^16,17^, including mammals^18^, fish^19^, birds^20^, and insects^21^. One such visuomotor behavior is the optomotor response (OMR), which helps animals compensate for displacement caused by moving water^22^ or air^23^. These compensatory actions are highly effective in stabilizing body position and retinal image ^23^, and successfully replicating these neural algorithms represents a decisive advantage to any artificial agent, such as robots^24–26^. However, to accurately model neural dynamics observed in real animals requires validation of sensorimotor loops within the complexities of the physical world and associated nuanced behavioral consequences. Unfortunately, the size, inaccessibility, and complexity of most neuroscience model systems do not offer the information to build complete, biorealistic simulations of visuomotor transformations.

As an important vertebrate model in vision research^27^, the translucent larval zebrafish offers genetic and optical access to almost all neurons^28^. Therefore, we leveraged the larval zebrafish and its well-characterized neural circuits underlying visually guided behaviors^4,6,29–31^ to investigate how embodiment affects neural architecture and function during the OMR. However, current models of zebrafish visuomotor transformations rely on correlation-based neural activity, lack realistic representations or visual perception of the retina or motor systems^6,31^, omit the ability to manipulate neural activity and exclude closed-loop observations of neural activation during behavior, which limits the integrative investigation of the system within its sensory and physical environment.

Therefore, we developed *simZFish*, a biologically inspired neuromechanical simulation of the larval zebrafish designed to investigate how sensory-driven behaviors depend on embodied neural circuits in natural environments. Benefitting from detailed experimental data^6,31–36^, and recent advances in physics simulation technologies^37^, *simZFish* faithfully reproduces the real zebrafish OMR, utilizing computational neurons modeled on experimentally observed neural activity in visuomotor circuits^6^. This simulation replicates the spontaneous and visually evoked neural activity and behaviors of real zebrafish in real experiments, suggesting that it correctly encapsulates the main visuomotor neural circuits underlying OMR. Furthermore, when presented with new simulated environmental challenges such as those experienced in a virtual river, the artificial embodied neural circuits enable *simZFish* to swim against a virtual water flow, performing a behavior known as rheotaxis^22^. Whereas multiple sensory modalities play a role in rheotaxis^22,38^, our work demonstrates that our experimentally derived OMR neural circuit is, in principle, sufficient to perform visually guided rheotaxis without other sensory input. These experiments highlight that *simZFish* is a powerful tool for testing embodied artificial neural circuits in new conditions, identifying the key computational components to achieve behavioral performance, and generating new hypotheses for experimental validation.

In this study, we used *simZFish* simulations to explore sensory-driven neural activity and behavior, revealing how these emerge from the complex interplay between visual stimuli, sensory morphology, and neural circuit architecture. Systematic manipulations of sensory features like the focal length of *simZFish’s* optical lens and neural connectivity, revealed key factors driving neural architecture design. These findings explain the receptive field properties observed in real zebrafish visual processing neurons in the retinorecipient pretectum ^33,39^. Optic flow analysis of the simulated visual input revealed why the lower posterior quadrant of the visual field plays a critical role in the OMR. Additionally, discrepancies in *simZFish’s* behavior when exposed to novel motion patterns predicted the existence of previously unknown functional subtypes of neurons. Guided by *simZFish* predictions, we identified neurons with these previously undiscovered response profiles through calcium imaging in volumetric two-photon microscopy in live zebrafish brains. Incorporating these newly discovered neural subtypes into the *simZFish* model improved its performance, aligning closely with real zebrafish behavior. Finally, *simZFish* directly informed the design of a zebrafish-like robot (*ZBot*), constructed to test whether these embodied artificial neural circuits are sufficient to stabilize its position and orientation in moving waters. Corroborating the results from our neuromechanical simulations, *ZBot* experiments in a natural forest river with rich visual stimuli and complex fluid dynamics confirmed that these artificial OMR circuits are effective in counteracting real water currents. These results underscore how embodiment shapes neural circuits and how neural mechanisms identified in idealized laboratory conditions can be implemented to adaptive behaviors in the real complex world.

By integrating neural and behavioral recordings of real zebrafish, neuromechanical simulations, and robotic validation, this study offers insights into how sensory environments shape neural architecture and presents innovative approaches for robotic control. Our results emphasize the necessity of considering the interaction of all participating components, from sensory to motor systems, brain, body, and environment. Most importantly, our open source *simZFish* platform and physical *ZBot* robot provide a unique opportunity to investigate visuomotor circuits during behavior, providing unprecedented insights into neural circuit function.

Through modeling and investigating artificial neural architectures, our study not only deepens the understanding of embodied neural circuits in the context of visually guided behaviors but also opens new avenues for bio-inspired machine intelligence. Ultimately, this integrated approach lays the foundation for future explorations of how embodied brains evolve to function in dynamic, real-world environments.

### Neural architecture of a simulated zebrafish

To construct realistic, neuromechanical simulations of a 6-day-old larval zebrafish (**Extended Data Fig. 1a**), *simZFish*, we used *Webots* ^40^, a physics-based robot simulator (**Methods**, **Supplementary Methods**). In this simulated reality, the head and body of the *simZFish* are modeled as seven segments connected by six hinge joints actuated by simulated servomotors (**Extended Data Fig. 1b**), with dimensions and masses matching biophysical measurements ^41^ of real zebrafish (**Extended Data Fig. 1c**). The *simZFish* head is equipped with two laterally positioned simulated cameras as the eyes of larval zebrafish (**Fig. 1a**, **Methods**). The *simZFish* platform generates a faithful virtual sensory environment, complete with fluid dynamics, modeling drag, and viscous forces (**Extended Data Fig. 1c**), resulting in realistic locomotion behaviors (**Extended Data Fig. 1d, e**) ^42^. Revealing the internal state of the simulated neural network processing, the *simZFish* interface allows access to the simulated camera views, all neuronal activity patterns, and locomotor variables in this complete, end-to-end simulation from retinal input to motor output (**Video 1**, **Extended Data Fig. 2a**). Only limited by computational time, the nature of the *simZFish* simulation allows us to experiment with an endless number of new, realistic, or unrealistic conditions (**Extended Data Fig. 2b**).

**Fig. 1.**
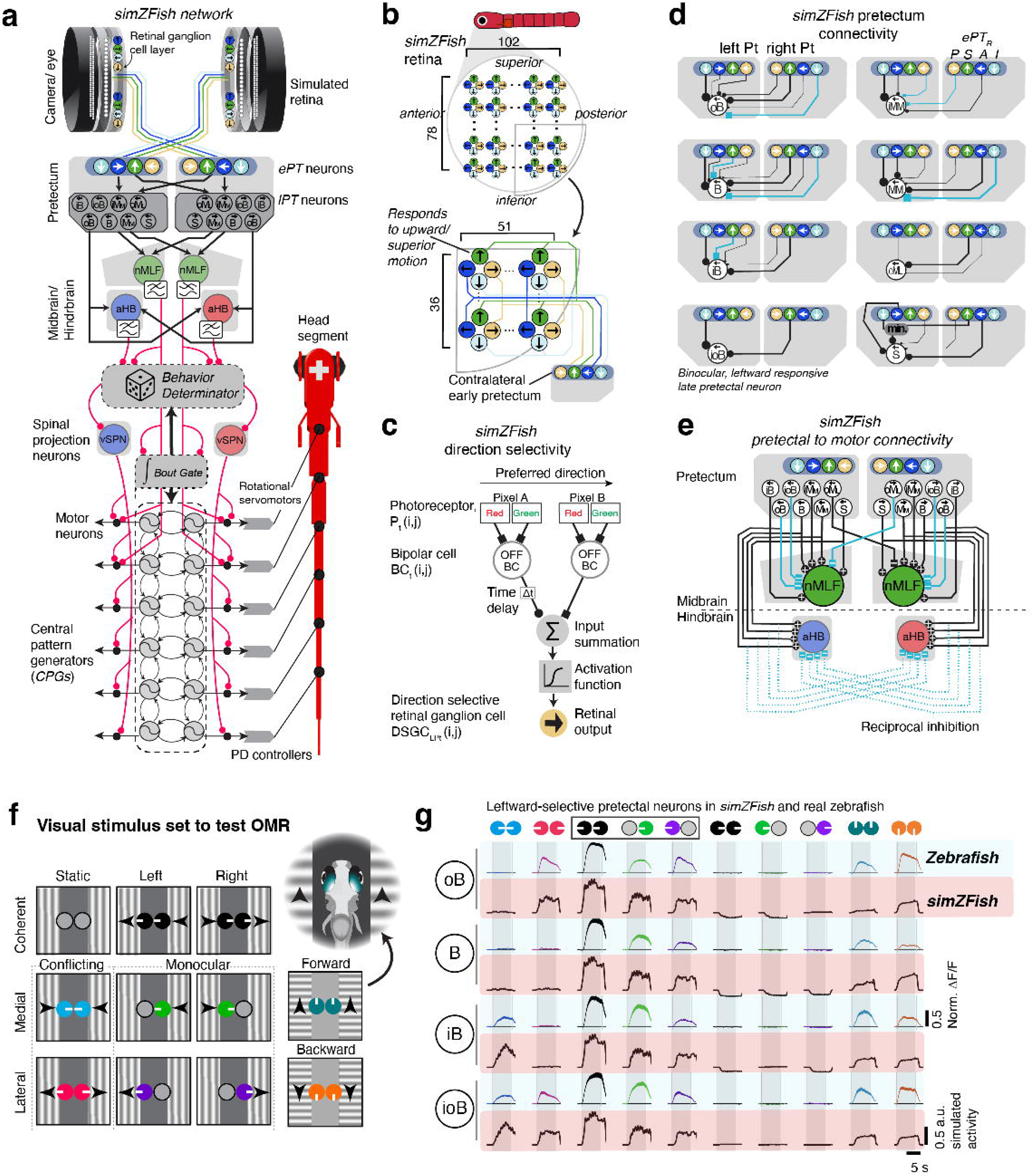
*simZFish*, a neuromechanical simulation replicating the zebrafish optomotor response. **a** *simZFish* neural network. Each node represents an artificial neuron. Motion detection starts in the artificial retina, computing direction selective information relayed to contralateral monocular neurons in the early pretectum (*ePT*). *ePT* neurons activate late pretectal neurons (*lPTs*) which project to the nucleus medial longitudinal fasciculus (*nMLF*) and the anterior hindbrain (aHB). The ‘*Bout Gate*’ integrates inputs from *nMLF* and aHB to activate spinal central pattern generators (*CPGs*) for rhythmic tail movement. The ‘Behavior Determinator’ calculates leftward, forward, or rightward movement probabilities. Motor neurons set the joint positions by integrating the signals from ventromedial spinal projection neurons (*vSPNs)* and *CPGs* (**Supplementary Methods**). **b** Side view of the left *simZFish* retina, consisting of 102 * 78 direction-selective retinal ganglion cells (*DSGCs*), artificial units that respond to the four cardinal directions. Only lower-posterior *DSGCs* project to the *ePT* ^33^. **c** Mechanism of motion direction detection in the *simZFish retina*. Pixel values represent artificial photoreceptors activating downstream bipolar cells, which compute the direction of motion using a classic delay line algorithm, which sums their inputs via a sigmoid transfer function. *DSGCs* selective motion in anterior (A), posterior (P), superior (S), and inferior (I) directions achieve their selectivity by their retinal wiring patterns. **d** Pretectal connections from monocular *ePT* to binocular *lPT* neurons in the left hemisphere. Neuron types: *oB*, outward-responsive binocular; *B*, binocular; *iB*, inward-responsive binocular; *ioB*, inward- and outward-responsive binocular; *iM_M_*, inward-responsive monocular (medial-selective); *M_M_*, monocular (medial-selective); *oM_L_*, outward-responsive monocular (lateral-selective); *S*, (coherent motion selective). Line color indicates excitatory (black) and inhibitory (blue) connections. Line thickness corresponds to normalized connectivity weights. *ePt* selectivity abbreviated as in **2c**. **e** Neuronal connections from binocular *lPT* neurons to *aHB* and *nMLF*. Note that reciprocal inhibition results in stable behavior and suppresses turns to the opposite side of motion direction. **f** Schematic of visual stimulus set composed of static, medial, and lateral motion and forward and backward-moving gratings. Arrowheads indicate the direction of motion, e.g., the inset shows ‘forward’ motion to both eyes. Circular icons represent eyes, and white tick marks show the direction of motion. **g** Average normalized ΔF/F neuronal recordings from pretectal neurons of real, live zebrafish (blue) and *simZFish* (red) for representative left-selective neurons: *oB, B, iB, ioB*. Shaded area, standard error of the mean (SEM) across neurons (**Methods**).

To achieve a realistic artificial neural architecture, we synthesized anatomical ^32,43^, functional ^6,33,34^, and computational findings ^6^, creating an embedded multi-layer neural network model of the OMR visuomotor pathways (**Fig. 1a**). This *simZFish* network comprises rate-coding artificial neurons summing across their inputs via a sigmoid transfer function (**Supplementary Methods**). Visual motion detection starts with simulated photosensitive pixel arrays (**Fig. 1b**) representing the color-sensitive zebrafish retina^44^. Using a classic delay line model ^45^, the *simZFish* retina computes the direction of motion via OFF bipolar cells ^46^ (**Fig. 1c**). This artificial retina drives four types of artificial direction-selective retinal ganglion cells (*DSGCs*) with preferred directions ^47,48^: superior (dorsal), inferior (ventral), anterior (nasal), and posterior (temporal). As in real zebrafish, all *DSGCs* project contralaterally, activating monocular, direction-selective neurons in the early pretectum (*ePT*s) ^34^. These *ePT*s activate neurons in the late pretectum (*lPTs*) ^6,32,35^, creating complex binocular response types (**Fig. 1d**). These neurons exist in real zebrafish in at least eight mirror symmetric sets of overrepresented, direction-selective functional response types ^6^. As suggested by our previous modeling, *lPTs* collectively control the movement direction by activating neurons of the anterior hindbrain (*aHB*) ^6,32^, which also suppresses behavioral oscillation via reciprocal inhibition (**Fig. 1e**).

Like real zebrafish^42^, *simZFish* executes discrete locomotion events, i.e., bouts consisting of rapid tail undulations and a passive glide phase (**Methods**, **Video 2**). To modulate the spontaneous bout frequency (*simZFish*, 0.6 Hz; zebrafish 0.67 Hz +/- 0.09 Hz), the *lPTs* connect to two artificial neural nodes, representing the diencephalic nuclei of the medial longitudinal fasciculus (*nMLF*) known to control bout frequency and swimming vigor ^29^, tail posture ^43^, and bout initialisation ^49^. To generate the characteristic burst and glide behavior ^50^, we implemented abstract neural models of the brainstem and spinal cord motor circuits composed of a ‘*Bout Gate*’ and ‘*Bout Determinator’* centers and artificial swimming central pattern generators (*CPGs)* ^51^. The *Bout Gate* receives input from both *nMLF* neurons, accumulating motion information across both eyes ^6^. If the *Bout Gate* reaches a threshold (**Extended Data Fig. 3a**), it activates the spinal *CPGs* modeled using coupled oscillators ^52^ that generate rhythmic tail undulations ^53^. The transitions between the burst and glide phases are handled by leaky integration at the *Bout Gate*, as suggested previously ^54^. Specifically, a swim bout ends when the *Bout Gate*’s activity decreases to less than 5% of its initial activation level. At the same time, the *Bout Gate* triggers a *Bout Determinator* event, which stochastically defines the turning direction for each bout. Thus, the *Bout Determinator* center replicates the function of a set of bilateral, ventromedial spinal projection neurons (*vSPNs*, **Extended Data Fig. 3b**), which have been demonstrated to control the direction of each bout ^30^. The probability of left, forward, or rightward bouts depends on the activation of *nMLF* and *aHB*. In the *simZFish* simulated spinal cord, the motor neurons determine the body curvature by integrating the output of both *vSPNs* and *CPGs* (**Extended Data Fig. 3c, d**). To compensate for self-generated neural activation during swimming (**Extended Data Fig. 3e, f**), we added low-pass filters, as suggested by their delayed and accumulating activity profiles in real zebrafish^29,31^. Initializing the *simZFish* neural network with our experimentally derived, best-fit ^6^ connection weights (**Methods**, **Extended Data Fig. 3g**), minimal tuning achieved functional connectivity that demonstrated similar neuronal activation patterns to zebrafish neurons to the direction and eye-specific motion patterns we previously tested in real zebrafish (**Fig. 1f**, **g**).

### simZFish replicates OMR behaviors

To compare visually guided behavior in *simZFish* and real zebrafish, we recreated a well-established experimental paradigm to study the OMR ^6^. Placing the *simZFish* in a simulated petri dish (**Fig. 2a**), we tested OMR behavior to drifting sinusoidal gratings (**Methods**). To present motion in a consistent direction, the stimulus display is locked to the body orientations of the *simZFish*, like in closed-loop experiments with real zebrafish ^6^. Comparing general kinematic functionality, *simZFish* performs forward, left, and right swim bouts like real zebrafish (**Fig. 2b, Video 3**). These bouts can be quantified based on distance traveled and angular change (**Fig. 2c**). Overall, *simZFish* recapitulates the characteristic trimodal behavioral distribution of real, live zebrafish, with forward swims (0°+/- 5°) and balanced leftward and rightward routine turns (∼+/-35°) for control, stationary gratings (**Fig. 2d**, **Extended Data Fig. 4a).** Comparing average histograms of bout angle distributions (**Fig. 2e**), both *simZFish* and zebrafish respond to converging motion, medial to both eyes (blue), with increased bout frequency, while diverging motion (red), decreases bout frequency for behavior during stationary gratings. Binocular leftward motion with increased leftward bout frequency (0.027 Hz, -46°), strongly increased biased forward bouts (0.089 Hz, -4°), and suppression of routine turns in the opposite direction (0.012 Hz, 34°). *simZFish* also matches bout angle distributions for monocular presented component motion stimuli, e.g., medial (green) or lateral (purple). As for real zebrafish, forward visual motion increases, while backward motion decreases the number of bouts in *simZFish*. All qualitative aspects (e.g., strong leftward turn and bout increase for leftward stimuli) are well matched within the variability of individual zebrafish. Still, as *simZFish* behavior was tuned to match bout probabilities rather than peak bout frequencies, the bout frequency peak locations differ slightly (**Extended Data Fig. 4b**). Instead, when comparing bout probabilities, i.e., relative proportion of right, forward, and left bouts, we find strong quantitative behavioral correspondence for *simZFish* and real zebrafish, with exceptions for challenges in fitting behavior to backward motion (**Fig. 2f**, **Extended Data Fig. 4c-e**).

**Fig. 2.**
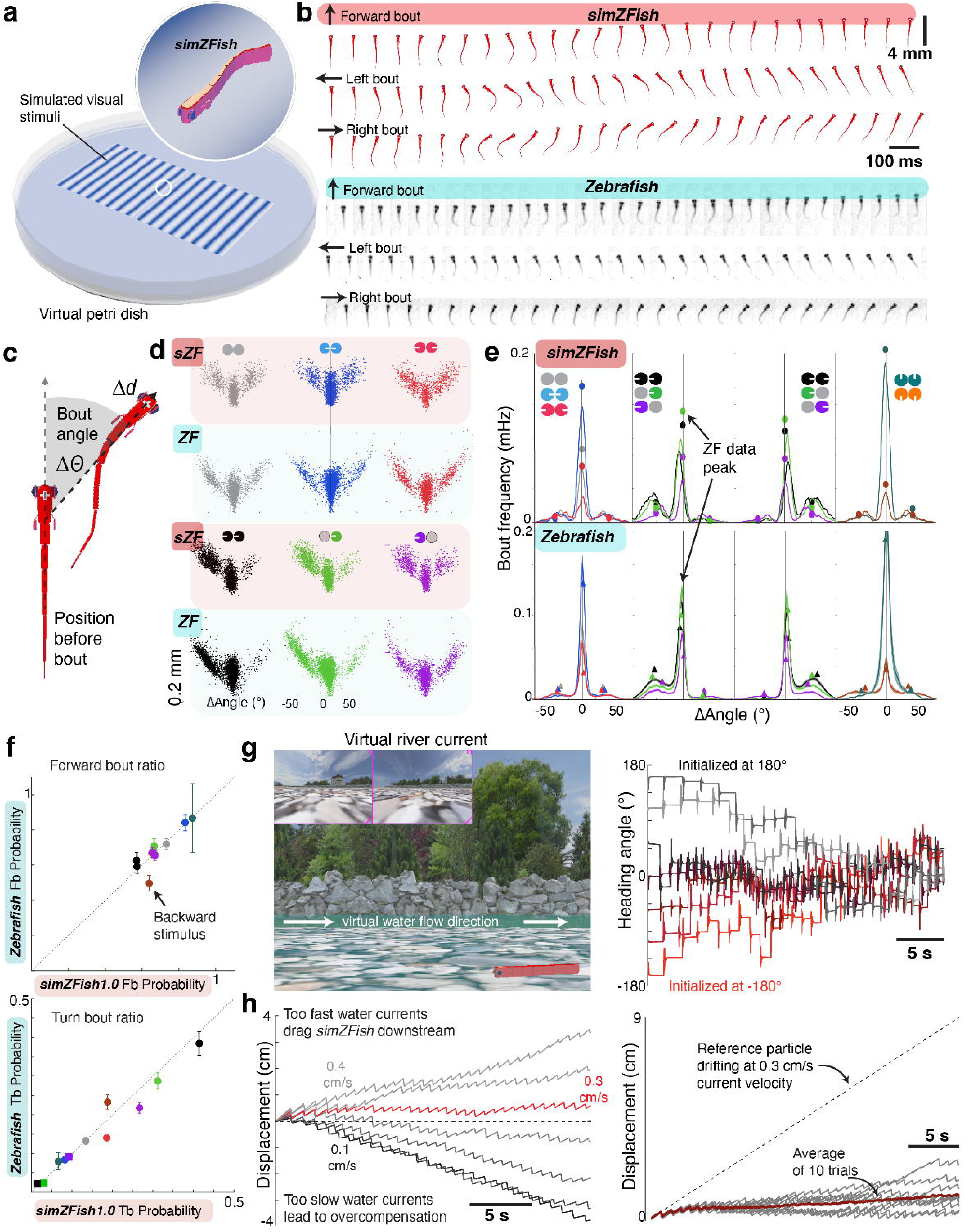
*simZFish* replicates optomotor response of real zebrafish. **a** *simZFish* in a simulated Petri dish exposed to visual stimuli as in real, live zebrafish experiments. **b** *Top*, snapshots of *simZFish* every 100 ms during a representative forward, leftward, and rightward bout. *Bottom*, snapshots of real, live larval zebrafish performing representative forward, leftward, and rightward bouts. **c** Cartoon of *simZFish* performing a single bout, illustrating how bout angle and distance change are measured. **d** Scatterplots of the bout angle and distance distributions for *simZFish* (2000 s) and a representative real zebrafish exposed to visual motion stimuli (20 * 30 s trials), colors as **2f**. Each dot represents a single locomotion event (**Methods**). *simZFish* and real zebrafish exhibit symmetric, trimodal distributions of left, right, and forward bouts to stationary (grey), converging (blue), and diverging (red). For binocular (black, n = 2135 *simZFish* bouts, n = 3295 zebrafish bouts), monocular medial (green), and monocular lateral (purple) leftward stimuli, both *simZFish* and real zebrafish increase turns and biased swim bouts in the direction of motion. **e** Average histograms of bout frequency for *simZFish* and real zebrafish to visual stimuli (symbols as in Fig 1f). Bout distributions of *simZFish* qualitatively replicate the characteristic trimodal zebrafish behavior. Filled circles plotted on the *simZFish* histograms show peak frequencies for real zebrafish’s left, forward, and right bouts at the main modes. *Bottom*, triangles show peak frequencies for *simZFish* on average zebrafish histograms, with shaded error representing SEM across N = 38 fish. *simZFish* replicates general visually evoked behavior patterns with differences in bout angle peaks (*cf.* **Extended Data** Fig. 4, **Methods**). **f** Comparison of normalized bout probability for real zebrafish (y-axis) and *simZFish* (x-axis) for forward bouts (Fb) probability and turn bout (Tb) probability (**Methods**). Each point represents the normalized integrated response for each stimulus. Points close to the diagonal indicate a good match between real zebrafish and *simZFish* data. Error bars represent S.E.M. **g** *Left*, Simulation still image of *simZFish* (red) in a virtual river. *Right*, Heading angle over 30 seconds for different initialized directions (-180°, -135°, -90°, -45°, 0°, 45°, 90°, 135°, 180°). As real zebrafish, *simZFish* first aligns with a few bouts and then swims against the current, an average heading angle of -5.27°, +/-3.03° in the last second. **h** *Left*, displacement over 30 seconds of *simZFish* in a virtual river at different flow speeds. *simZFish* overcompensates for too slow virtual currents but is dragged downstream if currents are too fast. *Right*, *simZFish* maintains its position on average (red) across 10 repetitions of being placed in virtual currents flowing at 0.3 cm/s.

After confirming *simZFish* OMR performance with moving sinusoidal gratings, we tested whether *simZFish’s* OMR neural circuits compensate for virtual water currents with naturalistic visual stimuli, which occur when *simZFish* is dragged in a simulated river with textured bottom (**Fig. 2g**, **Video 4**). When released with different starting heading angles, *simZFish* eventually oriented its heading direction against the flow (**Fig. 2g**, *right*), performing rheotaxis, a typical swimming behavior observed in fish maintain position in water flow. This demonstrates that the *simZFish* neural circuit enables position-stabilizing OMR behaviors, which require repeated alignment with varying optic flow directions to swim upstream.

When presented with different flow velocities, *simZFish* can maintain its position for speeds up to 0.3 cm/s, overcompensating for slower flow velocities but unable to stabilize for faster water flow (**Fig. 2h**). These simulated results show that the *simZFish* OMR algorithm contributes to maintaining position in moving water with visual input alone, effectively performing rheotaxis autonomously. In principle, this suggests that the visuomotor OMR computations can compensate for water current-induced displacement, especially in situations where mechanosensory information from the lateral line organ is not reliable^22^. Importantly, these results not only validate the functionality of our *simZFish* OMR neural network but also provide evidence for OMR’s purpose as a robust sensory-guided stabilization mechanism.

### Effects of embodiment on network function

In principle, while *simZFish* enables us to test infinite numbers of conditions, it can also serve as a tool to systematically explore the effects of manipulations on neural network connectivity, computations, and body morphologies, which may not exist or are difficult —if not impossible— to test in real zebrafish. For instance, we used *simZFish* to measure the effects of altering the sensory morphology, i.e., the properties of its optical lens ^55^. By equipping *simZFish* with lenses of different focal lengths that produced varying field-of-view angles (90°, 120°, 150°)^56^, we investigated how these optical properties affected the perceived image on the artificial retina (**Fig. 3a**, **Extended Data Fig. 5a**, **Video 5**). Since we could precisely enforce *simZFish’s* position in three-dimensional space, we imposed a specific perspective to directly compare lens effects on perception. By examining the visual scene through *simZFish*’s ‘eyes,’ we analyzed the artificial photoreceptor activations. As predicted from optical principles, the focal length affected the perceived visual projection onto the simulated retina with a wider field of view corresponding to shorter focal lengths. When analyzed for optic flow, focal length does not affect areas without contours, such as the simulated, featureless ‘blue sky’ above the horizon, appearing like the sky above for submersed real zebrafish when looking upward through Snell’s window, ∼42° elevation ^57^. During a bright day, this upper visual field contains little information about visual motion generated by being dragged underwater ^58^. However, contoured features, such as river rocks at the bottom, reveal that lenses with shorter focal lengths produce expected perspective exaggeration, rendering closer features larger and those that are more distant smaller (**Video 5**). Therefore, focal length affects the processing of any patterned bottom-projected motion stimuli (**Extended Data Fig. 6a, b**). Narrower angle lenses increase the optic flow magnitude of sinusoidal gratings moving orthogonally to the image plane of the camera or eye (e.g., leftward **Fig. 3a**, *top*) but have minimal impact optic flow magnitude of parallel stimuli (e.g., forward motion, **Fig. 3a**, *bottom*). Analysis of the *simZFish* perceived sensory information shows that focal length mainly changes the amount of measurable optic flow across the visual field for visual stimuli containing orthogonal light-dark edges (**Extended Data Fig. 5b-h**). Calculating the optic flow magnitude, we traced these embodiment effects throughout the entire neural network activity (**Fig. 3b, Extended Data Fig. 6b-f**) and resulting behavior (**Fig. 3c**). Thus, lens morphology directly affects neural activation and sensory-driven behavior even in identical neural networks (**Extended Data Fig. 6c, d**). These results illustrate how neural activation and behavior are emergent properties of embodiment.

**Fig. 3.**
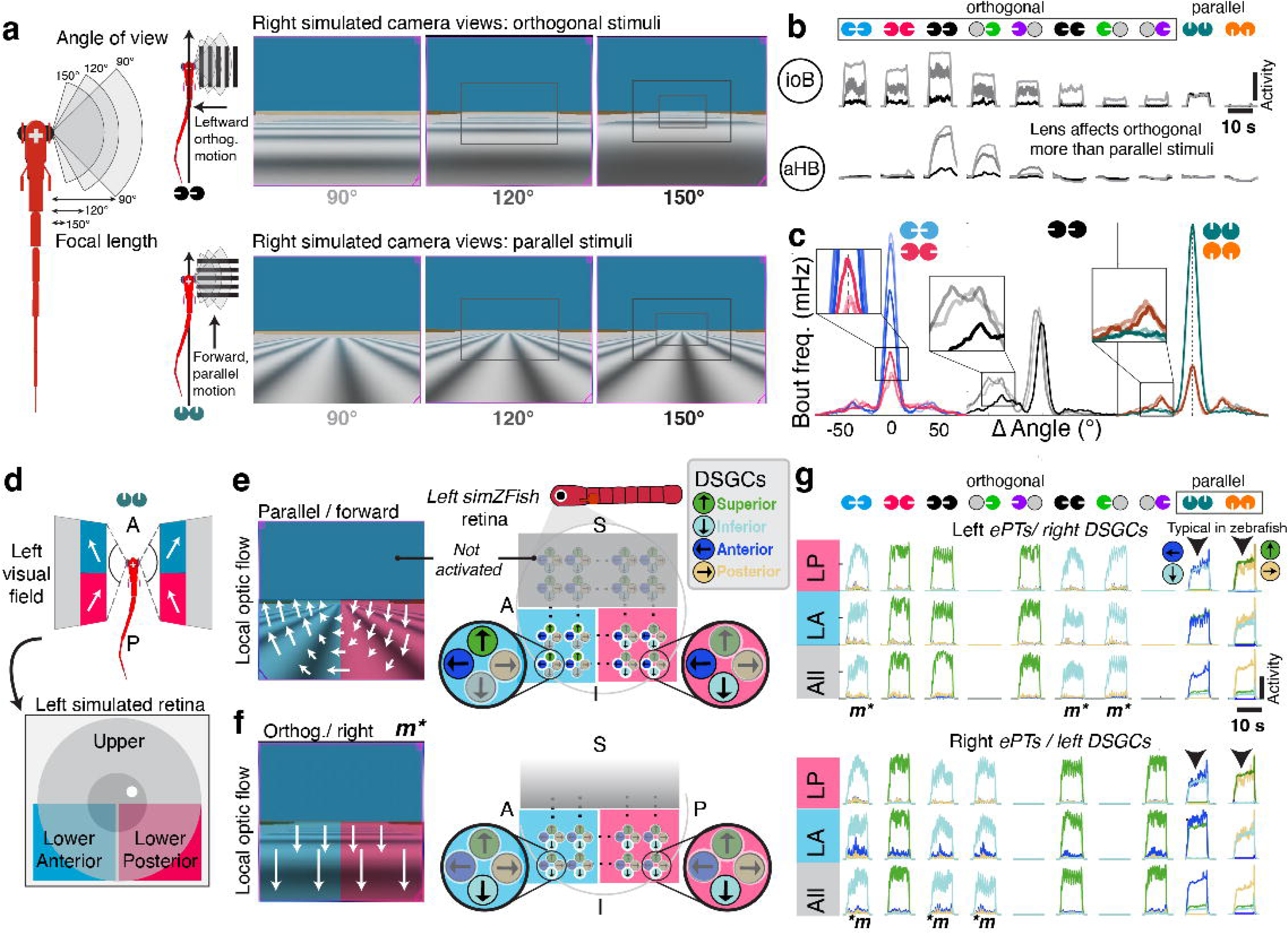
Effects of embodiment on neural activity and visually guided behaviors. **a** *simZFish* can be equipped with optical lenses with different focal lengths. *Right*, Snapshots of right camera recordings while sinusoidal gratings drift in parallel (e.g., forward) or orthogonal (e.g., leftward) to the body axis viewed through lenses with increasing angle of view (rectangular insets), light grey, 90°; grey, 120°; black, 150°. Narrow-angle lenses increase optic flow for orthogonal stimuli (top) but minimally affect parallel stimuli (bottom). **b** Neural recordings of pretectal (*ioB)* and hindbrain (*aHB)* illustrate that changing focal length has specific effects on neural activation, predicted by increased optic flow for narrower angle lenses, e.g., 90° lenses (light grey) increased *ioB* responses to orthogonal stimuli, but not to parallel stimuli. **c** Bout frequency histograms from *simZFish* equipped with different lenses (90°, 120°, 150°; lighter to darker color). OMR behaviors are especially affected for orthogonal stimuli, e.g., 90° lenses led to increased optic flow for these stimuli (**Extended Data** Fig. 5). **d** Dorsal view of *simZFish*’s left and right visual fields, which can be separated into quadrants, defined by their location in the upper, lower, anterior (blue), and posterior (red) areas of the retina. **e** Snapshots of recorded left eye/camera views during parallel forward motion. Perspective distortion leads to rotational optic flow for parallel sinusoidal stimuli. *Right*, forward motion results in rotational optic flow, with the lower anterior (*LA*, blue) retina perceiving upward (superior) and nasal (anterior) and the lower posterior (*LP*, red) perceiving anterior and inferior motion. **f** Snapshots of recorded left eye/camera views during orthogonal sinusoidal stimulation. *Right*, the entire lower visual field perceives downward (inferior) optic flow, thus only activating inferior selective *DSGCs*. **g** Neural recordings of *ePT*s when all (grey), only *LP* (red), or *LA DSGCs* are connected; colors as in **e**. Medial motion (*m\**) drives strong activation of inferior and anterior selective *ePTs* only when the *LP* retina is connected (arrowheads). *LP* connectivity optimally drives OMR as medial/forward coactivates the same *ePts*. These are the most common monocular response types in real zebrafish, e.g., pretectal *iMM* neuron^6^ (*cf.* ‘MoNL’ type neuron in Kubo et al., 2014^34^).

*simZFish* also permits the testing of neural connectivity that does not exist or is difficult to manipulate in real zebrafish. During early tuning, we found that connecting all *DSGCs* across the simulated retina to downstream pretectal targets led to poor OMR performance, with no increased bout frequency in response to forward motion. Optic flow analysis of perceived visual input to *simZFish* suggested that this occurred because the perspective-distorted rotational direction information generated by motion stimuli with parallel gratings cancels each other out (**Fig. 3d**, **Video 6**). This cancellation reduces the network’s ability to differentiate the direction of motion using local motion processing. As *DSGCs* process visual information locally, obfuscating the overall optic flow pattern is a phenomenon known as the ‘aperture problem’ ^59^. Due to perspective distortion of bottom-projected parallel, forward stimuli, the local optic flow co-activates inferior and anterior selective *DSGCs* in the lower posterior (*LP*) retina and superior and anterior selective *DSGCs* in the lower anterior (*LA*) retina (**Fig. 3e**). In contrast, orthogonal stimuli only activate either inferior or superior *DSGCs* across the lower visual field (**Fig. 3f**). Systematically altering the connectivity between *DSGCs* and *ePT*s revealed, the *LP* visual field is most effective for driving simulated OMR, as only this connectivity strongly activated the downstream circuit to drive increased bout frequency (**Extended Data Fig. 6e, f**). Notably, our findings align well with biological studies, which show that the zebrafish OMR is most strongly evoked by motion in the *LP* visual field, matching real zebrafish pretectal neurons with large, lower posterior receptive fields^33^. Moreover, medial/forward motion-responsive neurons are the most frequent monocular response type in real zebrafish (*iMM* ^6^, *cf.,‘*MoNL’ ^34^) which respond to medial (or nasalward) and forward (translational) motion. Therefore, in *simZFish,* we connected only the lower posterior *DSGCs* to drive *ePts*. While it is plausible that alternative neural circuits favoring lower anterior *DSGCs* connectivity could be equally effective, they would require downstream visuomotor circuits different from those experimentally observed. Conceptually, these results suggest that the sensory input—specifically, rotational optic flow patterns in the lower visual field— dictates the architecture of the neural circuits, with lower posterior *DSGCs* driving the OMR circuit. If evolutionary pressure favors minimizing neural connections, this configuration relying on the lower posterior part of the visual field of view provides an effective solution that maximizes optic flow information for downstream neural circuits while minimizing wiring and computational load. Practically, these simulated experiments showcase that *simZFish* serves as a predictive modeling tool to investigate the effects of embodiment on neural processing and behavior. This demonstrates that neural connectivity and ethologically relevant sensory input not only impact neural activity and resulting behavior but might even dictate neural architecture itself ^60^. Thus, our *simZFish* results demonstrate that neural characteristics are emergent properties, challenging to characterize in isolation without complete neuromechanical simulations.

### Simulation predictions drive network improvements (simZFish1.0 to simZfFish2.0)

As shown in **Fig. 2**, the first iteration, *simZFish1.0,* successfully replicates key behavioral aspects of the zebrafish OMR, partly due to its tuning to match previous results^6^ for a limited orthogonal stimulus set (**Fig. 1f**). However, *simZFish1.0,* which is based on neurons classified by their responses to these orthogonal stimuli exhibited difficulties in modeling neural activity and behavior to backward motion (**Fig. 2f**). Using *simZFish* as a predictive framework (**Fig. 4a**), we challenged *simZFish1.0* with an expanded monocular and binocular parallel motion stimulus set (**Fig. 4b**). The model unexpectedly predicted strong turning behavior when tested with these novel stimuli it had not previously encountered, i.e., forward-moving gratings to one eye and backward to the other (*FB*, **Fig. 4b**, **Extended Data Fig. 6c-f**). When we presented these stimuli to real zebrafish (**Fig. 4c**, *middle*), we observed significant discrepancies between *simZFish1.0* and real zebrafish behavior (**Extended Data Fig. 7a, b**), suggesting mismatches in their neural architectures.

**Fig. 4.**
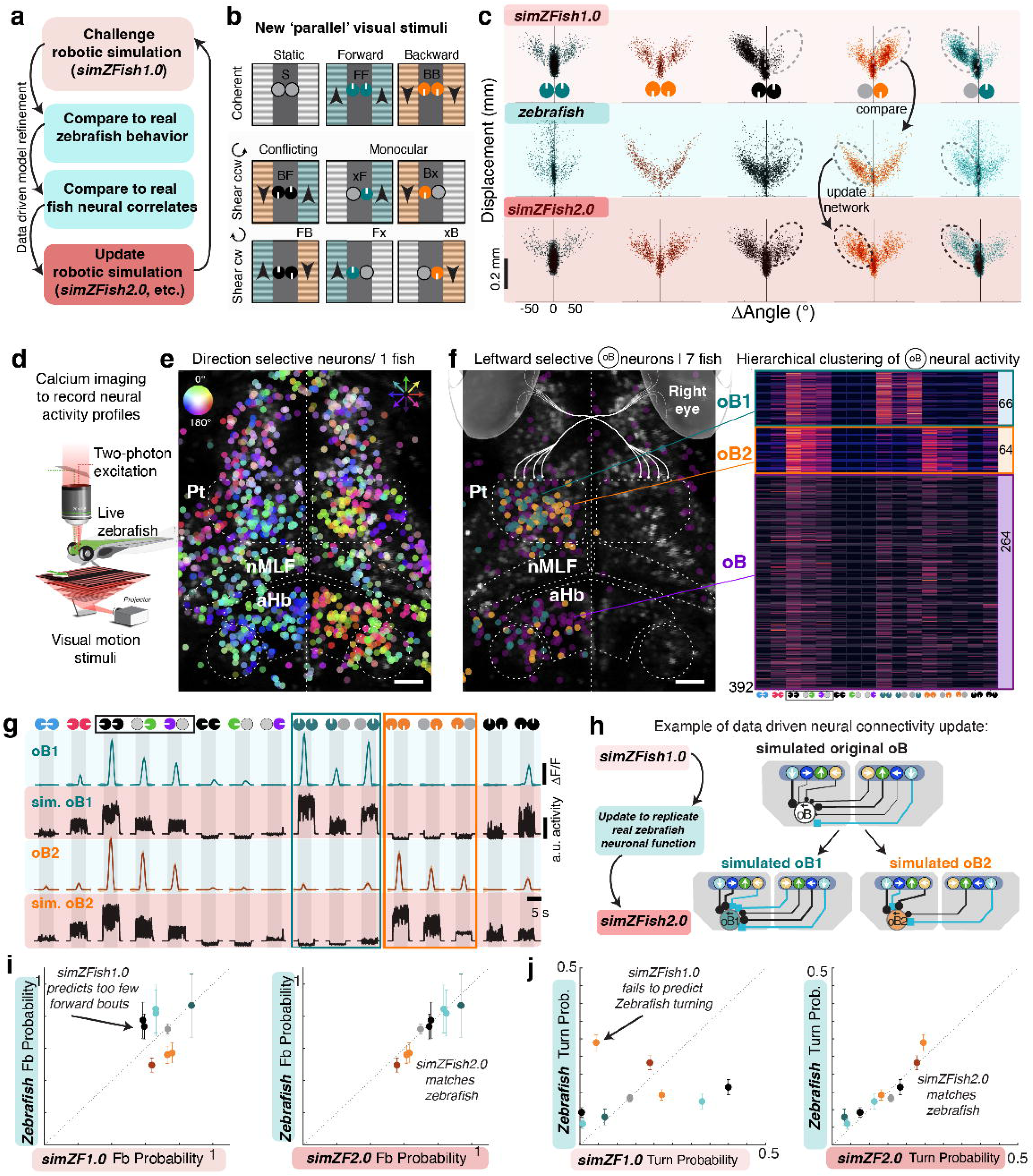
Iterative *simZFish* refinement with new behavioral and neural data. **a** Data-driven strategy to iteratively improve neuromechanical simulations. **b** Monocular and binocular forward (teal) and backward (orange) stimulus combinations move ‘parallel’ to the body axis. Previously, *simZFish1.0* had only encountered binocular *FF* and *BB*. Conflicting, ‘shearing’ combinations of forward and backward (black) to each eye test how these information channels interact. **c** Scatter plots comparing bout distributions in response to ‘parallel’ stimulus set for *simZFish1.0* (light red), representative real zebrafish (blue), and *simZFish2*.0 (darker red). *simZFish1.0* replicates zebrafish behavior to *FF* and *BB* but fails to predict various features of real zebrafish behavior when presented with monocular or conflicting (shearing) stimuli, suggesting specific network interactions of forward and backward stimulus information. Updated *simZFish2.0* matches the behavioral patterns of real zebrafish. **d** Schematic of two-photon microscopy set up to record *in vivo* neural activity from real transgenic *Tg(elavl3:H2B-GCaMP6s)* zebrafish while presenting motion stimuli from below. **e** Two-photon image of *Pt*, *nMLF*, and anterior hindbrain (*aHB*) from a representative zebrafish, overlayed with all detected motion-sensitive neurons as dots. Each dot encodes direction selectivity (hue) and activation level (brightness); see the color wheel (**Methods**). Neurons responding to leftward (blue hues) and rightward (red hues) motion cluster in the left and right hemispheres, respectively. **f** Same brain regions as in **e**, overlaid with all leftward selective *oB* type neurons (*oB_L_*, n = 394, 7 zebrafish). Each is colored based on its class, defined by hierarchical clustering of each neuron’s ΔF/F, shown as a heatmap (right). *oB_L_* neurons contain at least forward (*oB1 _L_*) and backward (*oB2 _L_*) subtypes. **g** Mean, normalized ΔF/F traces of *oB1_L_*, *oB2_L_*in response to the 16 stimuli in real zebrafish (blue) and recordings in simulated *sim oB1 _L_*, *sim oB2 _L_* neurons (red). The similarity shows that we can create matching neural response types in our neuromechanical simulation. Shaded area, SEM across neurons (**Methods**). **h** Data-driven strategy to update the *simZFish1.0* (light red) neural network function for classes that revealed specific subtypes. The neural recordings in real zebrafish inspired the addition of simulated neurons and neurobiological observed connectivity, updating the network to *simZFish2*.0 (darker red). **i** Comparison of measured probability for forward bouts in real zebrafish and *simZFish1.0* (left) and updated *simZFish2.0* (right) in response to the stimulus set. Each point is the mean peak response to a visual stimulus, showing that *simZFish1.0* predicts incorrect bout probabilities, as the neural network relies on incorrect neuronal interactions, repaired in the updated *simZFish2.0* (right) (**Supplementary Methods**). **j** Comparison of measured probability for turn bouts (only right turns are shown) in real zebrafish and *simZFish1.0* (left) and updated *simZFish2.0* (right) with added realistic neurons, repairing the OMR network.

As expected, both *simZFish1.0* and real zebrafish responded to binocular forward motion (*FF*) with increased forward bout frequency. In contrast, binocular backward (*BB*) reduced bout frequency and elicited turns (**Extended Data Fig. 7c-e**). In real zebrafish, monocular forward motion (*xF*, *Fx*) triggered ∼50% of the bout rate observed in binocular *FF*, suggesting near linear integration across eyes. In contrast, *BB* suppressed the bout rate, with monocular backward (*xB*, *Bx*) leading to slightly weaker bout frequency suppression. Notably, monocular backward motion (*xB, Bx)* induced turning in the direction of the stimulated eye, indicating that the brain interprets this as a turning stimulus despite the absence of formal directional information. However, when conflicting forward and backward motion was presented to either eye (*BF*, *FB*), mimicking rotational or shearing motion, real zebrafish bout rate remained near spontaneous levels. Nevertheless, *BF* and *FB* suppressed the strong turning response observed with monocular backward alone (*xB*, *Bx*), suggesting that forward motion suppressed the circuit driving turning. These behavioral results reveal complex interactions across eyes and motion directions in real zebrafish, which were not identified before and, therefore, absent from *simZFish1.0*. We hypothesized that these discrepancies stem from differences in neural response characteristics or connectivity. To improve *simZFish1.0*, we recorded neural activity across the brain via two-photon calcium imaging in response to all 16 parallel and orthogonal stimuli (**Fig. 4d, Methods**). Mirroring the behavior, some backward-selective neurons were suppressed by forward motion to the opposite eye, indicating a neural mechanism that reduces turning (**Extended Data Fig. 8a**). Since these computations were not captured in *simZFish1.0*, we classified all neurons into the eight overrepresented response classes used in *simZFish1.0* (**Fig. 4e**) and applied unbiased hierarchical clustering to reveal subtypes (**Fig. 4f, Extended Data Fig. 8b**). Notably, neurons most critical for turning (*oB, B, Mm*, *S*), split into forward and backward selective types, with additional neurons responding to both and showing suppression to the contralateral eye (**Video 7**). These forward and backward-preferring neural subtypes were prevalent across zebrafish, highlighting their importance in motion processing.

Updating the simulated neural architecture, we incorporated these subtypes into *simZFish1.0* by minimally adjusting network connections to better match the real behavior (**Fig. 4g, Extended Data Fig. 8c**). A quantitative comparison of bout probabilities confirmed that *simZFish2.0* functionally replicates real zebrafish behavior (**Fig. 4i**-**j**). This iterative process enhanced the simulated neural circuit’s accuracy and realism while offering important insights into circuit function. However, bridging the gap between simulation and real-world dynamics requires testing these networks in a physical system.

### Physical robot performs OMR in natural river

The complexity of the real world is challenging to replicate in simulation. For instance, real fish in natural rivers experience far richer visual inputs than we can simulate, including variations in textures, turbidity, and lighting conditions. Real-world fluid dynamics also present challenges, such as varying turbulences and irregular flow patterns. To validate the functionality of the *simZFish* neural network in real-world conditions, we implemented it in a real swimming zebrafish-like physical robot, *ZBot* (**Fig. 5a**).

**Fig. 5.**
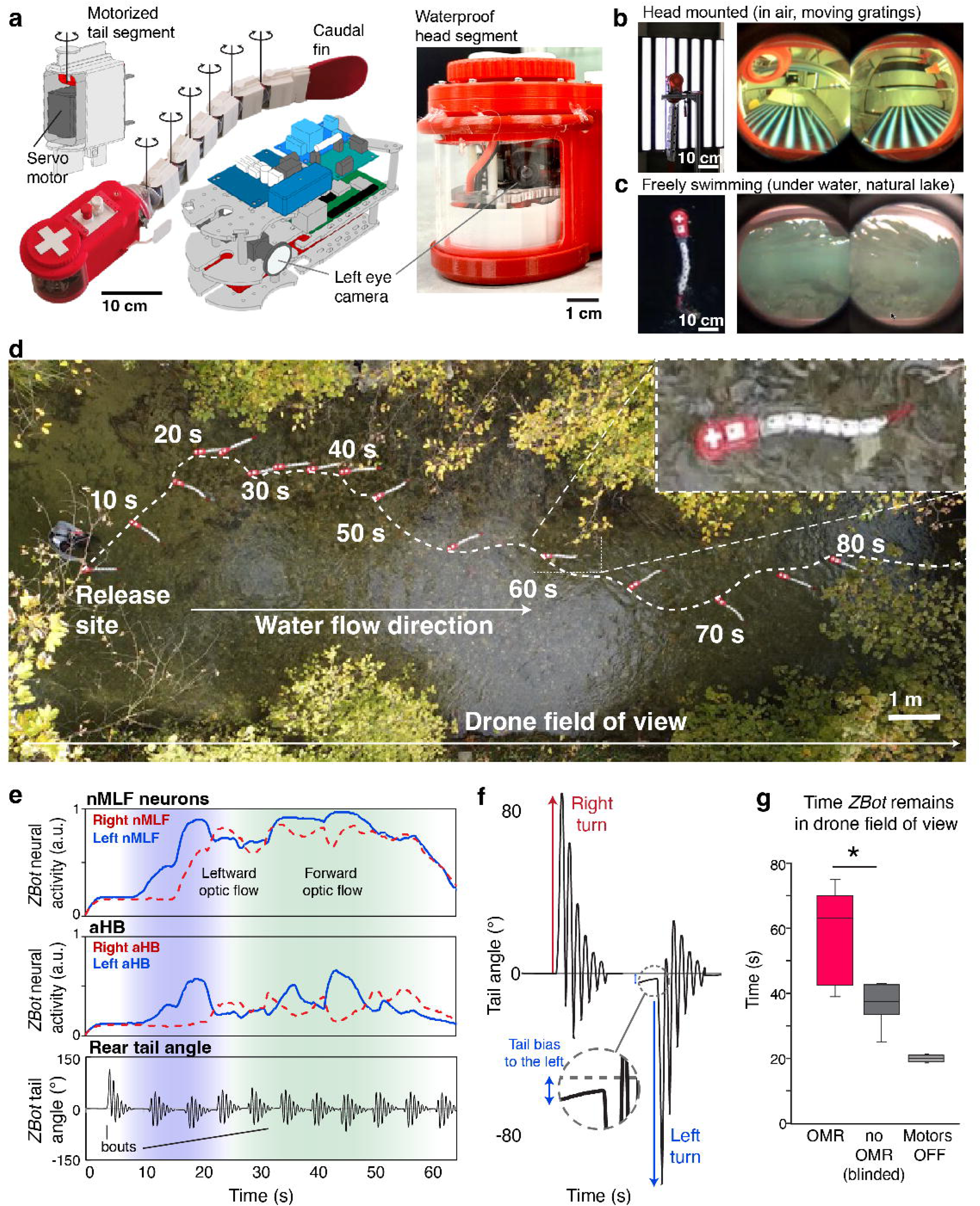
Robot with OMR circuit achieves positional stabilization in natural environments. **a** *Left*, photograph of *ZBot* with 3D renderings of a tail segment with hinge mechanism and the head segment with internal electronic control. *Right*, zoom in on the waterproof head, showing the left camera location. **b** Vertically head fixed *ZBot,* viewing visual stimuli. *Right*, views through *ZBot* cameras demonstrate the quality of visual input under ideal conditions. Note the characteristic perspective distortion effect on orthogonally oriented gratings (forward). **c** Aerial view of *ZBot* swimming in a lake. *Right*, views through *ZBot* cameras show visual input in naturalistic environments. The inferior visual fields contain visual contrast features that activate the OMR mechanism. **d** Drone aerial views of *ZBot* in a natural river, displayed as a composite image with snapshots 5 s apart, zoom in at 60 s. The experimenter releases the *ZBot* with an initial 0° heading direction. As the river’s water flow velocity exceeds the maximal swimming speed of the *ZBot*, even with ideal OMR positional stabilization, the *ZBot* leaves the drone field of view as it is swept downstream in the direction of water flow. **e** Time-aligned graphs of nMLF and aHB activation, and rear tail angle change, corresponding to ***d***. After a spontaneous bout, the *ZBot* drifts rightward (10 to 20 s), thus experiencing leftward optic flow (blue shading), which increases the relative activation of the left nMLF and aHB neurons, induces tail-bending bias to the left and alignment against the water flow direction. From 30 to 60 s, the *ZBot* moves to a water area with increased water flow velocity, experiencing visual forward motion (green shading), increasing activation of *nMLF* and bout frequency. From 65 to 80 s, *ZBot* enters an area with lower ambient light levels since trees block the sunlight, reducing image quality and leading to less *nMLF*, *aHB* activation, and reduced swimming. **f** Zoom-in on an example tail angle change during *ZBot* bouts to the left and right. **g** Bar graphs compare the average time before *ZBot* leaves the drone’s field of view, either with the activated OMR circuit (red, 57.6 s +/- 13.17 s), without OMR by blocking visual input (grey, 37.0 s +/- 5.72 s), or with motors off (light grey, 20.0 s +/- 1.0 s). The *ZBot* stays in view significantly longer with the activated OMR than without, but even random swimming (without OMR) aids positional stabilization. Error bars represent the standard deviation. Undisturbed number of trials: OMR, N = 5, no OMR, N = 8; motors off, N = 2. p = 0.0093, Mann-Whitney U-test for OMR vs. blinded OMR.

While the *ZBot* would ideally replicate the small size (∼4 mm) of the larval zebrafish body, current technology makes it impossible to build such a small robot with the required hardware (cameras, controller, motors, electronic boards, batteries, waterproofing, etc.). Thus, we constructed a larger-scale version (∼ 80 cm) that balances replicating key features of real larval zebrafish (eye position, mass distribution, tail mobility) with technological requirements (**Extended Data Fig. 9b**). The *ZBot* is equipped with two laterally positioned cameras, acting as eyes, a series of servomotors moving its tail segments, and a computational control board that serves as the central sensorimotor processing unit, permitting the *ZBot* to respond to visual input by implementing the same neural circuits as *simZFish* (**Methods**). We tested *ZBot*’s capabilities under three conditions: (1) with visual stimuli while suspended in air, (2) swimming freely in clear, stationary water, and (3) navigating a fast-flowing, shallow river with turbulent, sometimes muddy water.

First, we conducted head-fixed OMR experiments to investigate whether *ZBot*’s visual perception generates neuronal activations comparable to real zebrafish and *simZFish* simulations when presented with moving gratings (**Fig. 5b, Extended Data Fig. 9c, d**). Stimulating the *ZBot* with visual cues displayed on an LCD screen, we recorded videos from the onboard cameras and the activation of its artificial neurons (**Extended Data Fig. 9e**). These controlled experiments demonstrate the strong correspondence between *ZBot* and *simZFish* camera views (**Fig. 5b**, **c**). As expected, we observed comparable neuronal responses across *ZBot*, *simZFish,* and real zebrafish, suggesting that *ZBot* visual processing aligns with that of real zebrafish in these controlled laboratory conditions.

To evaluate the *ZBot’s* swimming capabilities, we first tested it in a shallow lab pool before releasing it into a natural lake environment (**Extended Data Fig. 9f, Video 7**). In the stationary lake water, despite the size differences, *ZBot* exhibited swimming kinematics like real larval zebrafish, demonstrating realistic bout and glide behavior with manually set bout and tail-beating frequencies at 0.09 Hz and 1.5 Hz, respectively. Each tail-beating bout phase lasted ∼5 s, completing five tail-beating cycles (**Extended Data Fig. 9g-h**). However, *ZBot* traveled twice as far per bout as the real zebrafish, likely due to differences in fluid dynamics. *ZBot* operates in a turbulent regime, with a higher Reynolds number (1.35*10^5^ when swimming at a velocity of 0.15 m/s), experiencing lower viscous drag than the live larval zebrafish, which experience intermediate Reynolds numbers: 10 to 1000^61^ and are influenced by both viscous and pressure drag^62^. Because viscous drag is higher than pressure drag during the gliding phase, the comparatively small body of larval zebrafish experiences higher total drag while gliding after removing size and speed effects^62^, resulting in proportionally shorter gliding distances compared to the *ZBot* (**Extended Data Fig. 9i**). Recordings of the *ZBot’s* cameras even in clear, stationary waters emphasized the importance of lower visual fields, as visual features above a certain elevation tend to be blurred and saturated by light (**Fig. 5c**, **Extended Data Fig. 10a**).

To test whether the *ZBot* can use the OMR circuit capabilities to stabilize its position in real water currents, we performed experiments in a natural river with rich visual scenery and irregular water flow due to riverbed features (**Video 9**). We chose a shallow, relatively clear river with sufficiently fast water flow to induce OMR by dragging the *ZBot* downstream. Standardizing experiments, the *ZBot* was always released at the same location, while a remote-controlled drone recorded videos of the *ZBot’s* position from above until the *ZBot* was swept out of the drone field of view (**Fig. 5D**, **Methods**). For each trial, we recorded the *ZBot’s* displacement via the drone, its artificial neuronal activation, and its behavior as tail angle (**Fig. 5e**). The remotely recorded neural activations demonstrate that the natural visual input in the river can activate the circuitry to orient upstream, adjusting heading direction with directed turn bouts and increasing forward bout frequency when aligned the direction of optic flow (**Fig. 5f**). As predicted from our simulations (**Fig. 2g**), the OMR circuit helped the *ZBot* to maintain its position in the river as it remained in the drone’s field of view notably (57%) longer than when its OMR circuit was blinded (cameras off), but spontaneously swimming in bouts in random directions. Further, the active OMR circuit allowed the *ZBot* to maintain its position far (188%) longer than drifting with motors off (**Fig. 5g**). Remarkably, the OMR neural circuit enabled the *ZBot* to turn against the real water current in the direction of the perceived visual flow despite the highly variable and often blurry camera input in the river (**Extended Data Fig. 10b**). Nonetheless, even with an active OMR circuit, the *ZBot* eventually drifted downstream, like the *simZFish* in faster virtual water currents, limited by the maximal swimming speed and bout frequency as *simZFish* and *ZBot* lack any ability for motor adaptation^5^, i.e., adjusting their speed to compensate for faster river flow. Together, these results demonstrate that our sensory-driven, active OMR neural circuitry can aid a physical robot using real-world visual input to reorient its body against the flow to compensate for being dragged downriver.

## Discussion

Solving the design principles of brain sensorimotor systems remains a challenge that cannot be solved with models decoupled from the body and sensory feedback^10,12^. To address this, we developed and analyzed zebrafish-inspired neuromechanical simulations and a physical robot, capturing visually evoked locomotion and neural activity. By synthesizing empirical neural and anatomical data into a physics-based simulator, we created the open-source *simZFish* framework, which accurately reproduces body-water interactions, neural circuits, and visual environments (**Fig. 1**), replicating the experimental settings for live zebrafish behavioral and neural recordings. Equipped with realistic brain-scale neural circuits, from photoreceptors to motor neurons, *simZFish* and the physical *ZBot* produce OMR behaviors akin to live zebrafish (**Fig. 2**). Our findings demonstrate that fish-like simulated and physical bodies when equipped with our artificial OMR circuits successfully compensate for downstream drift, showcasing the utility of our laboratory-derived neural circuits in simulated (**Fig. 2g**) and real-world environments (**Fig. 5d**). Notably, our simulation experiments exposed how embodiment influences neural architecture by permitting access to otherwise hidden variables, such as visual input from the fish’s perspective, highlighting how sensory morphology shapes emergent neural functionality and how the physical properties of visual stimuli dictate optimal neural connectivity (**Fig. 3**). Ultimately, *simZFish* served as a predictive framework that suggested novel behavioral and neural experiments, leading to the identification of neural characteristics critical for biorealistic OMR behaviors (**Fig. 4**). The physical *ZBot* further validated these neural algorithms in the wild, contributing to stabilizing its position in challenging, natural environments with fast-flowing, turbid water conditions (**Fig. 5**), reinforcing its potential for studying sensorimotor systems in real-world environments.

Robots^63^ and neuromechanical simulations^64^ are increasingly used to investigate specific aspects of adaptive animal behavior^10,65^, from insect navigation to lamprey and worm locomotion^10,14,64,66,67^ and sensorimotor coordination in rodents^12^ and humans^68^. Here, however, we model the complete sensorimotor transformation—from raw perceptual signals at the photoreceptors to muscle actuation and the resulting physical and sensory feedback loops. *simZFish* and *ZBot* provide novel tools to investigate vertebrate visuomotor coordination, enabling us to explore how embodied neural circuits perform in vastly different scenarios from those in which they were measured. This integrated approach overcomes the experimental limitations of investigating real zebrafish, allowing the testing of functionalities in virtual conditions that are difficult or impossible to test *in vivo.* While brain scale imaging^5,6,69^ and causal manipulations^70^ via optogenetics are possible in zebrafish, they are costly and technically challenging, particularly when analyzing neural activity during movement^71^. In contrast, simulations are cost-effective and allow for systematic testing of body or neural connectivity modifications ^2,12^. *simZFish* provides full access and control over sensory input and artificial neurons during free locomotion, offering virtually unlimited opportunities to manipulate neuronal properties, from the type and number of neurons and their connection weights to physical attributes like eye position, lens characteristics, or body proportions (see also **Supplementary Text A**).

Beyond replicating lab experiments, we show that *simZFish* circuits are sufficient to reorient upstream and maintain position when placed in virtual water currents (**Fig. 2**). Thus, this relatively complex behavior can emerge from sensing local optic flow, demonstrating that *simZFish* generalized to unseen visual environments, robustly and autonomously performing OMR or visually guided rheotaxis ^38^. Multiple sensory modalities, including vision, the lateral line, tactile sensation, the vestibular system, and proprioception, influence Rheotaxis^38,72^. While the role of vision in rheotaxis is not new, to our knowledge, our study is the first to demonstrate that visuomotor circuits alone can effectively drive rheotaxis in simulation and with a real robot (**Fig. 5**, **Supplementary Text B**). Thus, if visual information is available (e.g., the water is not too turbid), OMR circuits, in principle, are sufficient to perform rheotaxis without other sensory modalities. Demonstrating that vision is sufficient is a challenging and non-trivial result. This is particularly true in animal experiments, where isolating one sensory modality while deactivating all others is rarely possible. Therefore, we show here that the visually driven OMR circuit alone is enabling *ZBot* to navigate upstream.

Beyond validating the biologically inspired neuroanatomy, s*imZFish* provided a powerful tool for systematically analyzing how sensory morphology and neural connectivity shape circuit activity and behavior (**Fig. 3**). By analyzing visual input through the eyes of *simZFish*, we confirmed that optic flow processing is biased to the lower visual field where most contrast and chromatic content is found in underwater habitats^57^. Through controlled manipulation of visual input, lens properties, and retina*-*pretectal connectivity, we revealed how physical attributes and connectivity influence neural activity and, ultimately, neural circuit design. Specifically, our findings explain why pretectal neurons in real zebrafish preferentially respond to motion in the lower posterior (*LP*) visual field^6,32,33^. Our simulations demonstrated that, with laterally positioned eyes, forward-moving, bottom-projected visual stimuli generate opposing rotational optic flow patterns across the visual field (**Fig. 3e**). Restricting retinal-pretectal connectivity to only one quadrant, e.g., *LP* visual field, may thus confer computational advantages, allowing distinct neurons to process forward/medial and backward/lateral motion ^6^. These results, supported by the prevalence of these neural response types in real zebrafish, uncover neural circuit design principles influenced by embodiment (**Supplementary Text C**). Crucially, these insights prompted us to compare the visually evoked behavior of *simZFish* with real zebrafish using an expanded set of monocular and binocular forward/backward stimuli. The discrepancies between the model and actual behavior predicted functional neural subtypes with specific stimulus preferences (**Fig. 4d**). Using volumetric calcium two-photon imaging, we identified a significant set of neurons that matched this predicted response profile. Incorporating these neurons into the circuit improved *simZFish2.0’s* behavioral accuracy, providing a roadmap for how neuromechanical simulations can drive neuroscience research. This iterative interchange between simulation and empirical validation proved a valuable strategy for reverse-engineering visuomotor systems, leading to the discovery of new neural functional subtypes and circuit layouts. As a result, our simulations uncovered previously overlooked, behaviorally relevant neural computations (**Fig. 4**).

Fish-like robots have been designed to address various scientific questions about fish swimming and sensorimotor coordination, for instance, the fluid dynamics of fish swimming^73^, the potential role of tactile feedback^74^, vortex phase matching^75^, tradeoffs between maneuverability and stability ^76^, and the role of the lateral line^77^. In the same spirit, we have designed the physical *ZBot* to investigate OMR in naturalistic settings and validate our circuits in the real world. Moving beyond the approximate nature of simulations, *ZBot* is built with real components (physical visual sensors, actuators, embedded computers) and controlled by the same artificial neural architecture as *simZFish*. For about one hour, *ZBot* operated autonomously, swimming against the current, demonstrating the viability of using resource-efficient and bioinspired algorithms for robotic control in realistic environments.

Other non-fish-like robots have been developed with biologically inspired visual stabilization, e.g., for visually-guided bee flight^25,78^, and even several fish-like robots are equipped with cameras for navigation^63,64,79,80^. Nonetheless, *ZBot* is unique in using a biorealistic neurobiologically derived artificial neural architecture mirroring the real zebrafish brain in function and neuroanatomical design. Investigating the visual input of the *ZBot* demonstrates the complexity of the natural visual input, including turbidity, glare, and Snell’s window, which limits the field of view underwater due to the refraction of light entering the water^81^. Thus, the physical constraints of the underwater environment, as well as those imposed by embodiment, dictate the design of the optimal neural architecture in both real fish and these artificial agents. Our zebrafish-inspired artificial neural architecture with just two laterally positioned eyes may contribute to future robotics design by offering a bio-inspired algorithm for stabilizing fish-like robots in fluid flows without additional cameras. The current methods for position stabilization in floating robots like quadcopters and underwater vehicles often use multiple cameras pointing downward for motion perception^82^. Yet, additional cameras increase the cost and weight, reducing the available payload of the robot. Like real zebrafish, *simZFish* employs only two laterally positioned cameras, providing a physically and computationally lightweight position stabilization method for fish-like robots. Our design requires only a few artificial neurons with minimal connectivity and eliminates the need for extra downward-looking cameras.

Ultimately, our findings demonstrate how neural circuits, body morphology, and environmental context interact to shape neural circuits and adaptive behavior. Future studies should investigate how different sensory modalities, such as exteroception through mechanosensation of the lateral-line organ^83,84^, and proprioception via intraspinal stretch^85^ and cerebral fluid sensors^86^, contribute to behavior modulation. Our physics-based, open-source tools also provide a platform for investigating neural feedback mechanisms and gain control^5,87^ and visually guided behaviors, such as predator avoidance^88^, obstacle navigation^88^, and prey capture^89^.

By reverse engineering sensory gains using end-to-end neuromechanical simulations of real animals exposed to conflicting sensory stimuli, future versions of *simZFish* could incorporate other neurobiological data, such as connectivity mapping via electron microscopic reconstructions^90^, optogenetic manipulations^70^or transsynaptic labeling^91,92^. Moreover, *simZFish* would benefit from machine learning techniques^67,93^, including backpropagation^94^ for supervised learning or proximal policy optimization^95^ for reinforcement learning of target behaviors. Iterating between biorealistic simulations, neurobiological and behavioral experiments, and robotic testing, offers great potential to deepen our understanding of the neuromechanical principles underlying adaptive behavior in biological and artificial agents.

## Methods

### Neuromechanical simulations, *simZFish*

#### simZFish physical body

Our neuromechanical simulation of the larval zebrafish, called *simZFish*, is implemented using the *Webots 2021a* simulator (Cyberbotics Ltd, Switzerland). We chose Webots for several reasons: it is open-source, and it includes a physics engine for simulating body and fluid dynamics, good models of optical cameras, and 3D graphics for displaying behavior. We approximated the real larval zebrafish body dimensions with a length of 4 mm, a height of 0.044 mm, and a weight of 0.0938 mg based on average physical measurements of larval zebrafish^96^. The density of the *simZFish* is slightly lower (91.5 % of the water density) than the water itself. Therefore, the *simZFish* body floats just below the water’s surface, with the cameras fully immersed under simulated water. This ensures that swimming movements remain horizontal, like live, freely swimming larval zebrafish tend to perform^97^. The *simZFish* has seven body segments: the first head segment contains the cameras and bilateral pectoral fins, followed by six body segments, with the last segment connected to a tail fin. All the weights and sizes are based on the measurements of larval zebrafish^96^, matching their overall morphology (**Extended Data Fig. 1a**). Since a sophisticated geometry carries a high computational cost, we simplified the shape of each segment as a cuboid, with a smaller width from head to tail, increasing the simulation speed and simplicity (**Extended Data Fig. 1b**). Each segment is connected by a hinge joint under the actuation of a simulated servomotor, six in total. A servomotor is designed to precisely rotate to the desired joint angle by following the command from the swimming controller. In principle, we could have simulated muscle-like actuation instead, but we chose these simulated servomotors for simplicity and computational speed.

#### simZFish hydrodynamics

In the simulation, each simZFish body segment is subject to static and dynamic forces. The static forces include Archimedes’ thrust. The dynamic forces include both viscous resistance forces and inertial drag forces. Indeed, zebrafish larvae swim at Reynolds numbers between 10 and 1000, corresponding to intermediary regimes where both viscous and inertial forces play an important role ^61^. We tuned the hydrodynamics parameters (inertial drag and viscous resistance force coefficients) to match kinematic recordings of zebrafish swimming (**Extended Data Fig. 1c**). When replaying recorded kinematics from fish (i.e., segment angles) with the servo motors, our *simZFish* can produce forward and routine turning bouts that closely match recorded displacements of the real zebrafish (**Extended Data Fig. 1e**).

#### simZFish simulated neural retinal architecture

To allow the *simZFish* to perceive simulated visual information, the head module includes two simulated color (red, green, blue) cameras. To simulate the anatomical position of real zebrafish eyes, we configured the left and right cameras to face lateral directions, each equipped with a rectilinear lens (**Extended Data Fig. 1b**). While real zebrafish possess wide-angle, ‘fisheye’ lenses with distortion, *simZFish* uses no-distortion lenses measuring 120 degrees along the diagonal, which generates a rectangular view of 96 degrees in horizontal and 78 degrees in vertical directions. Thus, this configuration approximates the real zebrafish’s non-overlapping visual fields^98^. Each camera acquires 1000 frames per second (fps) at a resolution of 320 * 240 pixels. Combined with the 120-degree lens, the angular span of two pixels on the camera is about 0.3 degrees. This allows the simulated visual system to detect image motion with spatial resolution as small as ∼0.3 retinal degrees, similar to the retinal DSGCs of other vertebrates^99^. To simulate the motion fusion effects of biological retinal processing, we added a motion-blur effect using the function provided by *Webots* simulation. This blurring was achieved by computing the weighted average of the current camera output image (weight of 0.0088) and the previous image (weight of 0.9912). The controller (i.e., the *simZFish* neural network) runs at a frequency of 1000 Hz, receiving images from the cameras as input and generating the desired angles of the servomotors of the body. For convenient observation of the internal states of *simZFish*, we designed a built-in window in the graphical interface of the simulation (**Video 1**, **Extended Data Fig. 2a**). The window can display the internal states, including neuronal activation, time, bearing (head direction), position, and action possibilities. This simulation also can show the view from the simulated cameras. To present simulated visual stimuli to the *simZFish* retina, we constructed virtual displays in *Webots 2021a*, like the bottom projected visual stimuli in our real zebrafish experiments. Each simulated visual display is 8 * 8 cm with a 2048 * 2048 pixels resolution. In principle, the choice of visual environments is limitless, including various visual naturalistic scenes (**Extended Data** Fig. 2b). To test the OMR, we programmed the displays to show black and white gratings with a spatial frequency of 1 cycle per centimeter, moving at a velocity of 10 mm/s, and a refresh rate of at least 250 Hz, like during real zebrafish experiments^6^. To simulate closed-loop experiments performed on the real zebrafish, the visual displays rotate and move to lock to the simulated zebrafish’s head orientation and position (**Video 1**).

#### simZFish neural architecture of simulated OMR neural circuitry

The neural network architecture of *simZFish1.0* is composed of three parts: the visual processing circuits (retina and pretectum), the central sensorimotor circuits (anterior hindbrain (*aHB*), nucleus of the medial longitudinal fasciculus, *nMLF*, and ventral spinal projection neurons, *vSPNs*), and the locomotor circuits of the simulated spinal cord, including central pattern generators (*CPGs*) and motor neurons (**Supplementary Methods**). The visual processing circuits are modeled using rate-coding neuron models, which compute the firing rate of a neuron based on a sum of inputs and a sigmoid transfer function. The sensory inputs to the artificial neural network in the *simZFish* simulation are light intensity values of single pixels of simulated cameras, mimicking artificial photoreceptors as in the real vertebrate retina (320 horizontal x 240 vertical pixels). To process the visual information in the artificial retina, these specific pixel values are projected to the bipolar cells (*BC*) layer. These *BCs* are inspired by the transient-OFF-bipolar cells in real vertebrate retinas that detect the variations of ambient light levels and are computed using a classic delay line model. This artificial retina drives four types of artificial direction-selective ganglion cells (*DSGCs*) with preferred directions: anterior (nasal), posterior (temporal), superior (dorsal), and inferior (ventral). These are computed from *BC* neurons with a three times lower resolution to reduce computational costs (102 horizontal x 78 vertical *DSGCs* of each preferred direction). The simulated *DSGCs* project directly to their contralateral downstream targets, the monocularly selective early pretectal neurons (*ePT*). These *ePTs* have been experimentally identified and previously modeled by us as so-called relay pretectal neurons as they inherit the functional responses from the four types of *DSGCs*^6^. Thus, on each side of the pretectum, we implement four *ePT* neurons. After initially connecting the entire retina to the *ePT* layer, which results in poor simulated OMR behavior, given the experimentally derived response constraints, we connected only *DSGCs* responsive to the lower-temporal quadrant of the visual field (*cf*., **Fig. 3**), which is experimentally supported by receptive field measurements in the live larval zebrafish^33^. The *ePT*s project to complex, binocular neurons in the late pretectum (*lPT*) ^34,100,101^. In total, we implemented eight *lPTs* in *simZFish1.0* on the left and their mirror symmetric counterparts on the right side of the brain: *oB*, *B*, *iB*, *ioB*, *iM_M_*, *M*_*M*_, *oM_L_*, *S*. In *simZFish2.0* there are more *IPT* neurons, as we divided the *oB*, *B*, *Mm*, and *S* neurons into *oB1, oB2, B1, B2, Mm1, Mm2, S1,* and *S2* neurons, respectively, where the type 1 neurons respond to forward motion, whereas the type 2 neurons respond to backward motion. The *lPT* neurons project to *nMLF* and *aHB* (see **Fig. 1e**). The activation of the *nMLF* neurons determines bout frequency, and the *aHB* neurons determine turning behavior. Both types of neurons have low-pass filters to represent delayed and smoothed responses. The low pass filter of the *aHB* neurons aids in compensating for self-generated visual motion (**Extended Data Fig. 6**). To accomplish the characteristic bout and glide behavior of larval zebrafish, we implemented a specific ‘Bout gate’ center that functions as a leaky integrator, observed in the real anterior hindbrain ^102^. In *simZFish* the ‘Bout gate’ center initializes bouts by collecting input from left and right artificial *nMLF*. When the integrated value reaches a predetermined threshold, the integrator triggers, ‘opens the gate’, for both the ‘*Behavior determinator’* center and the activation of the *CPGs*. In the meantime, the leaky integrator resets itself, preparing for the next bout. Higher incoming activity in the *nMLF*, therefore, leads to higher bout frequencies. When activated by the ‘Bout gate’ center, a ‘Behavior determinator’ center stochastically generates forward, leftward turning, or rightward turning bouts, with probabilities based on the input from both left and right *aHB* and *nMLF*. Higher values of left and right nMLF increase the probability of a forward bout; a higher left aHB and right aHB promote the probability of a left and rightward turning bout, respectively. The undulations needed for the swimming bouts are generated by a *CPG* composed of twelve coupled phase oscillators activated by the ‘*Behavior determinator’* center. The amplitudes of the oscillators behave like an exponential function dampened by a low-pass filter. This configuration produces a simulated tail-beating amplitude with a quick rise and slow decay. A bout is terminated if the amplitude decreases to less than 10% of the initial value of the exponential function. To generate some randomness on the bout distance, we added noise in the bout duration. Turning bouts are mediated through ventrally located spinal projection neurons (*vSPNs*) in the hindbrain ^30^ that activate the motor neurons on the ipsilateral side. When the ‘*Behavior determinator*’ generates a turning bout, *vSPNs* on one side are more activated than the other one, and this leads to higher motoneuron activation, and hence a higher curvature, on that side.

### Physical robot - *ZBot*

#### ZBot physical construction

To substantiate our findings from the *simZFish* simulations in realistic conditions, we designed, constructed, and tested a robot (*ZBot*) whose length (∼80 cm) and weight (2.7 kg) represent a reasonable trade-off between approximating real larval zebrafish morphological features and satisfying technical requirements to equip a moving fish-like robot with cameras, electronic boards, batteries, motors, and waterproofing (**Extended Data Fig. 9a**). To approximate larval zebrafish morphology, the *ZBot* consists of seven independent, serially connected modules (one head and six similar body segments). These modules are made of 3D-printed polylactic acid (X-Max, Qidi Tech, China). Each module is connected by a hinge joint, which is actuated by a servomotor (XM430-W350-T/R, ROBOTIS Co., Ltd., South Korea). Such configuration allows the hinge joint to rotate in the horizontal plane, like the *simZFish*. By controlling the relative rotational angles of each joint, the *ZBot* can generate a head-to-tail traveling wave of undulation along the body and replicate real zebrafish swimming maneuvers. The head module (with a dimension of about 21 cm * 8 cm * 10 cm in length * width * height) is equipped with two laterally positioned color-sensitive CMOS cameras (MU9PC-MH, XIMEA GmbH, Germany). Each CMOS sensor contains 5 megapixels, each pixel size of 1.2 um for a 5.7 x 4.28 mm active sensing area. We choose this camera for its compact size (15 x 15 x 8 mm) and available acquisition rates (ranging from 5 to 240 frames per second (fps)). To replicate the optical configurations of real zebrafish eyes, we installed each camera with a wide-angle (125 °), low-distortion lens (M27289M07S, Arducam, China). Since experimental zebrafish data show that the spatial and temporal sensitivity in motion detection is about 0.5 to 3 degrees and 10 to 20 ms, respectively, we set each camera to operate at a resolution of 320 * 240 and a framerate of 50 fps. To maintain a frame rate of 50 fps, we set a fixed exposure time of 10 ms. To make the camera adapt to a wide range of environmental ambient light levels, we activated an automatically adjustable gain function.

Attached to the head module, the *ZBot*’s body consists of six connected modules (each with a dimension of about 7 cm * 4 cm * 6 cm in length * width * height). To communicate between the central controller and the servomotors, a USB cable connects the central controller to a USB-RS485 converter (DYNAMIXEL U2D2, ROBOTIS Co., Ltd., South Korea). Then, this RS485 bus connects the USB-RS485 converter and all the servomotors in series. On the end of the tail, the *ZBot* has a flexible fin with a length of about 15 cm and a height of 12 cm along the ventral dorsal axis. The *ZBot* was equipped with a Lipo battery (11.1V 3s 2500 mAh 30C, nVision NVO1811, Neidhart SA Switzerland) in the head to supply power to the sensors, controller, and motors, lasting for about one hour at full charge. To move the center of mass to the head, like in the real zebrafish, we attached steel weights to the head. The center of mass is, therefore, located at the level of the connection between the head and first tail module. To waterproof the *ZBot*, we applied coating on both the inside and outside of the hard shell of the head. The rest of the body is wrapped in a plastic (thermoplastic polyurethane) sleeve with a thickness of about 1 mm. This soft, waterproof sleeve allows each segment to rotate about 30 degrees. Further, we attached a hard shell to each segment outside the soft sleeve to ensure that the surfaces interacting with water always maintain the same form and area. This configuration provides a waterproof, flexible, and consistent robotic morphology.

The *ZBot* head module contains a central computer (Raspberry Pi 4 Model B, Raspberry Inc., UK) that runs the *simZFish* neural network software that models the neural circuitry of the eyes and brain, as for the *simZFish*. This central controller obtains images from each camera and executes the computation. The computer is powered by a microprocessor with four computational cores and 4 GB RAM. Such a multicore controller allows for the efficient processing of incoming images and sensory computations in parallel, like biological nervous systems (**Extended Data Fig. 9b**). As for the simulation, recordings of the activation of the artificial neurons are stored for post-processing.

#### ZBot robotic experiments - head fixed in air

To test the *ZBot*’s capabilities under idealized conditions, we performed head-fixed experiments, with the *ZBot* suspended in mid-air with the robotic head directed upward (**Extended Data Fig. 9c**). To present visual motion stimuli, we mounted a 75-inch television (FW–75BZ35F, Sony cooperation group, Japan) vertically with 20 cm between *ZBot* and display, mimicking visual stimuli from below in a hanging configuration. To reduce the vibration from the tail beating, the servomotors of the *ZBot*’s tail were switched off. Visual stimuli were presented with the default brightness and contrast settings, like stimuli presented to real zebrafish during OMR experiments (**Extended Data Fig. 9d, e**). To avoid the effects of light reflection on the display, we darkened the room. The visual stimuli were presented with a spatial period of 10 cm and moving at a speed of about 10 cm/s. During each trial, visual stimuli were presented for 100 seconds, while the *ZBot*’s OMR program automatically recorded the activation of the artificial neurons by writing log files for over 120 s.

#### ZBot robotic experiments - in stationary water

To evaluate the swimming behavior of the *ZBot*, we conducted stationary water experiments in the harbor of Lake Geneva (Port du Bief, Préverenges, Canton de Vaud, Switzerland). This harbor provided a vast, protected body of calm water that allowed the robot to execute several bouts in a sequence without reaching the margins (**Extended Data Fig. 9f**, **Video 8**). Compared to space-constrained experiments in our lab swimming pool, the open lake also reduced the effects of the reflecting waves caused by *ZBot*’s tail movements. To avoid other sources of waves, we conducted the experiments in mild weather conditions with wind speeds less than 5 km/h. To test the parameters for swimming actions, we manually set the robot to perform forward and turning bouts and recorded head direction, head position, and tail-segment angles. To measure the kinematic behaviors of the *ZBot* in the lake, we used camera drones (Mavic mini or Mavic mini-2, SZ DJI Technology Co., Ltd., China). This drone hovered in mid-air, about 5 meters, and recorded videos of the swimming *ZBot*. A low-distortion camera mounted on the drone recorded videos at a frame rate of 30 fps, a resolution of 3840 x 2160 (4k), and a viewing angle of 83 degrees. To reduce the image blur of the drone video frames, we set a low shutter time (with a constant value from 1/1000 s to 1/2000 s) and a low ISO (with a constant value of 100). On the head of the *ZBot*, we used a white cross as a marker for visual tracking. To track the movement of the tail, we marked each segment with a black dot on the rotational axis of each segment. Using the open-source Tracker^103^, we extracted the robotic kinematics, including the head direction, head position, and tail-segment angles (**Extended Data Fig. 9g-i**).

#### ZBot robotic experiments - in water currents of a natural river

To test the *ZBot*’s ability to navigate in natural, moving waters, we conducted experiments in the river *Chamberonne* (Lausanne, Canton de Vaud, Switzerland). We chose this river for several reasons: its water is often relatively clear, the riverbed consists of rocks, serving as rich visual stimuli, and its relatively shallow depth (0.3 to 0.6 m) and moderate width (6 m) offer safe experimental access. Before conducting any experiments, we checked the weather records to avoid rain in the surrounding area in the past five days, as rain muddied the water, decreasing visibility. Finally, we selected this river as it flows at a reasonable speed of approximately 0.5 m/s, which is critical to evoke visual responses of the onboard OMR neural mechanism. The water flow in natural rivers varies due to many aspects, including the time of day and weather of the past days. Therefore, to ensure comparable current velocity for all trials, we conducted all the experiments within one hour on the same day. For all river experiments in **Fig. 5**, we recorded the robot’s position via a drone (Mini SE, SZ DJI Technology Co., Ltd., China). Before each trial, a drone pilot operates the drone remotely to hover at ∼12 m above the river. At this height, the drone recorded an approximately 15 * 15 m field of view at 30 fps with a resolution of 2720×1530 (∼2.7k). Each trial started with the *ZBot* software operator remotely controlling the onboard *ZBot* OMR circuit software via a laptop with a Wi-Fi connection. Once started, the *ZBot* software operator signaled the physical *ZBot* operator to release the robot and drone pilot to start the aerial video recording. For each trial, the *ZBot* was released with the same initial upstream heading direction (0 degrees). Throughout each experiment, the *ZBot* recorded artificial neural activation (e.g., *ePt*, *nMLF*, etc.) internally by writing log files and the visual input data from the onboard cameras as long as memory constraints permitted. We terminated each trial after 120 s when the robot collided with the river’s edge or swam out of the camera view. In total, we performed 29 trials with OMR, without OMR, and with the *ZBot*’s motors off. After 29 trials, the robotic battery reached the low limit of its capacity. Of the 29 trials, we only included 14 trials in the final analysis because the other 16 trials were corrupted due to collisions with the river edge (4 trials), swimming upstream out of the drone camera view (3 trials), and failure of Wi-Fi communication (6 trials). Three examples are shown in the **Video 9**.

### Zebrafish

For all live zebrafish behavioral or two-photon imaging experiments, we used 6-9 days-old zebrafish larvae. These were collected as fertilized eggs as offspring from adult breeder zebrafish. All zebrafish were maintained on a 14/10 hour light/ dark cycle at 28.5 °C. Embryos were raised in small groups of about 30 fish in pH-buffered, filtered E3 solution (5 mM NaCl, 0.17 mM KCl, 0.33 mM CaCl2, 0.33 mM MgSO4). From 4 days post fertilization (dpf) onwards, we fed larvae with paramecia. We performed all imaging and behavioral experiments with 5-7 dpf zebrafish. At these stages, sex cannot be determined. For imaging, we used homozygous *Tg(elavl3:H2B-GCaMP6s)* in the Casper background^104^, lacking dark melanocytes across the skin but with normal retinal pigmentation. Before imaging, we screened zebrafish for bright fluorescence to achieve the best calcium signals. The *Tg(elavl3:H2B-GCaMP6s)*^105^ was a generous gift from Dr. Misha Ahrens. All experiments were approved by Duke University School of Medicine’s standing committee of the animal care and use program (IACUC).

#### Behavioral recordings of optomotor responses in live, larval zebrafish

To record the behavior of freely swimming, real larval zebrafish, we adapted closed-loop experimental routines to test responses to visual motion patterns, similar to our previous experiments ^6^. Using custom wired 850 nm illuminator panels (CMVision, CM-IR130-850NM) paired with an 830 nm long-pass filter (Edmund Optics, SCHOTT RG-830), we achieved infrared bottom illumination of the behavior arena, invisible to the animals. Zebrafish behavior was recorded from above at 163 Hz using an infrared-sensitive, high-speed, CMOS camera (FLIR, GS3-U3-41C6NIR-C: 4.1 MP, 90 FPS, CMOSIS CMV4000-3E12, NIR) with a 35mm 1” lens (Edmund Optics, 35mm Focal Length Lens, 1” Sensor Format, #63-247). Visual motion stimuli were projected onto a diffusing screen after reflection from a cold mirror (Edmund optics) using a P300 Pico Projector (AAXA technologies). Zebrafish x, y, and orientation were tracked using the open-source framework *Stytra*^106^, and this information was rerouted into a custom 3.7 *Python pandastim* visual stimulus rendering software. Briefly, using a self-triggering trial structure, each trial started with concentrically moving rings to orient zebrafish to the center of the dish. Detecting the zebrafish’s tracked position in the center triggered the start of a trial. For each trial, black and white gratings of different orientations were locked relative to the fish body axis, with a black bar (0.8 cm) presented directly underneath the live zebrafish to isolate visual stimulation of each eye.

These stimuli were locked to the position and angle of the fish to maintain a closed-loop configuration, ensuring the same direction of the visual motion stimulus despite the zebrafish’s movements. Starting with 3 s of non-moving, stationary gratings to control for orientation-related behaviors, orientation-locked gratings began to move at 10 mm/s (or remained stationary throughout the trial, depending on the stimulus identity). Visual motion stimuli were continuously presented for up to 30 s or until the fish aborted the trial early by leaving the active area. Only fish that completed at least 10 repetitions in less than 3 hours were included in further analysis. Extracted position and orientation data were further analyzed to extract individual bouts and compared to *simZFish* behavior.

#### Behavioral analysis of live zebrafish swim kinematics

All behavioral analyses were performed using custom *Python* or *Matlab* scripts (*Mathworks*, USA) similar to those described previously^6^. Briefly, we extracted individual locomotion bouts by detecting the x and y trajectories multiplied by traces of orientation changes. The heading angle change for each bout was calculated as orientation directly before (2 ms) and after (10 ms) a detected swim bout. Across stimuli and trials, this resulted in a data frame containing information about each bout’s angle change and distance swimming in response to visual stimulus. Average bout frequency was computed as the total number of detected swim bouts over the total stimulus time across zebrafish and plotted as bout frequency histograms. Bouts with a negative value indicated rightward (or clockwise) and positive values indicated leftward (or counterclockwise) heading angle change. All behavioral error bars represent the standard error of the mean (SEM) across zebrafish for previously published orthogonal set N = 38, for parallel moving stimuli, N = 16. For histograms, the shaded area is SEM across fish for angle bout type (**Fig. 1e, Extended Data Fig. 7**). Assuming the behavioral data were normally distributed, we used non-parametric statistics for all hypothesis testing.

#### Two-photon calcium imaging in live zebrafish

To record neural activity in live zebrafish, we performed volumetric calcium imaging with a custom two-photon laser-scanning microscope, using a pulsed Ti-sapphire laser tuned to 950 nm (*InsightX3*, Spectra Physics, USA) as an excitation source. Live, transgenic zebrafish expressing the calcium sensor GCamP6s, *Tg(elavl3:H2B-GCaMP6s)*, were head-fixed using 2 % weight/volume low melting point agarose (*Sigma Aldrich*) in embryo medium. The agarose was removed around the tail to record tail movements and monitor health during the experiment using infrared light and a camera (Grasshopper3-NIR, FLIR). We recorded fluorescent images with a 20X Olympus objective, with an imaging power of ∼10 mW at the specimen. For all functional, volumetric recordings, we imaged 300 x 300 pixels, spanning an area of approximately 300 µm x 300 µm at acquisition rates of 0.85 – 0.9 Hz. We recorded 8 planes per zebrafish, with each plane recorded 5 µm apart, spanning a 40 µm volume surrounding the pretectum and anterior hindbrain of each zebrafish, which can be identified via anatomical landmarks. As in *simZFish* and freely swimming zebrafish experiments, visual stimuli were presented with a black bar directly underneath the fish. Each stimulus lasted 15 s in total, beginning with a 10 s stationary period during which the orientation of the grating changed, followed by a 5 s moving stimulus (**Extended Data Fig. 8a**, **Video 7**). This allowed us to differentiate responses to orientation change and characterize each neuron’s direction and ocular preference. All visual stimuli were generated by custom Python 3.10 software (*pandastim*). Using a 60 Hz, P300 AAXA Pico Projector, only the red LED was activated, allowing for simultaneous visual stimulation and detection of green fluorescence. These red stimuli, which evoke strong OMR responses in freely moving zebrafish, were created as independently moving textures for each eye (stationary and moving phases, grating spatial frequency, direction, orientation, and speed), resulting in 16 unique motion stimuli and a control stimulus (**Supplementary Methods**).

#### Two-photon Ca^2+^ imaging data processing and analysis

Following data acquisition, uncompressed image stacks were movement corrected using *CaImAn*, an open-source calcium imaging processing library, and its implementation of the NoRMCorre algorithm, which calculates and aligns motion vectors with subpixel resolution^107,108^. The *Suite2p* software package was used for source identification and signal extraction^109^. To assess stimulus responsivity, we extracted the z-scored change in activity over time and smoothed this with a box filter. This metric was averaged over five stimulus presentations to generate the average response of each neuron. Neurons were defined as stimulus-responsive if their correlation to an idealized response trace was greater than 0.65. The eight overrepresented neuronal subtypes (*oB, B, iB, ioB, iMm, Mm, oMI, S*) were extracted using a barcoding method as described previously^6^. An unbiased hierarchal clustering was generated within these subtypes using the SciPy software package, enabling unbiased analysis of overrepresented response types within each subtype. This clustering within the eight symmetric functional subtypes, only four (*oB*, *B*, *Mm*, and *S)* neurons showed over at least ∼25% of neurons subclustered into clear subtypes, showing no significant response over 1.8 STD above baseline to either forward or backward stimuli. The remainder showed no significant or mixed preferences for forward or backward motion.

#### Comparison of simZFish and live zebrafish

Zebrafish and *simZFish* behavioral data were preprocessed to extract the specifics (heading angle, time, distance moved, x, y, coordinates) about each locomotion bout per visual stimulus for subsequent comparative analysis. For qualitative comparisons (**Fig. 2d, Extended Data Fig. 7**), all bouts were plotted across all trials for each stimulus for a representative zebrafish that completed at least ten trials and those of a continuous 2000 s per stimulus experiment. While repeated experiments with *simZFish* show low variability (**Extended Data Fig. 4b**), zebrafish behavior can vary substantially. Therefore, for quantitative comparisons for absolute bout histograms and bout probabilities, we used continuous 2000 s behavioral recordings for *simZFish* and zebrafish average behavioral data.

### Statistical Analysis

All values are reported as mean ± standard deviation unless otherwise stated. In behavioral bar graphs, error bars correspond to the standard error of the mean S.E.M. unless otherwise stated.

## Data Availability Statement

Raw two-photon microscopy data, as well as all the processed data (source-extracted fluorescence time series) for all experiments, are publicly available at https://dandiarchive.org/dandiset/001076. All other data and associated analysis code will be available upon publication on a public Gitlab repository hosted by Duke University.

## CONTACT FOR REAGENTS AND RESOURCE SHARING

Further information and requests for resources and reagents regarding experimental zebrafish data and robotic simulations should be directed to Eva Naumann and Xiangxiao Liu.

## Code Availability Statement

The complete code to run the *simZFish* within the Webots simulator is available at https://ponyo.epfl.ch/proj/zebrafish/simzfish. This repository also contains a detailed user guide (’simZFish_User_Instructions.pdf’) to guide the installation and execution of the simulations. All software for *simZFish* and zebrafish behavior and neural activity calcium imaging analysis is available at https://github.com/Naumann-Lab. All code used was created using Python or approaches in MATLAB 2023b and open-source code extensions. The visual stimuli presented to zebrafish were designed in custom Python software, PANDASTIM, running Panda3D (Disney). Instructions for running analysis code and reproducing published results are available via the code repository README (rendered by GitHub as a web page) and Jupyter Notebooks, which include code to produce specific figures.

## Acknowledgments

We thank Drs. Timothy Dunn and Henry Greenside for helpful comments and discussions; Misha Ahrens for the *Tg(elavl3:H2B-GCaMP6s)* transgenic fish lines; and Florian Engert for support with pilot behavioral experiments. We thank Duke School of Medicine, J. Burris, K. Olivera, and L. Frauen for zebrafish husbandry. We thank Alessandro Crespi for his technical support in constructing the robot. This work is supported by grants from the ERC (Synergy grant 951477, Salamandra) to A.I. Research reported in this publication was supported by the BRAIN initiative of the National Institutes of Health under Award Number RF1NS128895-01. The content is solely the authors’ responsibility and does not necessarily represent the official views of the National Institutes of Health. M. D. L. and E.A.N. also were supported by the Whitehall and Alfred P. Sloan Foundation.

## Author contributions

X.L., A.I., and E.A.N. conceived of this project. X.L. and L.Z. constructed and tested the neuromechanical simulations. M.D. L. performed and analyzed all live zebrafish calcium imaging experiments, and M.D.L., K.F. collected all live zebrafish behavioral data. M.D.L., K.F., X.L., and E.A.N. analyzed and compared robotic simulation data. X.L. F.L. and L.Z. constructed, tested, evaluated, and experimented with the physical robot. X.L., M.D.L., A.I., and E.A.N. wrote the manuscript and contributed to figure preparation with input from all authors. A.I. and E.A.N. supervised the project.

## Ethics Declaration

None

---

**Extended Data Figure 1.**
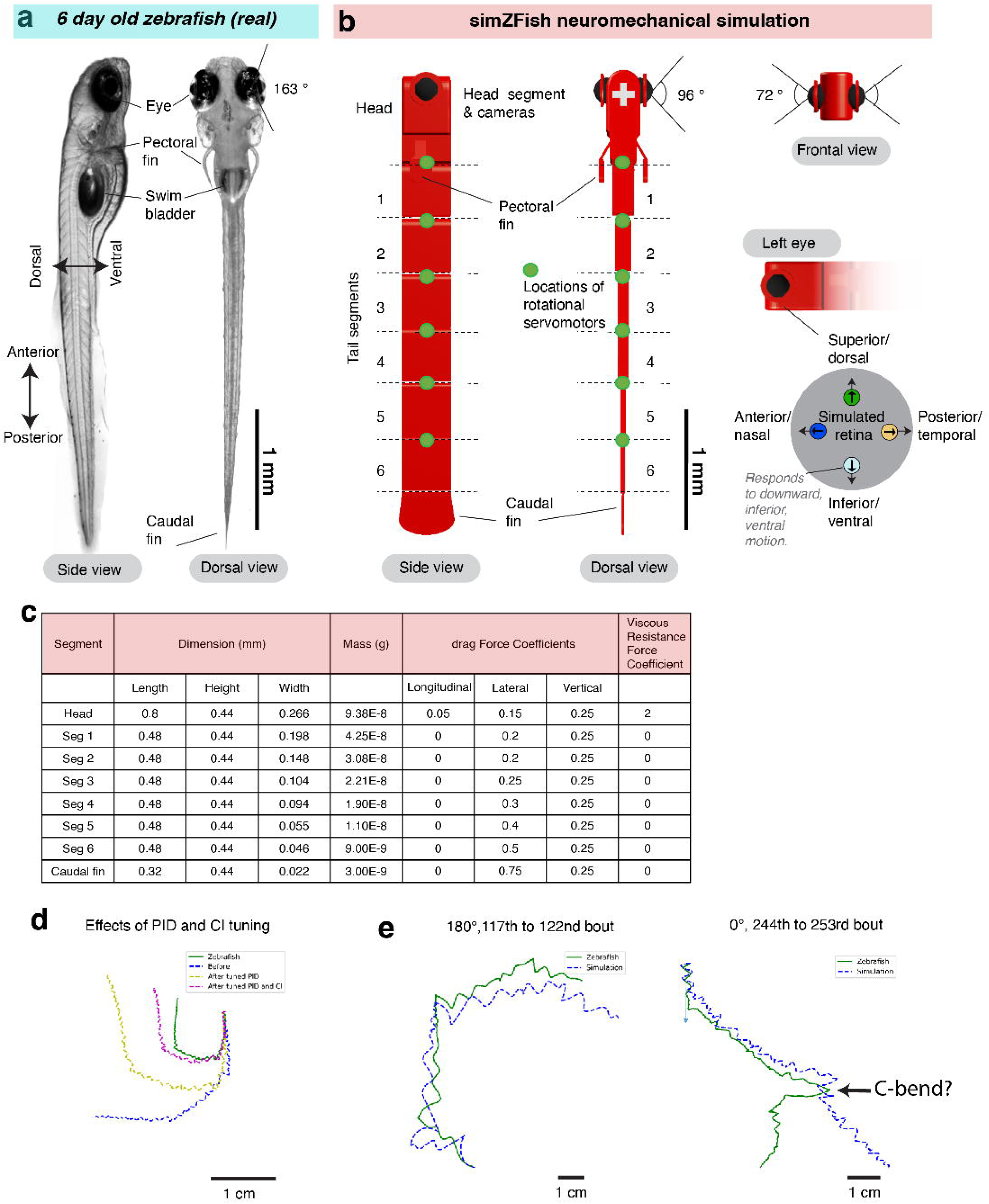
Larval zebrafish inspired *simZFish* body design.

**Extended Data Figure 2.**
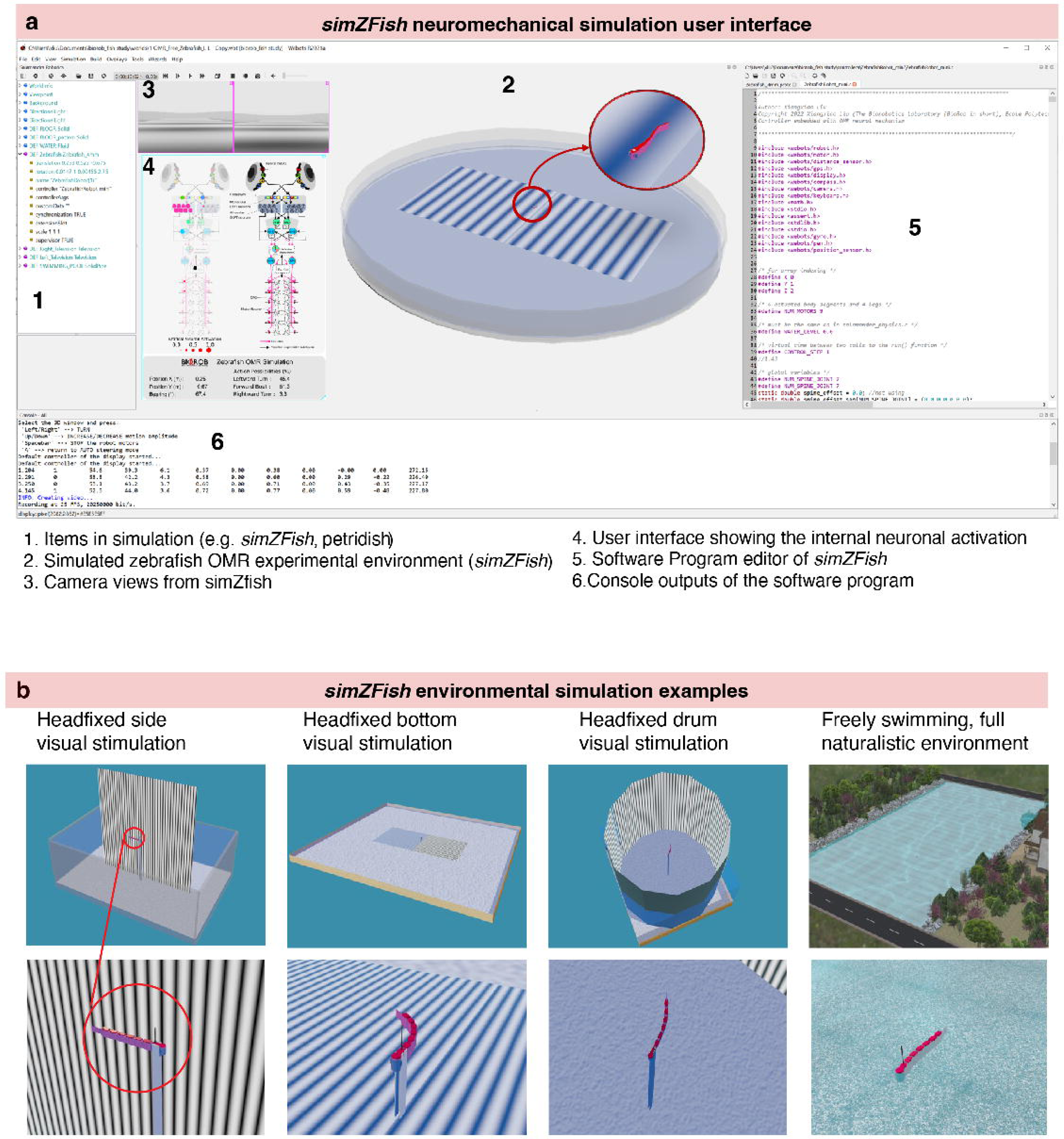
*simZFish* interface enables limitless simulated experimental designs.

**Extended Data Figure 3.**
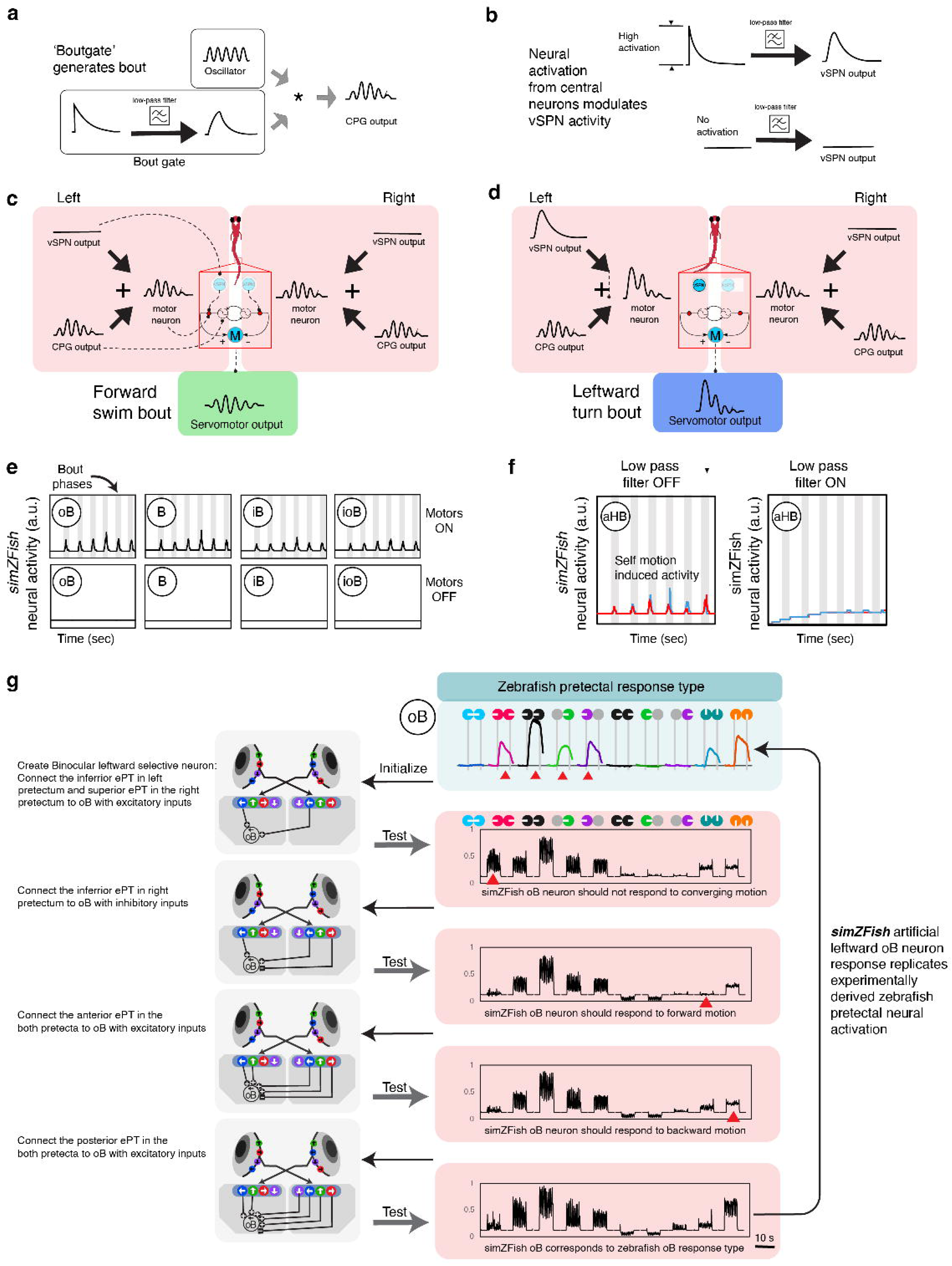
*simZFish* bout generation and neural connectivity tuning.

**Extended Data Figure 4.**
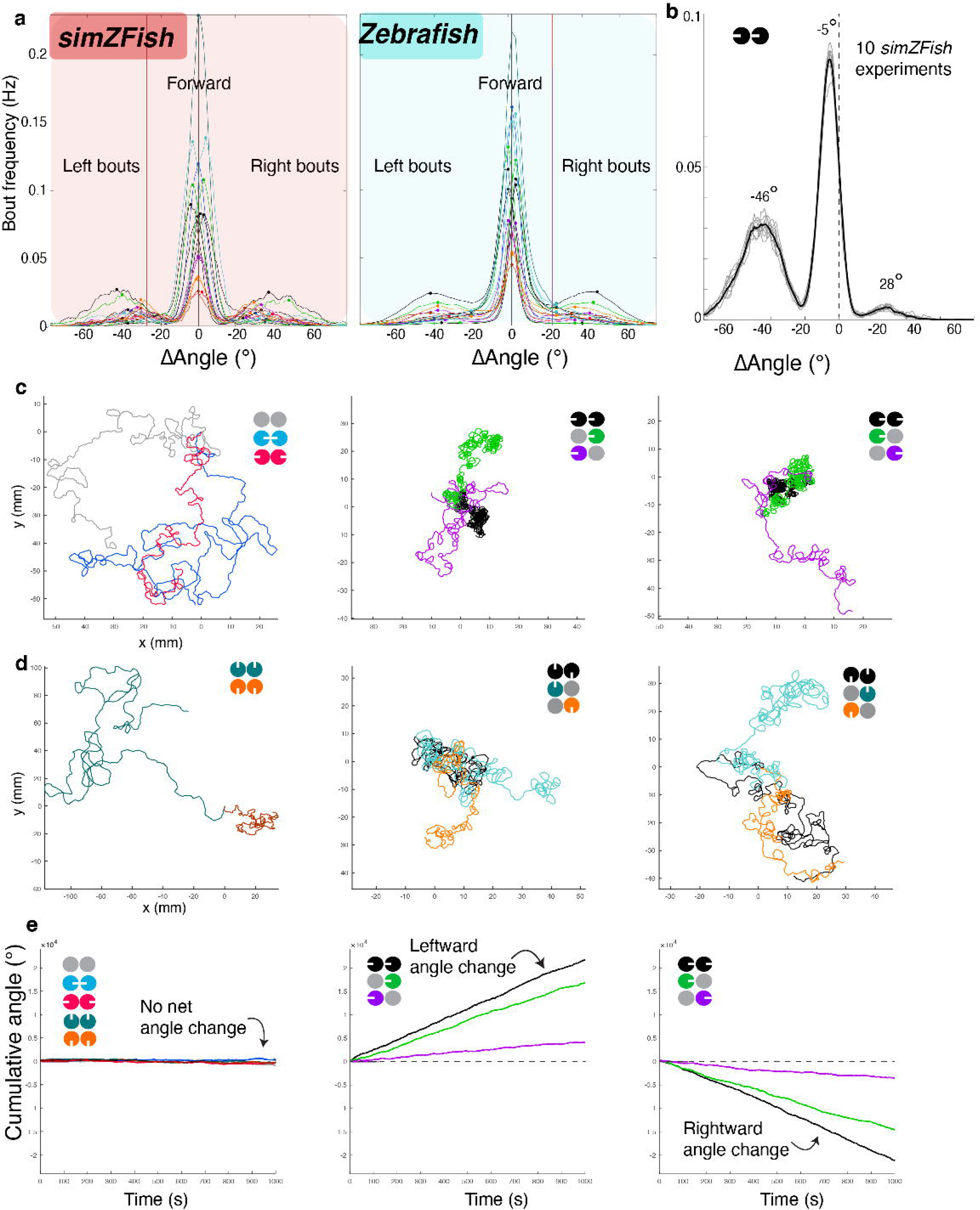
*simZFish* replicates zebrafish OMR behaviors.

**Extended Data Figure 5.**
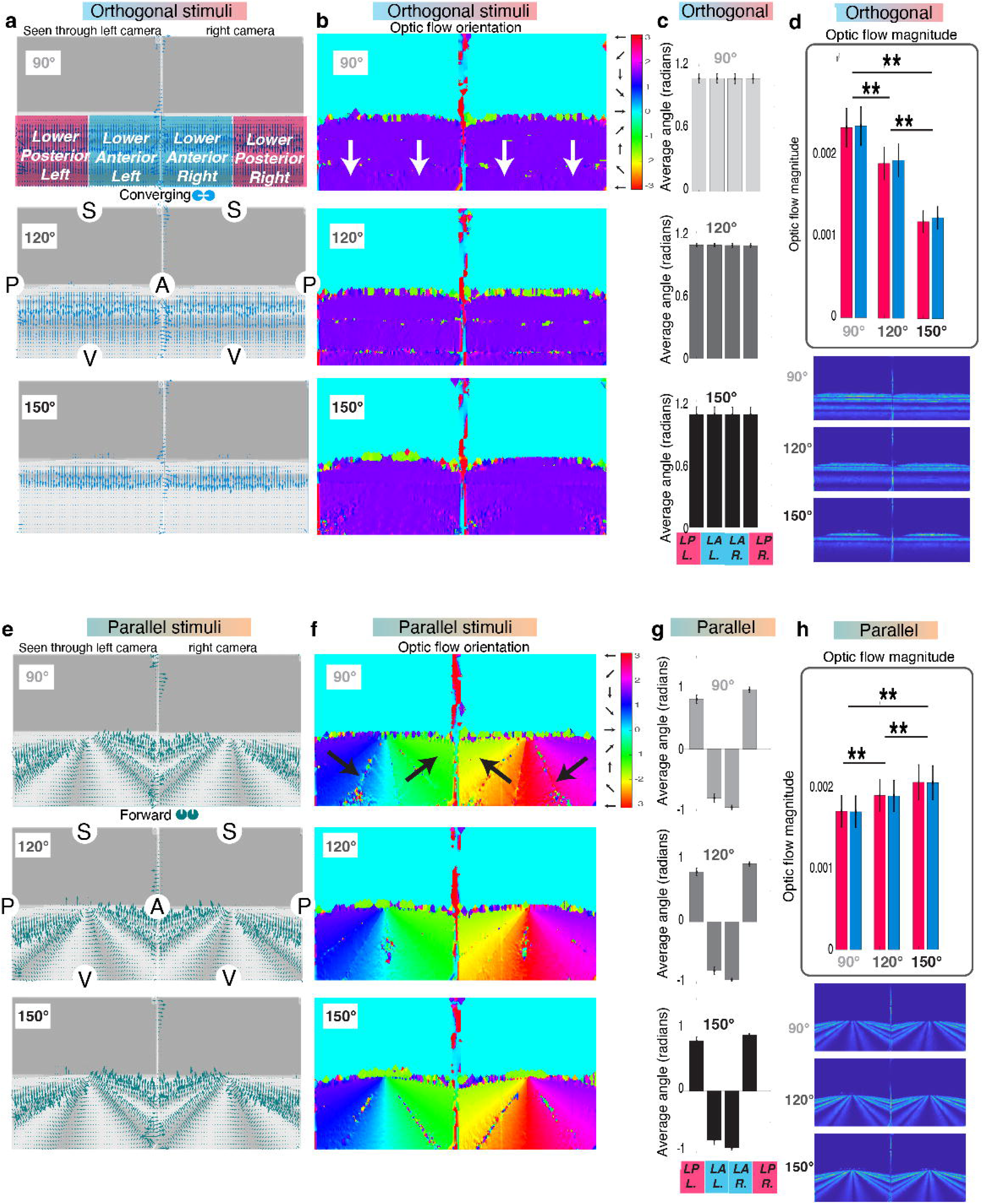
Optic flow analysis of visual stimuli seen through the eyes of *simZFish*.

**Extended Data Figure 6.**
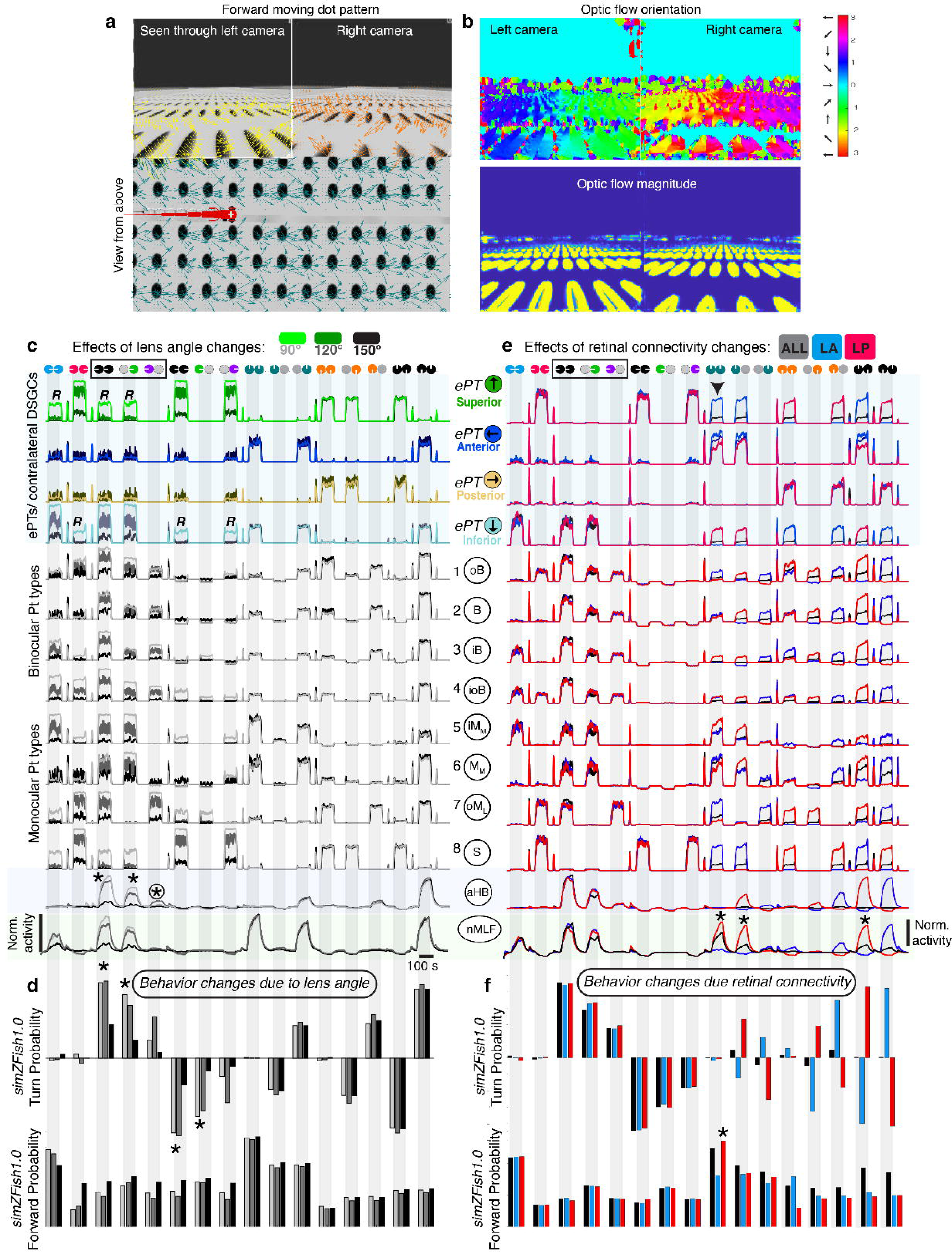
Embodiment effects on neural processing and behavior in *simZFish 1.0*.

**Extended Data Figure 7.**
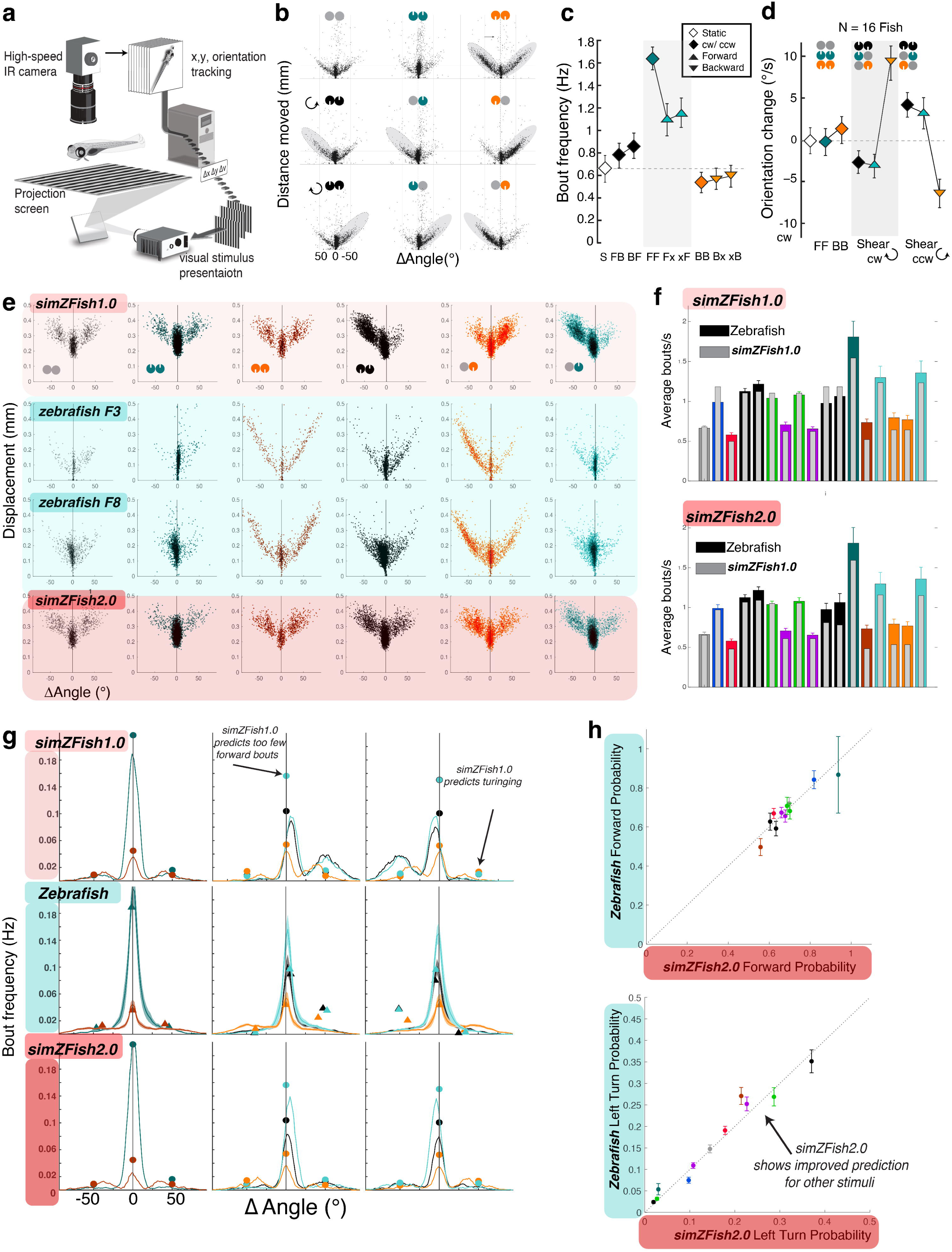
Zebrafish behavioral measurements suggest *simZFish 1.0* network updates.

**Extended Data Figure 8.**
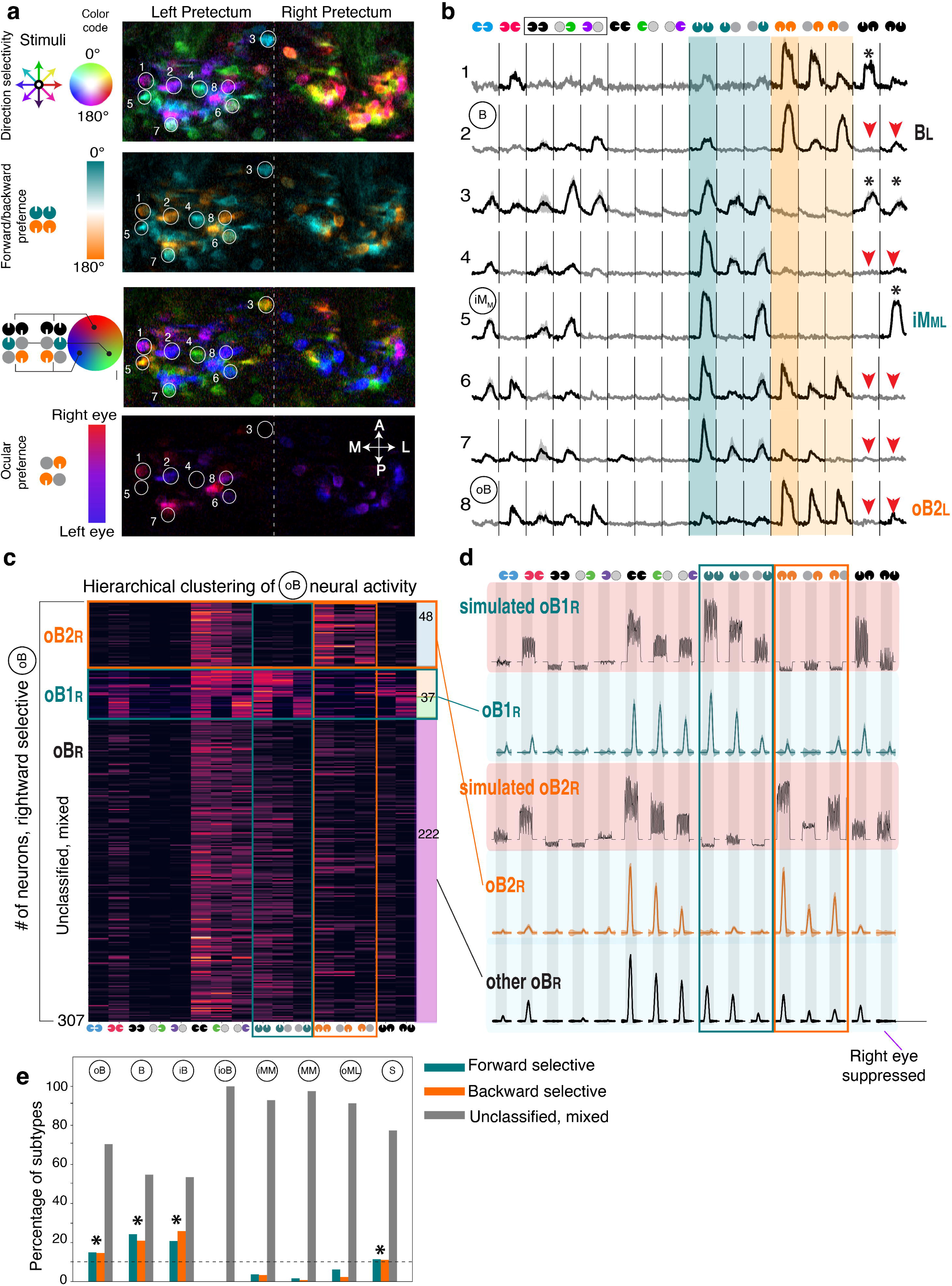
Zebrafish neural activity measurements suggest *simZFish 1.0* network updates.

**Extended Data Figure 9.**
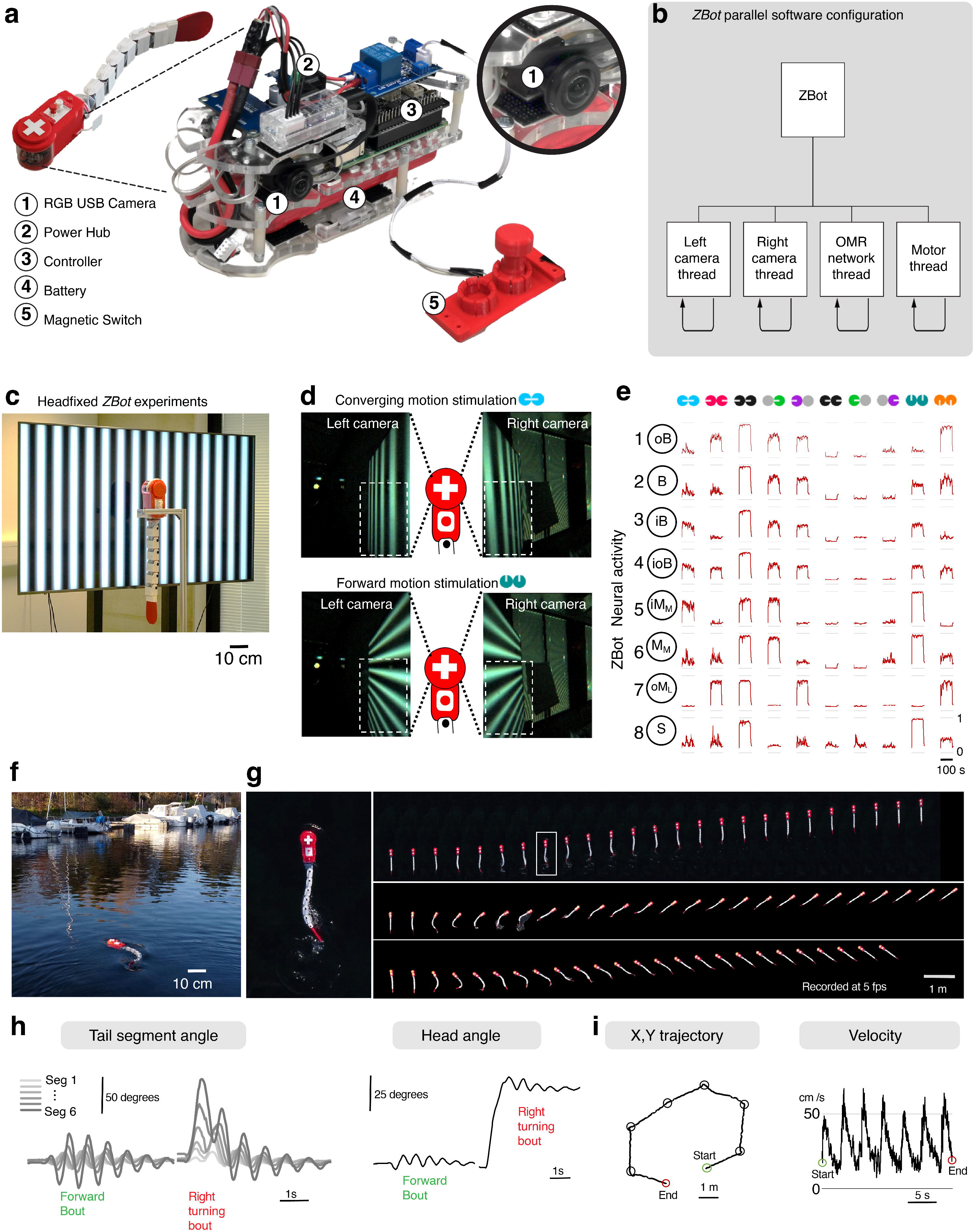
Physical, zebrafish inspired robot, ZBot replicates neural activation and OMR behaviors.

**Extended Data Figure 10.**
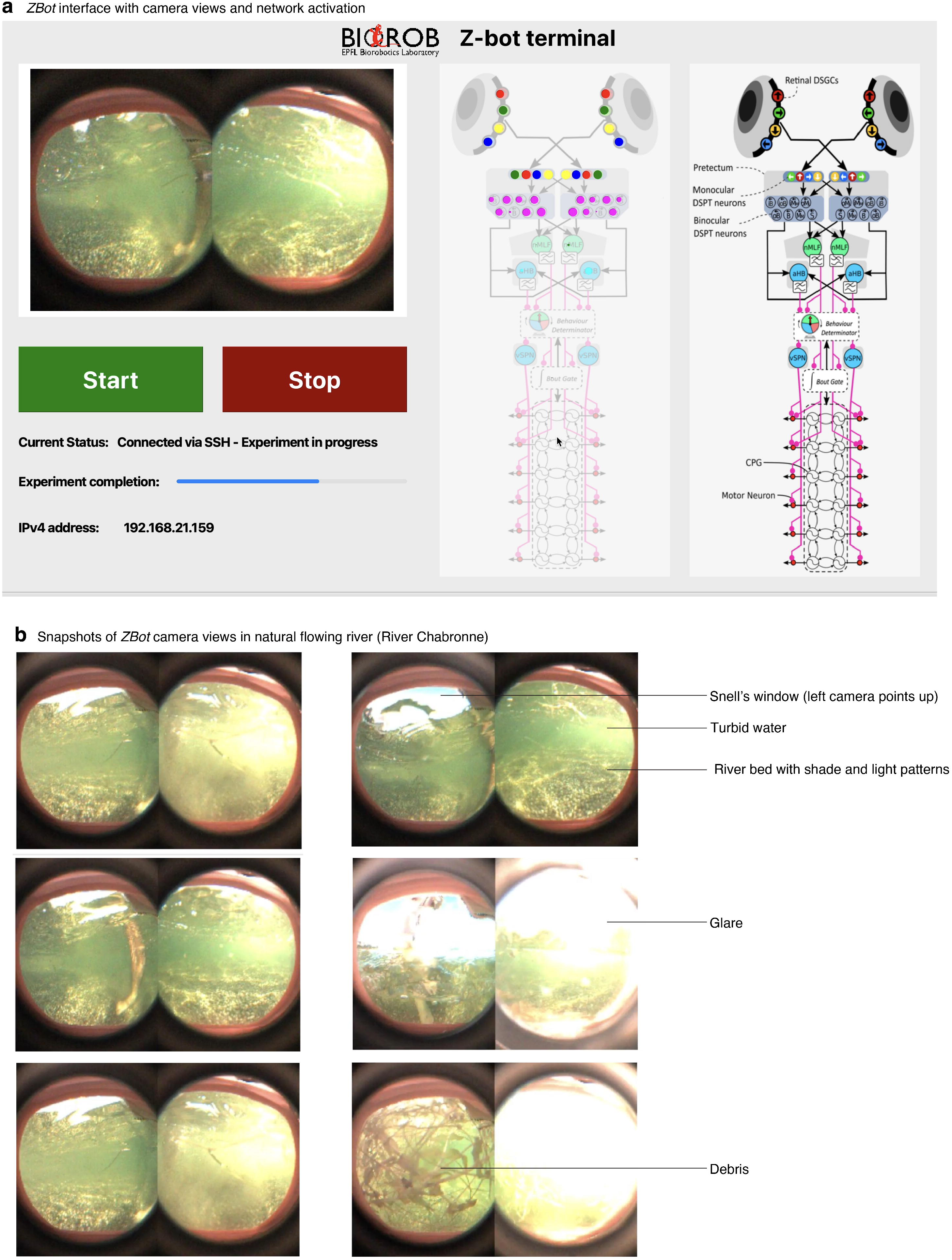
Zebrafish inspired ZBot performs OMR behaviors in naturalistic flowing river.

