## Supplementary Information for "Artificial Embodied Circuits Uncover Neural Architectures of Vertebrate Visuomotor Behaviors"

Affiliations :
[1] Biorobotics Laboratory, EPFL - École Polytechnique Fédérale de Lausanne, Lausanne 1015, Switzerland

[2] Duke School of Medicine, Department of Neurobiology, Durham, NC 27710, United States

[3] Institute for Systems and Robotics, Instituto Superior Técnico, 1049-001 Lisbon, Portugal

*These authors contributed equally to this work.
† These authors jointly supervised this work.

Corresponding Authors:



**Supplementary Information contents:**

1. Supplementary Text
2. Supplementary Methods
3. Supplementary Tables 1 to 3
4. Extended Data Figs. 1 to 10
5. Supplementary References
6. Supplementary Video 1 to 9 Legends

### 1. Supplementary Text:

#### A. Investigating brain scale visuomotor transformations in embodied artificial agents:

*Neuromechanical simulations as investigative tools*

To holistically understand the neural principles underlying visuomotor transformations *in vivo*, one must investigate the entire neuromechanical system[1]. This includes all visual sensors, neural circuit elements, the body, and the environment. Analyzing these subsystems in isolation is insufficient, as visual perception drives motor behaviors, which, in turn, affects perception, with complex non-linear interactions resulting in brain-wide multi-dimensional activity patterns[2,3]. In the past, simulations and robots were used to verify whether all subcomponents are sufficiently understood to replicate specific animal behaviors, e.g., the transition from swimming to walking in a salamander[4], the tradeoffs between stability and maneuverability of unseen forces during locomotion[5], how bees perform path integration[6], and to discover redundant central and peripheral mechanisms for undulatory swimming generation[7]. Leveraging the significant advances in neuromechanical simulations of walking[8,9], flying[10], or swimming[11], our study aimed to provide detailed insights into the neural representations of complex, varying sensory environments and the role of embodiment in neural functionality and behavior.

*Enhancing biorealism and neural circuit exploration in simZFish: pathways for future development*

Balancing complexity with computational requirements, we implemented and tested biologically realistic visuomotor neural circuits in simulations (*simZFish*) and (*ZBot*). Both are approximations of the real larval zebrafish and could be improved in several ways.

First, the *simZFish* body could be improved with additional degrees of freedom to allow 3D swimming (instead of swimming only under the surface of the water) and the addition of muscle models. Moreover, *simZFish'* and *ZBot's* position maintenance is not perfect, especially in currents with higher velocity. In future versions, speed control could be improved with more powerful motors, longer bouts, or bouts of higher oscillation frequencies or amplitudes.

Second, for the upstream visuomotor circuits, we can investigate how many different neural networks based on the described specific pretectal neurons can lead to similar behavior. For example, when considering the behavioral similarity in responding to the eleven types of visual stimulation used in previous studies (**Fig. 2**), both *simZFish*1.0 and *simZFish* 2.0 lead to similar behavior. However, when considering the similarity in response to novel visual stimulations, one observes that *simZFish*2.0 leads to different behavior compared to model *simZFish1.0*, better matching our new biological behavioral data (**Fig. 4c**). Therefore, our *simZFish* predictions motivated both new hypothesis-driven behavioral and neural recordings that uncovered novel functional neural subtypes contributing to visuomotor processing. Demonstrating the utility of our model, failure modes in *simZFish1.0* suggested new biological experiments, which led to updated network connectivity and behavior in *simZFish2.0*, both presenting an improved model and a roadmap for how neuromechanical simulations can be used as a neuroscience research tool. Incorporating this biological complexity enhanced *simZFish2.0*'s accuracy. This bidirectional interchange between computational modeling and empirical testing represents a valuable strategy for reverse-engineering biological systems. Through them, we identified new subtypes of pretectal neurons and revealed computations previously ignored (**Fig. 4f**). To systematically study variations of these networks, one potential avenue would be to instantiate parameters of the networks using machine learning, such as backpropagation[12] for supervised learning when recordings of target neurons exist, or proximal policy optimization [13] for reinforcement learning of target behaviors. Such approaches could be combined with statistics on multiple networks generated with different initial sets of random parameters to investigate how many networks can generate a particular behavior. Another avenue would be to take an automated science approach [14–16] in which new visual stimuli are generated autonomously and tested on both the real animal and the simulation in a manner that is optimal in terms of information theory (i.e., testing new visual stimuli that maximize the chances of identifying which simulated neural circuit matches the real one).

Third, our model of the zebrafish nervous system could be improved and extended. For instance, our currently very abstract reproductions of the locomotor circuits (*nMLF*, *vSPN*, and *CPGs*) could be replaced with more realistic neural networks, and the simplified rate-coded neuron model used for the visual circuits could be replaced with more realistic spiking neural models. Furthermore, these neural circuits are far from comprehensive, and networks could eventually include models of the cerebellum for motor adaptation [17], and integrate crucial sensory input from other modalities, such as mechanosensation of the lateral line organ [18], other visual input for prey capture predator avoidance [19,20]. Extending *simZFish* with other realistic multimodal brain circuitry and musculoskeletal actuation will be important for understanding neural control of the flexible body morphology. Recent efforts in mapping complete synaptic connectomes in the zebrafish [21] and the fruitfly (*Drosophila melanogaster)* and efforts [22] to model the neural functions within these networks will be important to increase the biorealism of these stimulations. Future studies could then investigate how different sensory modalities, both for exteroception (vision, olfaction, lateral-line organ, and tactile sensation [18,23]) and for proprioception (e.g., intraspinal stretch[24] and cerebral fluid sensors[25]). One could explore multiple sensory modalities to investigate the relative gains of different sensorimotor loops. Our open-source tools could also investigate different visually guided behaviors (e.g., predator avoidance, obstacle avoidance, and feeding behavior), and reverse engineer the respective gains of different visuomotor and other sensorimotor loops. Iteratively performing simulation, robot, and animal experiments has great potential to gradually improve our understanding of the neuromechanical principles underlying animal adaptive behavior.

**B. Optomotor responses and rheotaxis:**

Rheotaxis is a behavior influenced by multiple sensory modalities, including vision, the lateral line, tactile sensation, the vestibular system, and proprioception[26,27]. The potential role of vision in rheotaxis is not new, but we show that the visuomotor transformations underlying the zebrafish OMR circuit can perform rheotaxis without any other sensory input (**Fig. 2g**, **5d**, **Video 4**). Beyond visually driven OMR, other studies have indicated that the lateral line organ, which measures variations in water flow and pressure, could play an important role in rheotaxis in larval zebrafish [26,28]. Remarkably, studies in larval zebrafish[26] and blind cavefish[29] with disrupted lateral lines have shown that rheotaxis can be performed without vision cues and nor lateral lines, suggesting that even other sensory cues, such as tactile stimuli, can also be used for aligning against the flow. OMR neural circuits utilized by *simZFish* and *ZBot* are sufficient to perform rheotaxis without other sensory modalities if visual information is available. In the simulated and real river, these circuits allowed the *simZFish* and *ZBot* to reorient themselves against the water flow and help maintain position despite the highly variable visual input in the natural river (**Fig. 5**). The OMR is likely most useful when the water flow is highly turbulent and irregular, rendering the mechanosensory information from the lateral line organ and tactile sensors unreliable[18,28]. Conversely, information from the lateral line and tactile sensors (and other senses) are useful when visual input is absent (in the dark) or corrupted (in high turbidity water)[29].

**C. Effects of embodiment on neural architecture:**

Our *simZFish* experiments agree that it may be optimal to possess wide-angle lenses with *DSGCs* attending to the lower visual fields (**Fig. 3b**, **Video 5**), while for other species with different visual environments and sensory requirements, such as rodents[30], may lead to other asymmetries[31]. Our analyses (**Extended Data Fig. 5**, **6**) show that the perspective geometry of viewing bottom-projected visual patterns dictates the connectivity of neural architecture with optimal wiring. For instance, motion patterns parallel to the body axis, such as tail-to-head (forward) motion, as would be commonly experienced by fish, even with more natural stimuli than sinusoidal gratings (**Extended Data Fig. 6a**). This perspective distortion causes a rotational optic flow pattern which results in opposing optic flow in anterior and posterior visual fields (**Fig. 3e**, **Video 6**). While the eye should be a priori perceptually symmetric, this opposing optic flow information to the lower anterior (*LA*) and posterior (*LP*) quadrants presents a computational dilemma, considering evolutionary energetic constraints may require the fewest neuronal connections[32]. Instead of utilizing information across the entire eye, the zebrafish retina may have solved this by mainly using information from the lower posterior quadrant of the visual field, aligning with biological findings that pretectal neurons' preference for the stimuli in the lower posterior visual field[33]. While it may at one point in evolution have been plausible to construct a functioning neural network relying only on the lower anterior visual field, our simulations predict that medial and backward stimuli would co-activate the same pretectal neurons, which strongly disagrees with empirical data[33–35]. With retinal functional asymmetries well appreciated[31,36], we argue that it is, in fact, the physical nature (i.e., rotational optic flow) of the incoming sensory information that shaped these asymmetries in receptive field properties.

To test these ideas, we used *simZFish* to manipulate the connectivity of *DSGCs* to pretectal neurons in various visual receptive field locations (**Fig. 3g**). The results show that, indeed, *LP* connectivity results in strong co-activation of *anterior* and *inferior-selective DSGCs*, whereas *LA* connectivity leads to coactivation of anterior and superior *DSGCs.* Thus, when starting with a contralateral design (all retinal projections decussate at the midline, as in *simZFish*) present in most bilaterally symmetric nervous systems[37], *LP* receptive field connectivity would pair incoming (medial) with forward (tail to head) motion. Medial motion, which also drives strong locomotion[3], activates *inferior-selective DSGCs*, linking with the strong stabilizing response of the tail-to-head (forward) motion stimulus, which, with *LP* connectivity, activates *inferior* and *anterior-selective DSGCs*. This relevance is reflected in the fact that many neurons in real zebrafish show a combinatorial preference for medial and forward stimuli (both drive higher bout frequency) or lateral and backward stimuli (both reduce bout frequency)[3]. Indeed, these motion preference pairings might be a corresponding consequence of the early emphasis on contralateral neural pathways, as medial motion may be privileged due to the ethological relevance in other situations, such as approaching objects.

### 2. Supplementary Materials and Methods:

#### A. *simZFish*: Neuromechanical simulations of larval zebrafish

##### *Modeling the simulated simZFish body:*

The model of the body of *simZFish* is implemented using the Webots 2021a simulator (Cyberbotics Ltd, Switzerland). We chose *Webots* over other simulation environments for robotics, e.g., *mujoco* or *pybullet,* as it offers a complete, open source, platform with extensive sample world libraries, detailed API documentation in C, C++, Java, Python, and MATLAB or ROS combined with a user friendly graphical user interface (GUI)[38] (**Video 1**). Additionally, Webots offers extensive camera simulation options essential for the simulation of vision-based behaviors embedded in the *simZFish.* We approximated the real larval zebrafish body dimensions with a length of 4 mm, a height of 0.044 mm, and a weight of 0.0938 mg, based on average physical measurements of larval zebrafish described in Zhao et al. 2020[39] (**Extended Data Fig. 1a-c**). Unlike the slightly denser-than-water real larval zebrafish[40], the density of the *simZFish* is slightly lower than the water itself (91.5 % of the density of water). Therefore, the *simZFish* body floats just below the water surface, with the cameras fully immersed in the simulated water. We chose *simZFish* to float just below the surface as this simplifies the control of 3D movements and leads to swimming in the horizontal plane, resembling the real, live zebrafish experiments performed in shallow water. Notably, both *simZFish* and *ZBot* swim at the surface, with the density of the *ZBot* also slightly less than the density of the water. The center of mass is located slightly lower than the geometric center of each segment, improving its passive balance in water. Nonetheless, real 6-day-old zebrafish can perform skillful vertical underwater swim maneuvers[40]. Real zebrafish appear to actively maintain horizontal posture, engaging both tail and pectoral fin movements to overcome their slightly negative buoyancy and an observed tendency to sink[41]. Furthermore, another benefit of the *ZBot* floating just below the water surface permits wireless communication. The simulated *simZFish* body has seven segments: the first head segment, which contains the simulated cameras, and six body segments (the last body segment is connected to a flexible caudal tail fin). This approximation of the body of real six-day-old larval zebrafish was chosen over more sophisticated geometries to reduce computational cost. Therefore, to increase the simulation speed, we simplified the shape of each segment as a cuboid, with a smaller width from head to tail (**Extended Data Fig. 1b**). The center of mass of the *simZFish* body is located near the head segment, like real larval zebrafish. Each segment is connected by a hinge joint under the actuation of a simulated servomotor (six in total), composed of a PID controller that performs joint angle position control and tracks the desired joint angles provided by the swimming controller.

***Hydrodynamics***

Each segment of the *simZFish* body is subject to both static and dynamic forces from the surrounding water. The static forces include Archimedes' thrust, whereas the dynamic forces include viscous resistance forces and inertial drag forces. Laval zebrafish swim at Reynolds numbers between 10 at very young ages (2 to 5 days post fertilization) and 1000, corresponding to intermediate regimes where viscous and inertial forces play important roles[42]. To compute these physical forces, we used Webots' built-in fluid dynamics model. Webots uses drag models that are based on the linear and rotational velocities of each segment. Viscous forces and torques are proportional to the viscosity of the water and the segment's linear and angular velocities, respectively. Inertial drag forces are proportional to the fluid density, the apparent area, a (direction-dependent) drag coefficient, and the square of the linear velocity. For equations and more details, see Webots' documentation (https://cyberbotics.com/doc/reference/immersionproperties). Such a fluid dynamics model represents a first-order approximation of real fluid dynamics and provides a reasonable estimate of real swimming regarding forward speed and turning while being computationally light. More detailed simulations of larval zebrafish swimming exist based on solving the full Navier-Stokes equations on a grid (see, for instance, Zhao et al., 2012[39]), but they are computationally much more costly.

To generate realistic zebrafish behavior, the main coefficients to tune were the unitless coefficients for inertial drag forces ('dragForceCoefficients', **Extended Data Fig. 1c**). These coefficients are empirically known for simple objects (e.g., spheres, cylinders, or cubes) with specific orientations in water flow but need to be tuned for articulated bodies like *simZFish*. The longitudinal drag value was set to 0.1 for the head segment and to zero for the following segments, following the measures performed in McHenry and Lauder, 2005[41], and a similar difference between the head and the rest of the body in a simulation of lamprey swimming[43]. The lateral coefficients are set to small values for the rostral segments and larger ones for the caudal segments. This increase in drag coefficients is motivated by the fact that the rostral segments of zebrafish larvae are like cylinders. In contrast, the caudal segments are like plates, with higher drag coefficients than cylinders. The vertical coefficients are set to a constant value of 0.25 (but these do not affect swimming much since *simZFish* swims just below the water surface, with limited vertical displacements).

In larval zebrafish swimming, viscosity plays an important role in hydrodynamics, including viscosity during zebrafish gliding[41]. To modulate the viscosity coefficient, the *Webots* simulator provides a parameter called *viscousResistanceForceCoefficient* that can be set from 0 to infinity. In the simulation, we configured the *viscousResistanceForceCoefficient* to 2.0 for the head segment and 0.0 for the other segments (**Extended Data Fig. 1c**). We tried other combinations of these viscous coefficients, for instance all equal along the body, or with a gradient from head to tail, and these led respectively to worse or similar match to animal data (data not shown). Therefore, we kept the configuration where only the head segment has a non-zero *viscousResistanceForceCoefficient.* Finally, we tuned the hydrodynamics parameters (inertial drag force coefficients) to match kinematic recordings of zebrafish swimming (**Extended Data Fig. 1d**).

To tune and test the neuromechanical simulation, we compared *simZFish* with real, high-speed (> 200 HZ) zebrafish data from Marques et al., 2018. We extracted the point joint angles from the zebrafish recordings and replayed them using the simulated servomotors of *simZFish* to replicate real zebrafish body deformations over time (**Extended Data Fig. 1e**). Replaying recorded kinematics from real zebrafish, we demonstrate that *simZFish* produces forward and routine turning bouts that closely match recorded displacements of the real animal (**Video 2**). Comparing the *simZFish* bout distance to zebrafish kinematic recordings, we show that *simZFish* matches the forward displacements per bout within 14.4 %. Comparing the turning angle to the zebrafish, the *simZFish* simulation matches the turning angle per bout by 9.8 %. **Video 2** illustrates how the *simZFish* replicates a long series of swimming bouts from previously published[44] high-speed recordings by Marques et al., 2018. While this test reveals slight error accumulation over time, *simZFish* represents a good first-order approximation of real larval zebrafish swimming. *simZFish* can perform most routine turns and swims observed during the optomotor response (OMR). Nonetheless, in the future, further refinement might be necessary to replicate the entire natural behavioral repertoire, such as fast escapes (C bends, **Extended Data Fig. 1e**, *right*) and prey-hunting maneuvers.

***Dimensional units and integration time steps***

Six-day old zebrafish are tiny, larval fishes (~ 4 mm) that swim with a high tail-beating frequency (e.g., around 30 Hz during a routine forward swim bout). By default, *Webots* uses the International System of Units. Consequently, the tiny size of larval zebrafish leads to very small values for some of its physical properties (e.g., a total volume is ~$10^{-10}$ $m^3$). Such excessively small values can cause numerical errors in the simulation calculations. To cope with this, we redefined the unit of length from the meter (m) to the centimeter (cm) and adjusted all physical quantities (e.g., the gravity constant) accordingly (**Supplementary Table 1**). This unit transformation is identical from a physics point of view but significantly improved the numerical stability of the *simZFish* simulation by avoiding rounding errors of minimal values of state variables. It also allowed us to simulate a reasonable integration time step size (0.05 ms) using *Webots* default integration method (first order semi-implicit integrator). Note that the activity states of the *simZFish* neural network of the visuomotor neural control circuits are updated at a lower rate (every 1 ms).

***simZFish neural network architecture***

The neural network architecture of *simZFish* is composed of three parts: (1) the visual motion processing circuits (retina and pretectum); (2) the central sensorimotor circuits (hindbrain, nucleus of the medial longitudinal fasciculus *nMLF*, and ventral spinal projection neurons *vSPNs*); (3) and the locomotor circuits of the simulated spinal cord, including central pattern generators (*CPGs*) and motor neurons (**Fig. 1**, **Extended Data Fig. 1b**). Here, we first describe the implementation of *simZFish1.0* (**Fig. 1, 2**) and later explain the changes for the biologically constrained, updated *simZFish2.0*. Unless explicitly specified, the equations provided here are valid for both models.

All *simZFish* neural circuits are modeled using rate-coding neuron models, which compute the firing rate of a neuron based on the sum of inputs and a sigmoid transfer function. For example, equation (1) shows the mathematical expression of the firing rate $B$ of a neuron that receives the inputs $A_i(i = 1, 2, 3 \ldots)$ from other neurons:

$$B = \frac{1}{1 + e^{-\omega(\sum w_i A_i - b)}} \quad (1)$$

where $A_i$ and $w_i$ are the firing rates from the descending neurons and their weight, respectively. $\omega$ and $b$ are parameters of the sigmoid transfer function. For all artificial neurons in the *simZFish* networks, we manually set these two parameters through trial and error to achieve a similar response of the artificial neurons compared to the responses of corresponding neurons recorded in fish (see below, **Supplementary Table 2** for *simZFish1.0* and **Supplementary Table 3** for *simZFish2.0*).

**Photoreceptors to bipolar cells (*BCs*) Layer**

The simulated sensory inputs to *simZFish* artificial neural network are light intensity values of single pixels of simulated cameras, mimicking artificial photoreceptors of the real vertebrate retina that provide information to generate direction-selective neurons[45]. In *simZFish,* the pixel values are generated from simulated images streamed from two bilaterally installed red, green, and blue (RGB) cameras. The resolution of each image is

320 (horizontal) * 240 (vertical) pixels, a standard resolution for commercial cameras. While the simulation runs, these images are read at a frame rate of 1000 frames per second (fps). To mimic the real zebrafish eye's optical properties, these two cameras are placed laterally in the head segment and equipped with no-distortion lenses (**Fig. 1a**, **Extended Data Fig. 1a, b**). Thus, the sensory input layer can be described as $P_t(i,j)$ to represent each (artificial photoreceptor) pixel's value (by calculating the average value of the red and green channel) at time *t* with the location coordinates $(i,j)$ on the image. For example, $P_t(i,j)$ indicates a pixel from the current frame. $P_{t-25}(i,j)$ represents the same pixel value from a frame 25 ms ago. The location coordinate of $(1,1)$ indicates the most left, upper pixel on any given image. Therefore, each pixel value can be described via its specific location on the image as well as its value at a specific time point, representing a rectangular simulated retina. For instance, $P_t(50,50)$ represents the value of a pixel at time 0 ms (current frame) at a position of $(50,50)$ on the visual receptive field of the *simZFish* retina.

To process the visual information in the artificial *simZFish* retina, these specific pixel values are projected to the bipolar cells (BC) layer. In the *simZFish*, we used 320 (horizontal) * 240 (vertical) bipolar cells, directly matching the number of incoming pixel values. Like the definition of $P_t(i,j)$, each BC's activity is represented as $BC_t(i,j)$, with $(i,j)$ representing the location coordinates and $t$ representing timepoint in the simulation. These BCs are inspired by the transient-OFF-bipolar cells in real vertebrate retinas that detect the falling of ambient light levels[46,47]. Thus, the activation of each BC at time $t$ and location in the neural network on the left eye is defined as:

$$BC_t(i,j) = \frac{1}{1 + e^{-3(P_{t-\Delta t}(i,j) - P_t(i,j) - 1)}} \tag{2}$$

where $P_{t-\Delta t}(i,j)$ is the value of pixel $(i,j)$ at $(t - \Delta t)$ ms. $P_t(i,j)$ is the value of pixel $(i,j)$ at $t$. We empirically set the temporal gap $\Delta t$ as 25 ms. Furthermore, $i \in (1,240)$ and $j \in (1,320)$.

**Bipolar cells (*BCs*) to direction-selective ganglion cells (*DSGCs*) layer**

As mammals, larval zebrafish possess *DSGCs*[48–50] and other RGCs, which can be classified by their molecular profile and morphology within the retina[51,52]. *DSGCs* can be classified as ON, OFF, and ON-OFF, depending on their preference for light increments or decrements, and further subdivided into their direction preferences to visual stimuli moving in one of the 4 cardinal directions or 3 directions for *ON-DSGCs*[53]. While recent analysis explains the lack of the fourth direction possibly as a thresholding phenomenon, current zebrafish data point toward three obvious preferred directions in retinal terminals in AF5/6[34,48]. As the zebrafish OKR is not driven by OFF ganglion cells[54] and overwhelming evidence points to *ON-DSGCs* as driver of the OKR in mammals[55], it is possible that the zebrafish OMR is also mainly driven by *ON-DSGCs*. Nonetheless, as the OMR can also be evoked local dark light transitions[47], for computational simplicity, we implemented four direction-selective types of *OFF-DSGCs*, designed to respond to anterior (A), posterior (P), superior (S), and inferior (I) motion in each eye's visual field (**Extended Data Fig. 1b**).

Thus, to enable *simZFish* to compute direction selectivity in each simulated eye, we implemented 102 (horizontal) * 78 (vertical) direction-selective ganglion cells (*DSGCs*), which receive location-specific input from the *BCs*. The resolution of *DSGCs* (102*78) is approximately three times lower than the resolution of *BCs*. We implemented this reduction by skipping 2 out of 3 BC neurons to reduce the computational cost. For example, an anterior selective DSGC of the left eye ($DSGC_{LA}$) responds to motion moving from temporal to nasal, or a posterior selective ($DSGC_{LP}$) responds to motion moving the opposite direction[56]. The neural response of an anterior selective $DSGC_{LA}$ is calculated as:

$$DSGC_{LAt}(i,j) = \frac{1}{1 + e^{-100(BC_t(3i,3j) - BC_{t-\Delta t}(3i,3j+1) - 0.3)}} \tag{3}$$

where, $LAt$ indicates left eye (L), anterior motion selective (A), at time $t$. $BC_t(3i,3j)$ is the output of a bipolar cell located at $(3i,3j)$ at time $t$, hence leading to a three times lower resolution of *DSGC* neurons compared to BC neurons. $BC_{t-\Delta t}(3i,3j+1)$ is the value of bipolar cells located at $(3i,3j+1)$ at time $t-\Delta t\ ms$. We set the temporal gap $\Delta t$ as 25 ms, matching experimentally measured temporal asymmetries between excitation/inhibition that are suggested as underlying mechanism for generating direction selectivity[57]. The distribution of bipolar cells across the visual field can be described as for each $BC_t(3i,3j)$ located at the temporal side of the associated $BC_{t-25}(3i,3j+1)$. Furthermore, $i\ \in (1,78)$ and $j\ \in (1,102)$. Similarly, the posterior (P), superior (S), and inferior (I) *DSGCs* in the left simulated eye are calculated as:

$$\mathrm{DSGC_{LPt}(i,j)} = \frac{1}{1+\mathrm{e}^{-100(\mathrm{BC_t(3i,3j)}-\mathrm{BC}_{\mathrm{t}-\Delta t}(3\mathrm{i},3\mathrm{j}-1)-0.3)}}$$
$$\mathrm{DSGC_{LSt}(i,j)} = \frac{1}{1+\mathrm{e}^{-100(\mathrm{BC_t(3i,3j)}-\mathrm{BC}_{\mathrm{t}-\Delta t}(3\mathrm{i}-1,3\mathrm{j})-0.3)}} \qquad (4)$$
$$\mathrm{DSGC_{LIt}(i,j)} = \frac{1}{1+\mathrm{e}^{-100(\mathrm{BC_t(3i,3j)}-\mathrm{BC}_{\mathrm{t}-\Delta t}(3\mathrm{i}+1,3\mathrm{j})-0.3)}}$$

**Direction selective ganglion cells (*DSGCs*) to early PT neurons (*ePT*) layer**

In the zebrafish, all retinal ganglion cells, including *DSGCs,* exclusively project contralaterally to ten arborization fields (AF)[37,51,58]. AF5/6 contains both direction-selective ON and ON-OFF tuned synaptic terminals[34], which is likely the equivalent of the retinal projection of ON-DSGCs to the medial terminal nucleus (MTN) of the AOS in mammals[59,60]. We previously showed that unilateral ablation of AF5/6 is sufficient to mimic purely monocular OMR behaviors[61]. Thus, the simulated *simZFish DSGCs* project directly to their contralateral downstream targets, the monocularly selective early pretectal neurons. These early, monocularly responsive pretectal neurons (*ePT*) have been experimentally observed and previously modeled by us as 'relay pretectal neurons', as they inherit their functional responses from the four types of DSGCs[62]. Thus, on each side of the pretectum, we implement four *ePT* neurons. For instance, on the left side, we implemented anterior ($ePT_{LA}$), posterior ($ePT_{LP}$), superior ($ePT_{LS}$), and inferior ($ePT_{LI}$). After initially connecting the entire retina to the *ePT* layer, which resulted in poor simulated OMR behavior (no increased swimming to forward motion), we connected only *DSGCs* responsive to the lower-temporal quadrant of the visual field (**Fig. 3d-g**). Detailed, receptive field measurements in the live larval zebrafish experimentally support this, as the pretectal neurons strongly prefer motion stimuli in the lower, temporal visual field, and in turn, stimuli in those visual field locations strongly drive behavior[63]. Our computational results show that OMR behaviors in *simZFish* are best driven by the lower temporal visual field, given the network constraints that require 1) all DSGCs project to the contralateral pretectum and 2) simulated neurons are tuned to replicate real neuronal response types that we recorded previously. Thus, one may argue that the connectivity in real zebrafish is an evolutionary driven phenotype to constructing a network that best utilizes the available information about the visual scene, with the fewest neurons or least connections, to minimize energy costs.

Thus, perspective distortion effects of real eyes viewing the floor, as well as in our simulation, lead to a "rotational optic flow" effect, with the stripes of the moving gratings appearing to move upwards in the lower anterior quadrant and downwards in the lower posterior quadrant of the visual field. Notably, the upper visual field is unaffected by any of our bottom projected stimuli as, in the idealized simulation with perfectly clear water, quasi-infinite distance to the arena border, and no landmarks, no optic flow can be detected. Interestingly, Wang et al. 2018, previously found that the OMR can be evoked most strongly by stimulating the lower temporal (posterior) visual field, and Pt neurons' receptive fields concentrate in that area. Thus, our simulated experiments independently predicted the connectivity and receptive field, as a fully connected retina leads to poor OMR. Therefore, all *simZFish* simulations only contain specific DSGCs to pretectum connectivity, as described below.

As pretectal neural recordings of motion stimuli demonstrate delayed and smoothed response dynamics, we added a lower-pass filter on its output with a smoothing parameter $\beta$. For example, the temporal response of the left, anterior early pretectal neuron ($ePT_{LA}$) is calculated as:

$$ePT_{LAt} = \beta ePT_{LA(t-T)} + (1-\beta)\frac{1}{1+e^{-0.1(\sum_{i,j}^{48<i<78,51<j<102} DSGC_{RAt}(i,j)-60)}} \quad (5)$$

The equation above calculates the pretectal neuron $ePT_{LAt}$, with $L$ representing the left pretectum, $A$ representing anterior selective, and t indicating the timepoint. The set LP includes all pixels such that "$48 \leq i \leq 78, 51 \leq j \leq 102$", and defines the selected rows and columns of the array of DSGCs that provide input to this specific $ePT_{LAt}$. These values correspond to the lower-temporal visual field of the simulated retina, which we have found to be most effective for proper OMR behavior generation (see **Fig. 3a**). Specifically, we chose the row value 48 (instead of 39) since such a setting allows the *simZFish* to only see visual display, that is, no information above the horizon is visible to the network. To slightly smooth the neuronal activation induced by self-motion, we used two levels of values for the low-pass filter $\beta$. We empirically set $\beta = 0.998$ and $\beta = 0.99$ when the *simZFish* is bouting and gliding, respectively. We set a higher $\beta$ during the bout phase because the tail-beating induces stronger self-motion and might influence visual perception (a higher $\beta$ value results in a stronger smoothing effect). $T$ is the time step size for the updates of the neural architecture, which we have set to $T = 1\ ms$, i.e., 20 times larger than the 0.05ms time step of the mechanical simulation, which requires a finer time scale for numerical stability.

Similarly, the other *ePT* neurons in the left pretectum are calculated as follows:

$$\begin{aligned}
ePT_{LPt} &= \beta ePT_{LP(t-T)} + (1-\beta)\frac{1}{1+e^{-0.1(\sum_{i,j}^{48<i<78,51<j<102} DSGC_{RPt}(i,j)-60)}} \\
ePT_{LSt} &= \beta ePT_{LS(t-T)} + (1-\beta)\frac{1}{1+e^{-0.068(\sum_{i,j}^{48<i<78,51<j<102} DSGC_{RSt}(i,j)-90)}} \quad (6) \\
ePT_{LIt} &= \beta ePT_{LI(t-T)} + (1-\beta)\frac{1}{1+e^{-0.068(\sum_{i,j}^{48<i<78,51<j<102} DSGC_{RIt}(i,j)-90)}}
\end{aligned}$$

**Early PT neurons (*ePT*) to late PT neurons (*lPT*) layer**

The population of monocular, direction- selective, retinorecipient neurons in the early pretectum (*ePT*) project to the more complex, binocular neurons in the late pretectum (*lPT*, **Fig. 1D**, **Extended Data Fig. 3a**), as supported by anatomical evidence[35,64,65] and our previous modeling results[61]. Therefore, in the *simZFish* these *ePT* neurons connect to *lPT* neurons to achieve the eight complex bilaterally symmetric response types (together sixteen) to bottom-projected motion stimuli that we identified in our previous work[61]. These eight bilaterally symmetric response types occurred with elevated frequency, representing over 66% of the pretectal population in real zebrafish, with four responding binocularly (activated by both medial and lateral stimuli) and others "monocular" classes activated to at least one monocular stimulus. We labelled these neurons to indicate their specific response profiles. As neurons on the left side of the brain generally respond to leftward motion, these neurons are named for their additional responsiveness to outward (o) or inward (i) moving stimuli and whether they respond to leftward motion in both eyes (B for binocular), or just the contralateral eye (M for monocular). For instance, the binocular neuron, *ioB,* exhibits responses to conflicting inward (***i**oB*) and outward (*i**o**B*) motion, responding to both conflicting, orthogonal stimuli. The monocular classes were distinguished both by patterns of excitation in response to monocular medial or lateral stimuli and by inhibition in response to conflicting motion. For example, response class $iM_M$ was a monocular ($i\boldsymbol{M}_M$) neuron, responsive to medial motion ($iM_{\boldsymbol{M}}$), and activated by inward motion ($\boldsymbol{i}M_M$). In total, we implemented these experimentally observed eight *lPTs* in *simZFish1.0* on the left and their mirror symmetric counterparts on the right side of the brain:$oB, B, iB, ioB, iM_M, M_M, oM_L, S$ (more neuron types have been modeled in *simZFish2.0*, see below). The responses of the *lPT* neurons are computed by the following equations:

$$x_{jt} = \frac{1}{1+e^{-\omega(\sum_{ij} w_{ij} ePT_{jit} - \mathrm{b})}} \quad (7)$$

$$S_{Lt} = \frac{1}{1+e^{-4(\min(0.8ePT_{RSt}, ePT_{LIt}) + 0.75ePT_{RAt} + 0.75ePT_{LAt} + 0.25ePT_{LPt} + 0.25ePT_{RPt} - 0.6)}} \quad (8)$$

In equation (7), $x$ indicates an *lPT* neuron ($x \in \{oB, B, iB, ioB, iM_M, M_M, oM_L\}$), $j$ can either be left or right, $t$ indicates the current instant, $w_{ij}$ is the weight associated with the ePT neuron of type $i$ ($i \in \{S, A, I, P\}$) in the $j$ side. Finally, $\omega$ and $b$ are the parameters of the sigmoid transfer function. Equation (8) computes the response of the $S$ neuron in the left pretectum at instant $t$ (an analogous formula is used to compute the activation of the $S$ neuron in the right PT), where $\min(0.8ePT_{RSt},\ ePT_{LIt})$ indicates the minimal value of $0.8ePT_{RSt}$ and $ePT_{LIt}$. The equations for the *S* type neurons ($S_{Lt}$ in the equation above) are different from the other neurons because an *S* type neuron only activates when both eyes are stimulated, which is one of the overrepresented response types found in the real zebrafish.

**Late pretectal neurons (*lPT*) to nucleus of the medial longitudinal fasciculus (*nMLF*) neurons**
As microscopic tracings of processes using photoactivatable green fluorescent proteins and correlational calcium imaging activation in our data suggest, the complex neurons in the later pretectum drive spinal projecting neurons in the *nMLF*[61]. In the *simZFish*, we represent these ~20 *nMLF* neurons on each left and right side of the midbrain as two simplified *nMLF* neurons (**Fig. 1a**, **e**). As the activation of the *nMLF* neurons is thought to control swim vigor and bout frequency[66], stronger activation of these *nMLF* neurons results in more bouts in the *simZFish* simulation. As our previous neural modeling suggested[61], all *lPT* neurons may project to the *nMLF* neurons. Since *nMLF* neurons exhibit delayed and smoothed responses[61], we again added a low-pass filter on the output of each nMLF neuron, with $\beta = 0.9995$. Thus, the *nMLF* neuron on the left brain is calculated as:

$$nMLF_{LT} = \beta nMLF_{L(t-T)} + (1-\beta)\frac{1}{1+e^{-6(<\theta_L,(-0.2,0.16,0.25,-0.05,0.6,0.4,-0.8,0.2)>-0.45)}} \qquad (8)$$

where $\theta_L = (oB_L, B_L, iB_L, ioB_L, iM_{ML}, M_{ML}, oM_{ML}, S_L)$ is the vector of activations of the *lPT* neurons, and $<.,.>$ denotes the inner product between two vectors.

**Late PT neurons (*lPT*) to anterior hindbrain neurons (*aHB*)**
To determine turning behaviors, we and others[64] previously hypothesized that some translation-selective late pretectal neurons project to the motor command neurons in the anterior hindbrain (*aHb*) to drive turning behaviors. Therefore, in *simZFish*, we represent *aHB* neurons as left-right symmetric single command centers on each side of the brain. To achieve appropriate turning behavior, each artificial *aHB* neuron receives inputs from *lPT* neurons from both sides of the brain. To compensate for self-generated visual motion (**Extended Data Fig. 3f**, **g**), we empirically implemented at a low pass filter with $\beta = 0.9995$, calculating the left *aHB* using the equation (9):

$$\begin{gathered} aHB_{Lt} = \beta aHB_{L(t-T)} \\ +(1-\beta)\frac{1}{1+e^{-3.85(0.575oB_L + 0.533B_L + 0.333M_{ML} + 0.392S_L - 0.333oB_R - 0.333B_R - 0.21M_{MR} - 0.58S_R - 0.76)}} \end{gathered} \qquad (9)$$

**Medial longitudinal fasciculus (*nMLF*) and late PT neurons (*lPT*) to '*Bout gate*' center**
To accomplish a characteristic bout locomotion event, a swimming burst followed by a glide phase, behavior observed in real, live zebrafish, we implemented a specific neuron to function as a '*Bout gate*' center. This '*Bout gate*' center functions as a (leaky) integrator, observed in the anterior zebrafish hindbrain for evidence accumulation[67]. In *simZFish,* the '*Bout gate*' center initializes bouts by collecting input from the left and right artificial *nMLF* neurons. When the integrated value reaches a predetermined threshold, the integrator triggers, '*opens the gate'*, for both the '*Behavior determinator*' center and the activation of the *CPGs*. In the meantime, the integrator resets itself and prepares for the next bout by resetting to baseline activity. Higher incoming activity in the *nMLF*, therefore, leads to higher bout frequencies. The mathematical expression of the leaky integrator of the '*Bout gate*' center is shown in equation (10), with $a$ representing the rate of leaking.

$$I_t = aI_{t-T} + (nMLF_{Lt} + nMLF_{Rt}) + b_t \qquad (10)$$

$$I_t = 0 \text{ if } I_t > threshold\ or\ during\ a\ bout$$

For simplicity, we set $a$ to a value of 1 in the *simZFish* simulation (meaning this is a pure rather than leaky integrator). After reaching the threshold ($I_t > 436$), the *simZFish* simulation generates a swim bout while the *'Bout gate'* center is reset. In the equation, $b_t$ is a positive baseline to ensure spontaneous bouts even in the absence of visual stimuli. To add randomness to the bout frequency over time, we draw $b_t$ from a folded zero-mean normal distribution with a standard deviation of 0.6 ($b_t = 0.6|N(0,1)|$). This leads to an average bout frequency of about 0.4 Hz.

**Medial longitudinal fasciculus (*nMLF*) and anterior hindbrain (aHB) to '*Behavior determinator*'**

After receiving the signal from the *'Bout gate'*, an artificial neuron termed the '*Behavior determinator'* chooses an action from three options: forward bout, leftward turning bout, and rightward turning bout. The '*Behavior determinator'* calculates a probability for these actions based on the input from *aHB* and *nMLF* as:

$$P_l = \frac{aHB_l + b_l}{aHB_l + aHB_r + nMLF_l + nMLF_r + b_l + b_r + b_f}$$

$$P_r = \frac{aHB_r + b_r}{aHB_l + aHB_r + nMLF_l + nMLF_r + b_l + b_r + b_f} \tag{11}$$

$$P_f = \frac{nMLF_l + nMLF_r + b_f}{aHB_l + aHB_r + nMLF_l + nMLF_r + b_l + b_r + b_f}$$

where, $b_l$, $b_r$, and $b_f$ are positive values to ensure the baselines of turning/forward-bout rates. We set them as 0.066, 0.066, and 0.2, respectively. For instance, higher values of *nMLF* neurons increase the probability of a forward bout; a higher left *aHB* and right *aHB* promote the probability of a leftward turning bout and rightward turning bout, respectively.

**Swim bout generation**

To generate the characteristic zebrafish burst and glide locomotion pattern (**Fig. 2a, Extended Data Fig. 3b-e**), the amplitude of simulated tail oscillations must increase rapidly at the onset of the bout and then slowly decrease[44]. To achieve such tail undulation, the '*Bout gate*' allows activation of all the twelve central pattern generation oscillators (*CPGs*) in the simulated spinal cord and generates symmetric tail undulations. After the initialization of *CPG* activation, the amplitude of *CPGs* behaves like an exponential function dampened by a low-pass filter. This configuration produces a simulated tail-beating amplitude with a quick rise and slow decay. A bout is terminated if the amplitude decreases to less than 10% of the initial value of the exponential function ($E_{CPG0}$).

$$E_{CPGt} = E_{CPG0}(a + b_t)^{t-t_0} \tag{12}$$

$$A_{CPGt} = \beta A_{CPG(t-T)} + (1 - \beta)E_{CPGt} \tag{13}$$

In equations (13) and (14) $E_{CPG0}$ is the initial amplitude of *CPG*, $a$ is a decrement factor, and $b_t$ is a Gaussian noise sample. This Gaussian noise can induce randomness in the bout duration and bout distance. Furthermore, $t_0$ is the initial moment of the bout, $\beta$ is a parameter of the lower-pass filter, and its value is between 0 and 1. In our simulation, we set $E_{CPG0}$ , $a$ , $b_t$ , and $\beta$ to a value of 1.2, $2.73 \times 10^{-7}$ , $\sim 4.16 \times 10^{-10} N(0,1)$, and 0.965, respectively.

During a routine turn bout (**Fig. 2a, Extended Data Fig. 3e**), the *simZFish* begins the bout with a tail bias (i.e., a higher curvature) to either leftward or rightward during the first undulation, similar to real larval zebrafish[68].

To achieve this behavior in the simulated *simZFish*, we implemented left and right *vSPN* neurons, representing a class of real, ventrally located spinal projection neurons in the hindbrain[68]. These simulated *vSPNs* project directly to the motor neurons on the ipsilateral side[69]. Similar to the time evolutions of real zebrafish tail-beating amplitudes, the bias of the tail increases rapidly at the onset of the bout and then slowly decreases[44]. Consequently, we also used an exponential function dampened by a low-pass filter to describe the output of *vSPNs.*

$$E_{vSPNt} = (E_{vSPN0} + b_t)a^{t-t_0} \tag{14}$$

$$A_{vSPNt} = \beta E_{vSPN(t-\mathrm{T})} + (1-\beta)E_{vSPNt} \tag{15}$$

where $E_{vSPN0} + b_t$ is the initial amplitude of a *vSPN* neuron. For instance, to generate a leftward bout, we set the $E_{vSPN0}$ of the left vSPN as 1 and the $E_{vSPN0}$ of the right *vSPN* as 0. Furthermore, $b_t$ is the Gaussian noise used to induce the randomness of the turning angle. $\beta$ is a parameter of the low-pass filter, and $t_0$ is the initial moment of the bout. In the *simZFish* simulation, we set $a$, $b_t$, and $\beta$ to a value of $3.37 \times 10^{-16}$, $\sim 0.16|N(0,1)|$, and 0.935, respectively. The generation of a forward swim bout is similar to routine turning bouts, but with the $E_{vSPN0}$ of both *vSPNs* set to 0 (**Extended Data Fig. 3c**). All the other parameters were fixed (identical values) for both forward and turning bouts.

**Central pattern generators (*CPGs*)**
To execute tail undulations, we modeled the spinal locomotor *CPGs* as a system of coupled phase oscillators similar to other modeling studies, such as the lamprey[70]. Here, we use a double chain of coupled oscillators following our previous study[71] (shown in equations (16) and (18)). We implemented a pair of these oscillators in each of *simZFish* six tail segments.

$$\dot{\theta}_{ijt} = 2\pi v_{ij} + \sum_{kl} \sin(\theta_{klt} - \theta_{ijt} - \Phi_{ij-kl}) \quad (16)$$

$$x_{ijt} = 1 + \sin(\theta_{ijt}) \quad (17)$$

In the equations presented above, $\theta_{ijt}$ is the state variable indicating the phase of the oscillator in *i*-th segment and *j* side (*j* can be left or right). $x_{ijt}$ is the oscillatory output of the oscillator that will be sent to the motor neurons (see below). $v_{ij}$ is the intrinsic frequency of the oscillator in *i*-th segment and *j* side. Couplings are only between neighbor oscillators. Furthermore, $w_{ij-kl}$ and $\Phi_{ij-kl}$ are the weights and phase biases between oscillator $ij$ and oscillator $kl$ respectively. In the model, $w_{ij-kl}$ are set as 20 for all coupling, while $\Phi$ is set as $-\pi$ of the bilateral oscillators. For the neighbor oscillators along the chain, we set the $\Phi$ to a value of $0.2\pi$ for the descending couplings, and $-0.2\pi$ for the ascending couplings, to generate a traveling wave along the body.

***CPGs*, *nMLF* and *vSPN* to motor neurons**
The motor neurons in the two segments closer to the head receive the projections from *CPGs* and descending inputs, including the ipsilateral *vSPNs* and *nMLF* (**Fig. 2a**). We added the projection from *nMLFs* to motor neurons since the *nMLFs* modify the rostral tail bias in zebrafish[72,73]. The motor neurons in the third to sixth segments only receive the inputs from the *CPGs* and ipsilateral *vSPN* as expressed by

$$m_{Lit} = x_{Lit} + nMLF_L + A_{vSPNt}, i = 1, 2 \quad (18)$$

$$m_{Lit} = x_{Lit} + A_{vSPNt}, i = 3, 4, 5, 6 \quad (19)$$

where, in $m_{Lit}$, $Lit$ indicates the Left, *i*-th segment counting from the head, and time. Turning bouts are therefore generated by left-right asymmetries of both *nMLFs* and *vSPNs* centers, with *nMLFs* influencing the increased curvature of only the two most rostral joints, and *vSPNs* the increased curvature of all joints.

**Motor neurons to servomotor command**
The neural circuit's final output is represented in the activity of these motor neurons, which directly control the servomotors (**Extended Data Fig. 3b**). Thus, motor neurons and servomotors form the interface between the *simZFish* neural circuit model and its body. Here, for simplicity, we calculate the differences of the corresponding motor neurons on both sides of the body, to determine the desired angles for each actuated joint in the body, as

$$o_{it} = A_{seg}(m_{Lit} - m_{Rit}) \quad (20)$$

where $o_i$ is the angular output of the i-th segment, and $A_{seg}$ is the preset value of the amplitude of the undulation of each segment. We set the values of $A_{seg}$ as 0.20, 0.36, 0.48, 0.59, 0.70, and 1.00 for the segment 1 to segment 6, respectively. $m_{Lit}$ and $m_{Rit}$ are the respective left and right motor neurons in the *i*-th segment at time *t*. In the *simZFish*, the servomotor receives the inputs of the $\theta_{it}$ and rotate the linked segments.

**Data-driven updating simZFish*1.0* to *simZFish2.0***

As described in the main manuscript related to **Fig. 4**, we used the *simZFish1.0* simulation which was based on our previous neural circuit models derived from recorded behavior and neural responses to the orthogonal stimulus set to predict real zebrafish behavior for the parallel stimulus set. The observed differences suggested the missing computations, revealing nonrealistic connectivity in the *simZFish1.0* simulation. For instance, *simZFish1.0* did not account for suppression by monocular forward motion and a contralateral activating projection of neurons carrying information about 'backward' stimuli.

To update the neural network to match the new data, we considered at least two ways: (1) change the connectivity of existing (8 bilateral symmetric) *Pt* neurons or (2) add subclassify neuron types from experimentally measured neurons and adjust connectivity to improve the function in the network. While we could indeed fix some of *simZFish1.0* behavior by changing connectivity (data not shown), we reasoned that investigating how the real zebrafish brain handles these stimuli would lead to a more realistic model. Thus, we performed calcium imaging to record from neurons during presentation of the full orthogonal and parallel stimulus sets (**Extended Data Fig. 8a**). We observed that at least four of the previously identified response classes, which form the basis of motion computation in *simZFish1.0*, segregate into forward and backward selective subclasses, with additional neurons responding to both, with some showing strong suppression to the contralateral eye. This correlational evidence suggested that a reasonable approach would be to replace each of the previous classes with their subtypes, replicating their measured response properties. Therefore, to repair the *simZFish1.0* circuit we augmented the network by splitting each of these 4 neuronal classes into their most frequent subclasses. We then tuned the connectivity of these artificial subtypes to matching them to those recorded in real, live zebrafish brain (**Video 7**).

Therefore, *simZFish2.0* is the result of identifying new functional subtypes of real pretectal neurons in real zebrafish, providing data driven evidence for mechanism of how all motion stimuli interact. The updated *simZFish2.0* neural network thus integrates calcium imaging results for the *lPT* neurons in response to the parallel stimulus set. Their responses to forward backward visual stimulus combinations indicate that there exist forward and backward selective subtypes of at least four of the previously identified neural response types. We show the rightward *oB* subtypes in (**Extended Data Fig. 8b**). Therefore, inspired by these findings, we divided the *oB, B, Mm*, and *S* neurons into *oB1, oB2, B1, B2, Mm1, Mm2, S1,* and *S2* neurons, respectively, where the type 1 neurons preferentially respond to forward motion, whereas the type 2 neurons respond to backward motion (**Extended Data Fig. 8c**). The computations of the responses of the *lPT* neurons are:

$$x_{jt} = \frac{1}{1 + e^{-\omega(\sum_{ij} w_{ij} ePT_{jit} - \mathrm{b})}} \tag{22}$$

$$S1_{Lt} = \frac{1}{1 + e^{-5(\min\,(0.8ePT_{RSt}, ePT_{LIt}) + 0.5ePT_{RAt} + 1.5ePT_{LAt} - 2ePT_{RPt} + 1ePT_{LPt} - 0.4)}} \tag{23}$$

$$S2_{Lt} = \frac{1}{1 + e^{-5(\min(0.8ePT_{RSt}, ePT_{LIt}) - 1ePT_{RAt} - 1.5ePT_{LAt} + 1ePT_{RPt} + 1.5ePT_{LPt} - 0.4)}} \tag{24}$$

where, $x$ indicates an *lPT* neuron ($x \in \{oB1, oB2, B1, B2, iB, ioB, iM_M, M1_M, M2_M, oM_L\}$), $j$ can either be left or right, $t$ indicates the current instant, $w_{ij}$ is the weight associated with the ePT neuron of type $i$ ($i \in \{S, A, I, P\}$) in the $j$ side. Finally, $\omega$ and $b$ are the parameters of the sigmoid transfer function. Equations (23) and (24) allow to compute the responses of the $S1$ and $S2$ neurons in the left pretectum at instant $t$ (analogous formulas are used to compute the activations of the $S1/S2$ neurons in the right *PT*).

Unbiased cluster analysis of the neurons revealed that some *IPT* neurons fall into specific subclasses with type 1 *IPT* neurons, responding to forward motion, and type 2 *IPT* responding to backward motion (**Fig. 4g, Extended Data Fig. 8**). In their projection to the *nMLF* neurons, the type 1 *IPT* neurons, which respond to forward motion, have excitatory effects. Conversely, the type 2 *IPT* neurons exert an inhibitory effect on the *MLFs* given by

$$nMLF_{Lt} = \beta nMLF_{L(t-T)} + (1-\beta)\frac{1}{1+e^{-6.5(0.1oB1_L - 0.1oB2_L + 0.32B1_L - 0.08B2_L + 0.25iB_L - 0.05\,ioB_L + 0.6iM_{ML} + 0.4M1_{ML} + 0M2_{ML} - 0.8oM_R + 0.3S1_L - 0.1S2_L - 0.4)}} \quad (25)$$

In their projection to the *aHB* neurons, we split their connection weights with manual modifications to align the simulation behavior more closely with that of the animal. For example, in *simZFish1.0*, the connection weights between the $oB_L$ neuron to the $\mathrm{nMLF}_L$ is 0.0. In *simZFish2.0*, we split the $oB_L$ neuron into $oB1_L$ and $oB2_L$. Since $oB1_L$ responds to forward motion, we set a positive connection weight with a value of 0.1 to $\mathrm{nMLF}_L$. Since $oB2_L$ responds to the backward motion, we set a negative connection weight with a value of -0.1 to the $\mathrm{nMLF}_L$. All parameters for *simZFish1.0* and *simZFish2.0* are provided in text files with the simulation software.

#### *simZFish Source code documentation*

The complete code to run the simZFish robot within the Webots simulator is available at https://ponyo.epfl.ch/proj/zebrafish/simzfish. The repository also contains a user guide ('simZFish_User_Instructions.pdf') to facilitate the code's installation and execution.

#### *Optic flow analysis*

For optic flow analysis in **Extended Data Fig. 5**, **6**, video clips from *simZFish* simulation experiments were created by recording the simulation's left and right camera views, corresponding to the visual input to the right and left simulated *simZFish* retina, respectively. Each video was recorded while *simZFish* was held stationary in the center of the virtual petri dish at a fixed distance from the display, while identical visual sinusoidal grating stimuli were presented below. In other camera view videos, we analyzed for optic flow (**Extended Data Fig. 6a**, **b**), we presented moving dot patterns or used video from recordings during virtual river experiments related to (**Fig. 2g**). Videos for lens comparisons were captured as *simZFish* simulation experiments with three different lens angles with diagonal angles of 90° (f ≈ 0.707mm), 120° (f ≈ 0.577mm), and 150° (f ≈ 0.268mm). All other experiments were recorded with the 120° lens. These short focal lengths are calculated under the generous assumption that the diagonal retinal sensor size is 1 mm. In reality, a 6-day-old zebrafish lens is even smaller, with a maximal lens size of 150 µm[74], likely causing some barrel distortion for even the 90° lens. Nonetheless, these recordings represent a significant advance in testing the effect of sensory morphology, such as the optical properties of the eye's lens, on network function. Sinusoidal grating patterns either moved orthogonally (perpendicular to the fish's head-to-tail orientation) as in the converging, inward stimulus or parallel to the body axis (forward motion stimulus). RGB videos were recorded at a slowed to 0.2x speed, over 1200 frames seconds long, 25 fps.

For optic flow computation analysis, we converted each image to greyscale, cropping all camera views to 236 * 640 pixels. Using custom *Matlab* scripts, we applied the *Horn-Schunck* method to compute the optic flow fields across these *simZFish* videos. This global optic flow technique estimates the optic flow by iteratively solving an optic flow constraint equation, introducing a global smoothness term across the entire image. We chose the Horn-Schunck approach over other local methods like *Lucas-Kanade* because it handles global motion stimuli, e.g., uniform light intensities of the sinusoidal grating stimuli[75]. Indeed Lucas-Kanade method resulted in optic flow estimates that appear to predict no or little optic flow in the peripheral visual field. Thus, for each stimulus and lens condition, we calculated optic flow at each pixel location per frame using the Horn-Schunck algorithm. We plotted every 5$^{th}$ vector, which visualized the estimated motion magnitude and direction, emphasizing that most motion detection occurs around the translating dark light grating edges and none in the featureless sky. Heatmaps of these calculated flow vector fields were plotted using a circular (HSV) color map (**Extended Data Fig. 5b**). The circular mean angle for the lower anterior and posterior quadrants was calculated across 1000 frames (**Extended Data Fig. 5c**), discarding the first frame (which cannot be used as this method compares two consecutive frames). We then further analyzed the magnitude of optic flow estimates across these quadrants to compare the effect of different focal lengths on optic flow perception (**Extended Data Fig. 5d**). Finally, we analyzed optic flow data across 1000 frames to detect deviations from the expected translational motion patterns (e.g., orthogonal motion), corresponding to perceived reversed motion likely due to the 'wagon wheel illusion' effects introduced by discrete sampling of our periodic stimulus pattern[76]. Notably, the unnaturally narrow lens (90°) causes the strongest perception of the true direction of motion and significant reverse optic flow (**Fig. 3b**, light grey trace). This is likely due to antialiasing, a wagon wheel illusion effect expected for periodic stimuli with discrete sampling[76]. While the strong activation via the narrow 90° lens appears to indicate greater functionality at first, possibly the contaminating perception of the reverse optic flow for the 90° lens and the fact that the higher, more fish-like lens (150°) results in higher bout frequencies suggests superior OMR for the biologically realistic lens (**Extended Data Fig. 6c**).

***Additional calcium imaging analysis methods***

To identify functional subtypes, we first classified all neurons as before as responsive to a set of 8 monocular, leftward, and rightward drifting stimuli (see **Video**). Next, we applied unbiased hierarchical clustering to all neurons falling in these specific classes, identifying subclasses within (**Fig. 4f**, **Extended Data Fig. 6**). For instance, the responses to the set of eye-specific forward-backward stimuli of *oB* neurons were previously unknown. Via clustering of responses in this class, we discovered that this class contains a significant proportion of forward selective *oB* and backward selective *oB* subclasses. Similarly, other subtypes also fell into forward and backward selective classes, displaying suppression when the other eye is presented with the other direction. These neurons, which respond to monocular forward or backward motion but show reduced activity when these stimuli are shown simultaneously, match, in principle, the behavioral results. Detailed, documented analysis workflow for the calcium imaging data analysis and hierarchal clustering, including all parameters, can be found at *https://github.com/Naumann-Lab/caImageAnalysis*

### 3. Supplementary Tables 1 to 3

| | Unit (Before) | Value (Before) | Unit (After) | Value (After) |
|---|---|---|---|---|
| Approx. volume of *simZFish* | $m^3$ | 0.23 $10^{-9}$ | $cm^3$ | 0.23 $10^{-3}$ |
| Gravity | $m \cdot s^{-2}$ | 9.81 | $cm \cdot s^{-2}$ | 981 |
| Water Viscosity | $kg \cdot m^{-1} \cdot s^{-1}$ | 0.001 | $kg \cdot cm^{-1} \cdot s^{-1}$ | 0.00001 |
| Water Density | $kg \cdot m^{-3}$ | 1000 | $kg \cdot cm^{-3}$ | 0.001 |
| Moment of inertia | $kg \cdot m^2$ | 1 | $kg \cdot cm^2$ | 10000 |

**Supplementary Table 1**. **Unit conversion for *simZFish* body simulations.**

| | $\omega$ | $b$ | $w_{LS}$ | $w_{LA}$ | $w_{LI}$ | $w_{LP}$ | $w_{RS}$ | $w_{RA}$ | $w_{RI}$ | $w_{RP}$ |
|---|---|---|---|---|---|---|---|---|---|---|
| $oB_L$ | 2.5 | 0.8 | 0.0 | 0.25 | 1.0 | 0.5 | 0.8 | 0.25 | -1.0 | 0.5 |
| $B_L$ | 2.5 | 0.8 | -0.8 | 0.4 | 1.0 | 0.5 | 0.8 | 0.4 | -1.0 | 0.5 |
| $iB_L$ | 2.5 | 0.8 | -0.8 | 0.0 | 1.0 | 0.25 | 0.8 | 0.0 | 0.0 | 0.25 |
| $ioB_L$ | 2.5 | 0.8 | 0.0 | 0.0 | 1.0 | 0.0 | 0.8 | 0.0 | 0.0 | 0.0 |
| $iM_{ML}$ | 4.0 | 0.6 | 0.0 | 0.4 | 1.0 | -0.4 | 0.0 | 0.4 | 0.0 | -0.4 |
| $M_{ML}$ | 4.0 | 0.6 | 0.0 | 0.75 | 1.0 | 0.25 | 0.0 | 0.75 | -1.0 | 0.25 |
| $oM_{ML}$ | 5.0 | 0.4 | 0.0 | 0.15 | 0.0 | 0.0 | 0.8 | 0.15 | 0.0 | 0.0 |
| $S_L$ | 4.0 | 0.6 | 0.0 | 0.75 | 1.0 | 0.25 | 0.8 | 0.75 | 0.0 | 0.25 |
| $oB_R$ | 2.5 | 0.8 | 0.8 | 0.25 | -1.0 | 0.5 | 0.0 | 0.25 | 1.0 | 0.5 |
| $B_R$ | 2.5 | 0.8 | 0.8 | 0.4 | -1.0 | 0.5 | -0.8 | 0.4 | 1.0 | 0.5 |
| $iB_R$ | 2.5 | 0.8 | 0.8 | 0.0 | 0.0 | 0.25 | -0.8 | 0.0 | 1.0 | 0.25 |
| $ioB_R$ | 2.5 | 0.8 | 0.8 | 0.0 | 0.0 | 0.0 | 0.0 | 0.0 | 1.0 | 0.0 |
| $iM_{MR}$ | 4.0 | 0.6 | 0.0 | 0.4 | 0.0 | -0.4 | 0.0 | 0.4 | 1.0 | -0.4 |
| $M_{MR}$ | 4.0 | 0.6 | 0.0 | 0.75 | -1.0 | 0.25 | 0.0 | 0.75 | 1.0 | 0.25 |
| $oM_{MR}$ | 5.0 | 0.4 | 0.8 | 0.15 | 0.0 | 0.0 | 0.0 | 0.15 | 0.0 | 0.0 |
| $S_R$ | 4.0 | 0.6 | 0.8 | 0.75 | 0.0 | 0.25 | 0.0 | 0.75 | 1.0 | 0.25 |

**Supplementary Table 2**. **Neural connectivity matrix for *simZFish1.0*.**

The parameters and synaptic weights compute the responses of the *lPT* neurons for *simZFish1.0* neural network; in particular, $\omega$ and $b$ are the parameters of the sigmoid transfer function, and $w_{L/R\ S/A/I/P}$ indicate the weight associated with the *ePT* neuron in the left pretectum. The subscript letters *S, A, I, and P* indicate the type of *ePT* neuron receiving input from direction-selective ganglion cells preferring either superior, anterior, inferior, or posterior motion. The rows in the table are related to the various *lPT* neurons composing *simZFish1.0* neural network, replicating experimentally observed neural activation of neurons in real zebrafish[61].

| | $\omega$ | $b$ | $w_{LS}$ | $w_{LA}$ | $w_{LI}$ | $w_{LP}$ | $w_{RS}$ | $w_{RA}$ | $w_{RI}$ | $w_{RP}$ |
|---|---|---|---|---|---|---|---|---|---|---|
| $oB1_{Lt}$ | 2.5 | 0.8 | 0.0 | 1.0 | 1.0 | -1.5 | 0.8 | 1.5 | -1.0 | -2.0 |
| $oB2_{Lt}$ | 2.5 | 0.8 | 0.0 | -1.0 | 1.0 | 1.5 | 0.8 | 0.0 | -1.0 | 0.0 |
| $B1_{Lt}$ | 2.5 | 0.8 | -0.8 | 1.0 | 1.0 | 0.0 | 0.8 | 1.0 | -1.0 | -2.0 |
| $B2_{Lt}$ | 2.5 | 0.8 | -0.8 | -1.0 | 1.0 | 2.5 | 0.8 | 0.0 | -1.0 | 0.0 |
| $iB_{Lt}$ | 2.5 | 0.8 | -0.8 | 0.0 | 1.0 | 0.5 | 0.8 | 0.0 | 0.0 | 0.0 |
| $ioB_{L}$ | 2.5 | 0.8 | 0.0 | 0.0 | 1.0 | 0.0 | 0.8 | 0.0 | 0.0 | 0.0 |
| $iM_{MLt}$ | 4.0 | 0.6 | 0.0 | 0.8 | 1.0 | 0.0 | 0.0 | 0.0 | 0.0 | -0.8 |
| $M1_{MLt}$ | 4.0 | 0.6 | 0.0 | 1.0 | 1.0 | -1.0 | 0.0 | 1.5 | -1.0 | -1.0 |
| $M2_{MLt}$ | 4.0 | 0.6 | 0.0 | -1.5 | 1.0 | 1.5 | 0.0 | 0.0 | -1.0 | 0.5 |
| $oM_{MLt}$ | 5.0 | 0.4 | 0.0 | 0.0 | 0.0 | 0.0 | 0.8 | 0.3 | 0.0 | 0.0 |
| $S1_{Lt}$ | 4.0 | 0.6 | 0.0 | 1.5 | 1.0 | -1.0 | 0.8 | 0.5 | 0.0 | -2.0 |
| $S2_{Lt}$ | 4.0 | 0.6 | 0.0 | -1.5 | 1.0 | 1.5 | 0.8 | -1.0 | 0.0 | 1.0 |
| $oB1_{Rt}$ | 2.5 | 0.8 | 0.8 | 1.5 | -1.0 | -2.0 | 0.0 | 1.0 | 1.0 | -1.5 |
| $oB2_{Rt}$ | 2.5 | 0.8 | 0.8 | 0.0 | -1.0 | 0.0 | 0.0 | -1.0 | 1.0 | 1.5 |
| $B1_{Rt}$ | 2.5 | 0.8 | 0.8 | 1.0 | -1.0 | -2.0 | -0.8 | 1.0 | 1.0 | 0.0 |
| $B2_{Rt}$ | 2.5 | 0.8 | 0.8 | 0.0 | -1.0 | 0.0 | -0.8 | -1.0 | 1.0 | 2.5 |
| $iB_{Rt}$ | 2.5 | 0.8 | 0.8 | 0.0 | 0.0 | 0.0 | -0.8 | 0.0 | 1.0 | 0.5 |
| $ioB_{R}$ | 2.5 | 0.8 | 0.8 | 0.0 | 0.0 | 0.0 | 0.0 | 0.0 | 1.0 | 0.0 |
| $iM_{MRt}$ | 4.0 | 0.6 | 0.0 | 0.0 | 0.0 | -0.8 | 0.0 | 0.8 | 1.0 | 0.0 |
| $M1_{MRt}$ | 4.0 | 0.6 | 0.0 | 1.5 | -1.0 | -1.0 | 0.0 | 1.0 | 1.0 | -1.0 |
| $M2_{MRt}$ | 4.0 | 0.6 | 0.0 | 0.0 | -1.0 | 0.5 | 0.0 | -1.5 | -1.0 | 1.5 |
| $oM_{MRt}$ | 5.0 | 0.4 | 0.8 | 0.3 | 0.0 | 0.0 | 0.0 | 0.0 | 0.0 | 0.0 |
| $S1_{Rt}$ | 4.0 | 0.6 | 0.8 | 0.5 | 0.0 | -2.0 | 0.0 | 1.5 | 1.0 | -1.0 |
| $S2_{Rt}$ | 4.0 | 0.6 | 0.8 | -1.0 | 0.0 | 1.0 | 0.0 | -1.5 | 1.0 | 1.5 |

**Supplementary Table 3**. **Neural connectivity matrix for *simZFish2.0*.** This table summarizes the parameters and the weights used to compute the responses of the *lPT* neurons in the left pretectum for *simZFish2.0* ; in particular, $\omega$ and $b$ are the parameters of the sigmoid transfer function, and $w_{L/R\ S/A/I/P}$ indicates the weight associated with the *ePT* neuron in the left pretectum, with S, A, I, P in subscript specifying the type of *ePT* neuron, selective for superior, anterior, inferior, posterior direction selective preference, respectively. The rows in the table are related to the *lPT* neurons of the updated *simZFish2.0* neural network, which includes the newly identified response subtype neurons found in the real zebrafish (**Fig. 4f**).

### 4. Supplementary Figures and Legends:

#### Extended Data Fig. 1 | Larval zebrafish inspired *simZFish* body design.

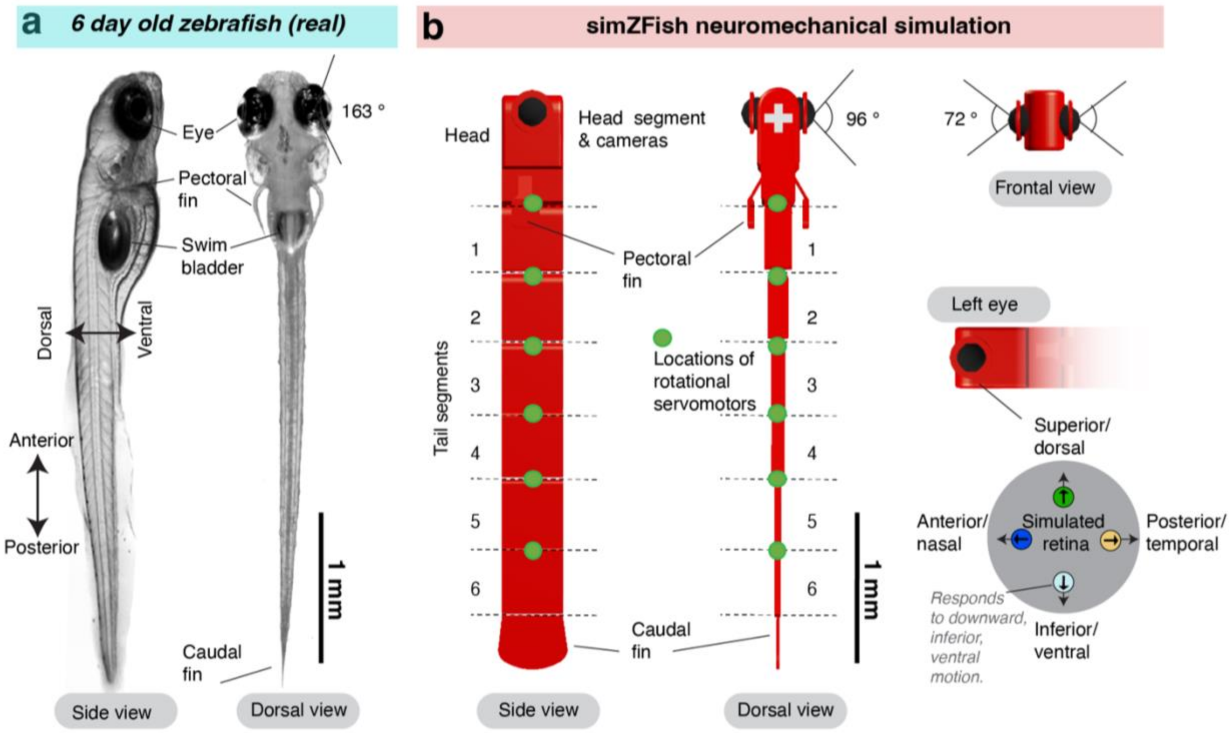


c

| Segment | Dimension (mm) | | | Mass (g) | drag Force Coefficients | | | Viscous Resistance Force Coefficient |
|---|---|---|---|---|---|---|---|---|
| | Length | Height | Width | | Longitudinal | Lateral | Vertical | |
| Head | 0.8 | 0.44 | 0.266 | 9.38E-8 | 0.05 | 0.15 | 0.25 | 2 |
| Seg 1 | 0.48 | 0.44 | 0.198 | 4.25E-8 | 0 | 0.2 | 0.25 | 0 |
| Seg 2 | 0.48 | 0.44 | 0.148 | 3.08E-8 | 0 | 0.2 | 0.25 | 0 |
| Seg 3 | 0.48 | 0.44 | 0.104 | 2.21E-8 | 0 | 0.25 | 0.25 | 0 |
| Seg 4 | 0.48 | 0.44 | 0.094 | 1.90E-8 | 0 | 0.3 | 0.25 | 0 |
| Seg 5 | 0.48 | 0.44 | 0.055 | 1.10E-8 | 0 | 0.4 | 0.25 | 0 |
| Seg 6 | 0.48 | 0.44 | 0.046 | 9.00E-9 | 0 | 0.5 | 0.25 | 0 |
| Caudal fin | 0.32 | 0.44 | 0.022 | 3.00E-9 | 0 | 0.75 | 0.25 | 0 |

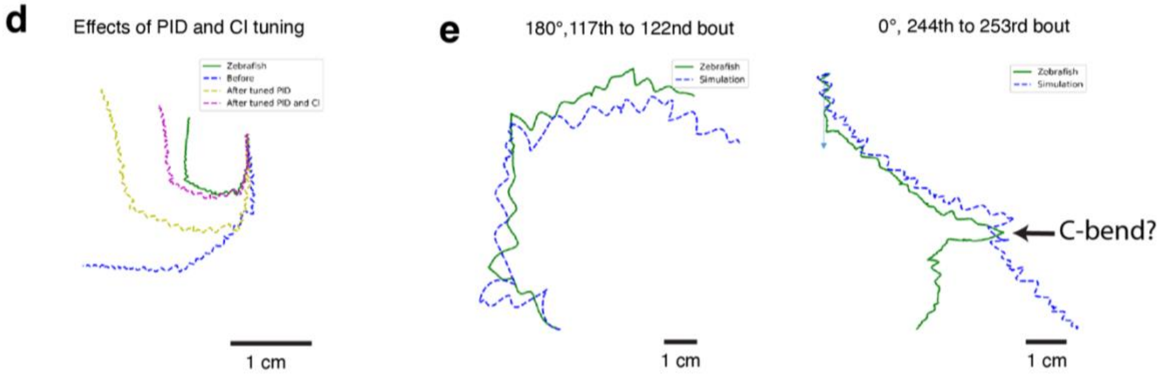

**Extended Data Fig. 1 | Larval zebrafish inspired *simZFish* body design.**

**a** Side and dorsal view of an agarose-embedded 6-day-old live zebrafish. According to Easter et al., 1996, the distance between the left and right optical centers of each eye has been previously described as 458 μm, resulting in a visual field of 163°[77].

**b** 3D rendering of the *simZFish* body, consisting of a head segment, six tail segments, and a caudal fin. The caudal fin attaches to the last segment (Segment 6). We equipped the *simZFish* with six servomotors (green markers) on the tail to enable tail movements to propel. Two RBG cameras are positioned laterally in the head segment, replicating the real zebrafish eye positions. All simulation experiments, except stated otherwise, were performed with rectilinear lenses, resulting in an angle of views of 96° horizontally, 72° vertically, and 120° diagonally. *Right*, laterally positioned *simZFish* eyes contain direction-selective retinal ganglion cells (*DSGCs*) in four variations, each responding to four cardinal directions of motion across the eye.

**c** Parameters of dimension, mass, drag Force Coefficients, and viscous Resistance Force Coefficient of each segment. We set the dimensions and weights to match measurements of real larval zebrafish measured by Zhao et al., 2020[39]

**d** Comparison of example swim trajectory of a real zebrafish and *simZFish* with different tuning parameters for the drag force coefficient (Cl) and PID controller.

**e** Comparisons of example swim trajectories of zebrafish[44] and *simZFish* with the kinematic sequences of full field backward (180°) and forward (0°) stimulation. While we find that *simZFish* and real zebrafish show a qualitative match for most behaviors seen during OMR, such as routine swims and turns, the behavioral kinematic repertoire of *simZFish* is limited compared to real behavior. For instance, for the forward (0°) stimulation, the real zebrafish performs a fast escape type movement, called C-bend, with a high angle, which is not matched by the simulation. Although the real fish and the simulation perform rightward bouts, the simulated one has a smaller amplitude.

**Extended Data Fig. 2 | *simZFish* interface enables limitless simulated experimental designs.**

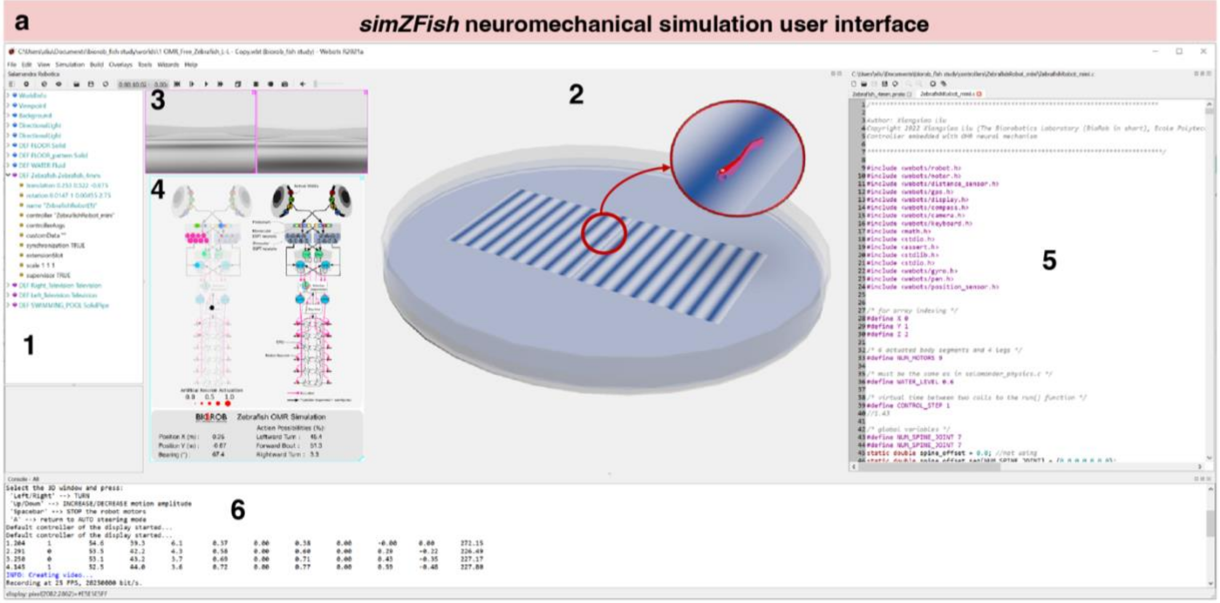


1. Items in simulation (e.g. *simZFish*, petridish)
2. Simulated zebrafish OMR experimental environment (*simZFish*)
3. Camera views from simZfish
4. User interface showing the internal neuronal activation
5. Software Program editor of *simZFish*
6. Console outputs of the software program

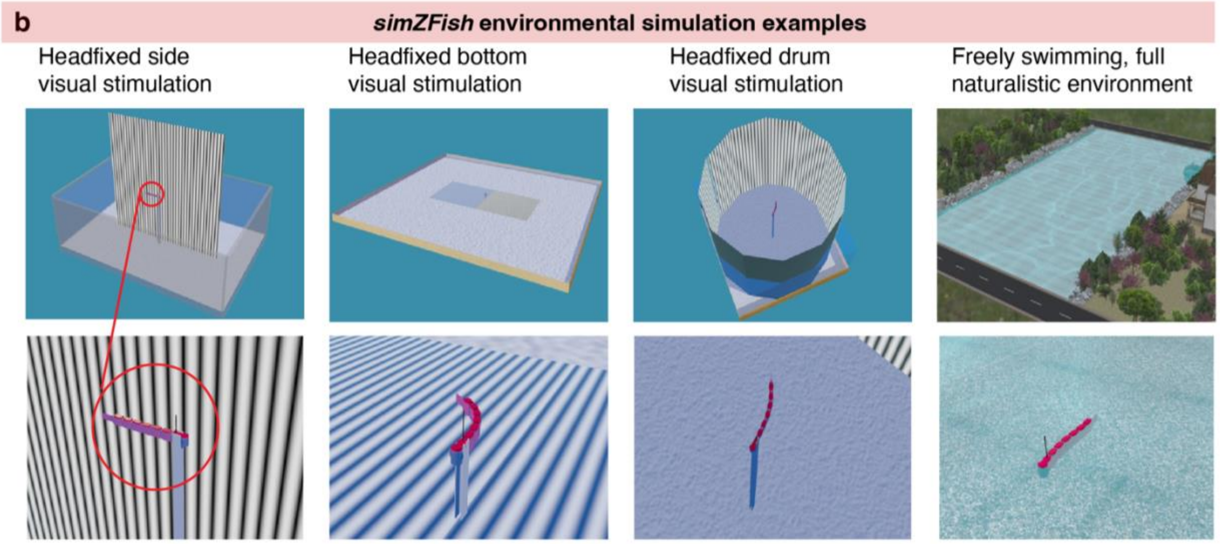

**Extended Data Fig. 2 | *simZFish* interface enables limitless simulated experimental designs.**

**a** Screenshot of the *simZFish* graphical user interface (GUI) on the Webots platform. When running *simZFish* simulation experiments, the GUI displays various simulated items to provide ongoing information, e.g., current camera views and current neural activity states, etc. Importantly, users can visualize *simZFish* internal states, including the neural activity of all simulated neurons and all behavioral variables, and test various scenarios, e.g., present visual stimuli in different directions in a preset trial structure.

**b** Screenshots of variations of different experimental designs, testing the response of *simZFish* in different situations requiring many days of experimental troubleshooting for real zebrafish and new hardware and software designs. For instance, while testing head-fixed *simZFish* next to a screen with different visual stimulation patterns would require minimal programming, performing similar experiments in the real fish would require agarose embedding of a larval zebrafish on a suspended translucent platform and mounting of a physical projector. Thus, we can use *simZFish* to test various standard lab tests (head fixed bottom, drum stimulation) far faster than in real life. Also, the simulation nature of *simZFish* allows us to test scenarios that are impossible to perform in real life.

**Extended Data Fig. 3 | *simZFish* bout generation and neural connectivity tuning.**

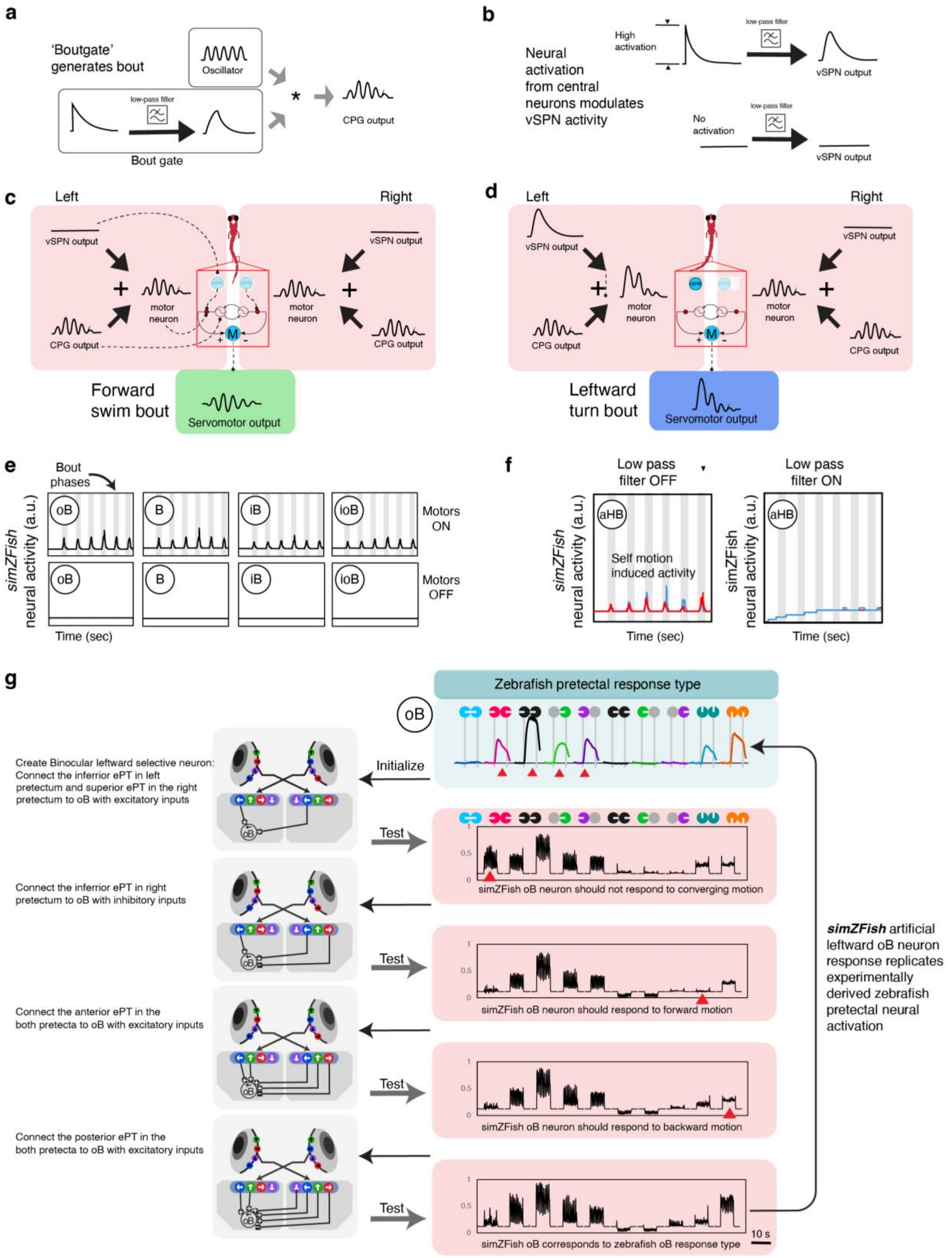

**Extended Data Fig. 3 | *simZFish* bout generation and neural connectivity tuning.**

**a** To initiate several cycles of tail beating during a *simZFish* burst and glide swim, the '*Bout gate*' determines the amplitude of the oscillator's output. The amplitude is defined as an exponential function passed through a low-pass filter.

**b** *Top*, illustration of the output of the activated ventral spinal projection neuron (*vSPN*). *Bottom*, silent *vSPN*. To establish the amplitude of an activated *vSPN*, we set the initial value ($E_0$) of an exponential function through a low-pass filter. For the silent *vSPNs,* we set the initial value ($E_0$) of the exponential function to zero. To introduce variations in the turning angle, we set $E_0$ as a sum of a constant and an absolute value of Gaussian noise.

**c** During a forward bout, left and right *vSPNs* are nearly silent. The *CPG* output predominantly defines the activation of motor neurons. The difference between the left and right motor neurons determines the angle of the servomotor output. The servomotor produces periodic and undulatory rotation to propel the *simZFish* body forward during swimming.

**d** To initiate a leftward turning bout, the left *vSPN* generates a highly activated output, leading to a strong activation of the left motor neuron compared to the right motor neuron. This difference induces a biased undulatory rotation of the servomotor output in the left direction, prompting the *simZFish* to execute a leftward turn.

**e** *simZFish* pretectal neural activity during swimming. When motors are switched on, and *simZFish* generates swim bouts (grey bars), *lPT* neurons are activated by self-generated visual stimulation, even without visual motion presentation. When motors are off, as expected, no self-generated visual stimulation is registered by these neurons.

**f** *simZFish* aHB (right, red; left, blue) neurons are activated by self-generated visual stimulation during swimming (grey bars). Low pass filtering reduces swim bout-induced activity.

**g** All neuronal connections were initialized according to synaptic weights proposed by our previously published average best fit model[61]. We manually tuned the neuronal connections that were not suggested in previous studies, such as the connections between early pretectal (*ePTs*, receiving direct input from DSGCs) and later pretectal neurons (*lPTs*, receiving input from *ePTs*) to match recorded pretectal neurons in real, live zebrafish[61]. This process entailed an iterative refinement of neural connections, achieved by first recording initialized *simZFish* neural responses to various monocular and binocular visual stimuli (symbols as in **Fig. 1f**) and subsequently comparing them with real zebrafish neural responses (top right). For example, the initialized connectivity for a leftward, outward selective binocular pretectal neuron ($oB_L$) resulted in erroneous responses to converging motion (red triangle). After updating the connectivity, here adding inhibitory connections from an inferior selective motion sensitive *ePT*, we measured the neural responses to validate an improved match of simulated and real observed zebrafish neuronal activity. Through this iterative approach, we obtained neuronal connections that result in artificial neural responses like those of zebrafish neurons.

**Extended Data Fig. 4 | *simZFish* replicates zebrafish OMR behavior.**

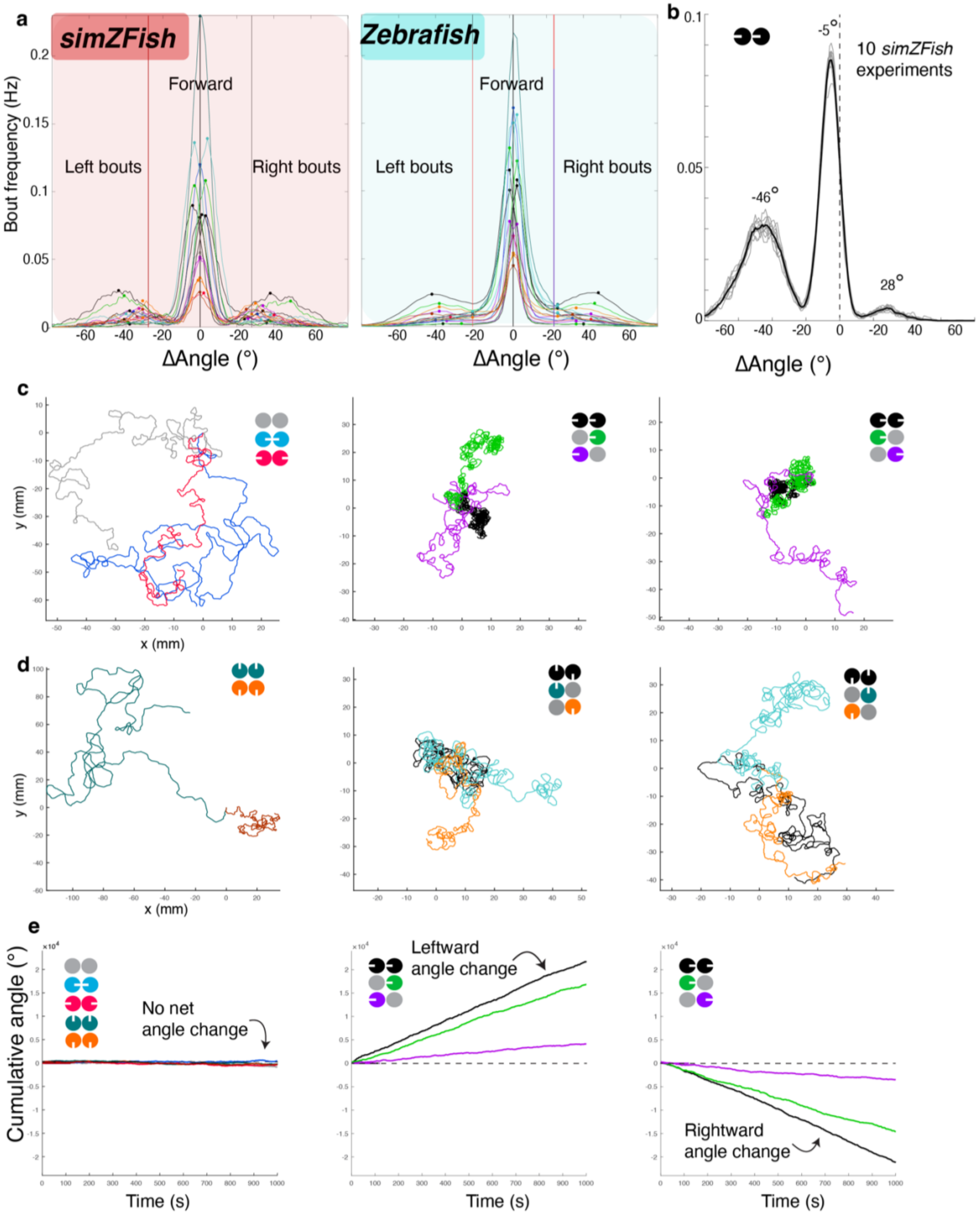
a
simZFish
Zebrafish
Forward
Left bouts
Right bouts
Bout frequency (Hz)
ΔAngle (°)
b
-5°
10 simZFish experiments
-46°
28°
ΔAngle (°)
c
y (mm)
x (mm)
d
e
Cumulative angle (°)
No net angle change
Leftward angle change
Rightward angle change
Time (s)

**Extended Data Fig. 4 | *simZFish* replicates zebrafish OMR behavior.**

**a** *Left*, *simZFish(red)* behavioral summary plotted as absolute bout frequency histograms for all 17 motion stimuli, for 2000 second presentation of both the orthogonal stimulus set (**Fig. 1f**) and parallel stimulus set (**Fig. 4B**). Dots mark the maximal amplitude for the left, center, and right peak for each stimulus histogram. Note the different peak locations. *Right*, real zebrafish average absolute bout frequency histogram for all 17 stimuli. Vertical lines in both plots show limits for bout classifications for leftward, forward, and rightward bouts for bout probability plots in **Fig. 2f**.

**b** Average *simZFish* bout frequency histogram (black) for 10 repetitions (grey thin lines) of 2000 seconds presentation of binocular leftward motion. As for real zebrafish, binocular leftward motion strongly increases left turns to the direction of optic flow (left peak, -46 degrees) and biased forward swims (center peak, -5 degrees) while suppressing turns in the opposite direction (rightward, positive angles) of optic flow (right peak, 28 degrees).

**c** X, Y trajectories for 2000 seconds *simZFish* behavior during continuous, closed-loop stimulation with monocular and binocular motion stimulation with gratings moving orthogonally (stimulus set as in **Fig. 1f**). Note that *simZFish* covers large distances for stimuli, not increasing turning in one direction (left panel), converging (blue), diverging (red), stationary gratings (grey). For binocular (black) or medial (green) left or right stimuli, *simZFish,* just like real zebrafish, responds with tight circling, lateral stimuli (purple) less so.

**d** X, Y trajectories for 2000 seconds *simZFish* behavior during continuous, closed-loop stimulation with monocular and binocular motion stimulation with gratings moving parallel (**Fig. 4b**).

**e** Cumulative angle plots for 1000 seconds of *simZFish* behavior during continuous, closed-loop stimulation with monocular and binocular motion stimulation with gratings moving orthogonally (stimulus set as in **Fig. 1f**).

**Extended Data Fig. 5 | Optic flow analysis of visual stimuli seen through the eyes of *simZFish*.**

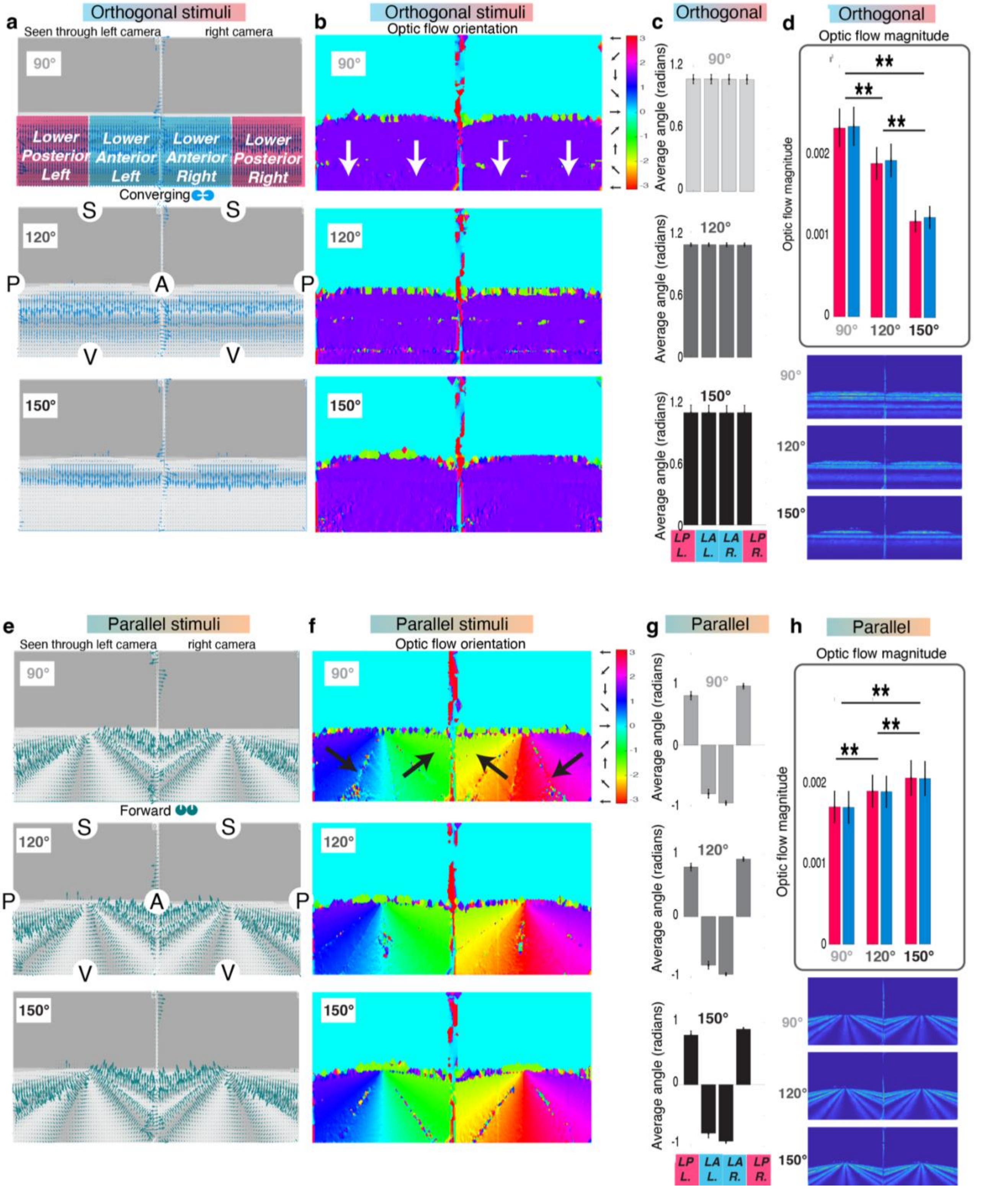

**Extended Data Fig. 5 | Optic flow analysis as seen through the eyes of *simZFish*.**

**a** Snapshots of left and right visual scene as seen through *simZFish* cameras while viewing converging sinusoidal gratings, i.e., left and right eye see medial motion. Calculated optic flow with the Horn- Schunck method is represented as vectors (blue). The top (90°) views delineate the lower visual field quadrants most relevant to motion processing, as the 'sky' is featureless in these idealized conditions. Lower posterior left (LPL), lower anterior left (LAL) for the left eye, and lower anterior right (LAR), lower posterior right (LPR) for the right eye are the quadrants with significant optic flow (length of arrows). Optic flow estimates in the upper quadrants are too small to be seen. The 150° snapshot illustrates the perspective distortion that renders the grating's 'stripes' larger towards the ventral visual field, reducing optic flow. Anterior (A), posterior (P), superior (S), ventral (V).

**b** Flow orientation analysis of stimulus seen by right and left *simZFish* camera while viewing converging motion when equipped with 90°, 120°, and 150° lenses. All lower visual field quadrants show optic flow analysis largely moving ventrally (purple hues, see circular colormap). Some opposite or reverse direction optic flow (green hues) is measured due to antialiasing, creating a 'Wagon wheel illusion' effect.

**c** Average flow orientation (circular mean) of each lower quadrant of a 1000-frame movie for simZFish viewing converging motion, when equipped with 90°, 120°, and 150° lenses, shows that flow orientation stays consistent across retinal quadrants and for different focal lengths for stimuli moving orthogonally. Error bars are standard deviations. Paired Wilcoxon. **: <= 0.0001, n = 1000)

**d** Flow magnitude analysis of 1000 frame movie for *simZFish* viewing converging motion when equipped with 90°, 120°, and 150° lenses, showing flow orientation stays consistent across retinal quadrants and for different focal lengths for stimuli moving orthogonally. Error bars are standard deviation. Paired Wilcoxon. **: <= 0.0001. *Bottom*, heat maps of optic flow magnitude with 90°, 120°, and 150° lenses; note that the 150° lens shows reduced magnitude of the perceived optic flow.

**e** Snapshots of left and right camera views while *simZFish* views forward sinusoidal gratings, optic flow represented as vectors (teal). Abbreviations as in **a**. Optic flow estimates display a rotational vector field with posterior perceiving net downward and anterior net upward optic flow information.

**f** Flow orientation analysis of right and left *simZFish* camera while viewing forward motion when equipped with 90°, 120°, and 150° lenses. Note smooth rotational optic flow field in lower quadrants.

**g** Average flow orientation (circular mean) of each lower quadrant of a 1000-frame movie for *simZFish* viewing forward motion, when equipped with 90°, 120°, and 150° lenses, show rotational effect consistent across different focal lengths for forward gratings. Error bars represent standard deviations.

**h** Flow magnitude analysis of 1000 frame movie for *simZFish* viewing forward motion when equipped with 90°, 120°, and 150° lenses, showing flow orientation stays consistent across retinal quadrants and for different focal lengths for stimuli moving orthogonally. Error bars represent standard deviation. Paired Wilcoxon. **: <= 0.0001, n = 1000)

**Extended Data Fig. 6 | Effects of embodiment on *simZFish* neural activity and behavior.**

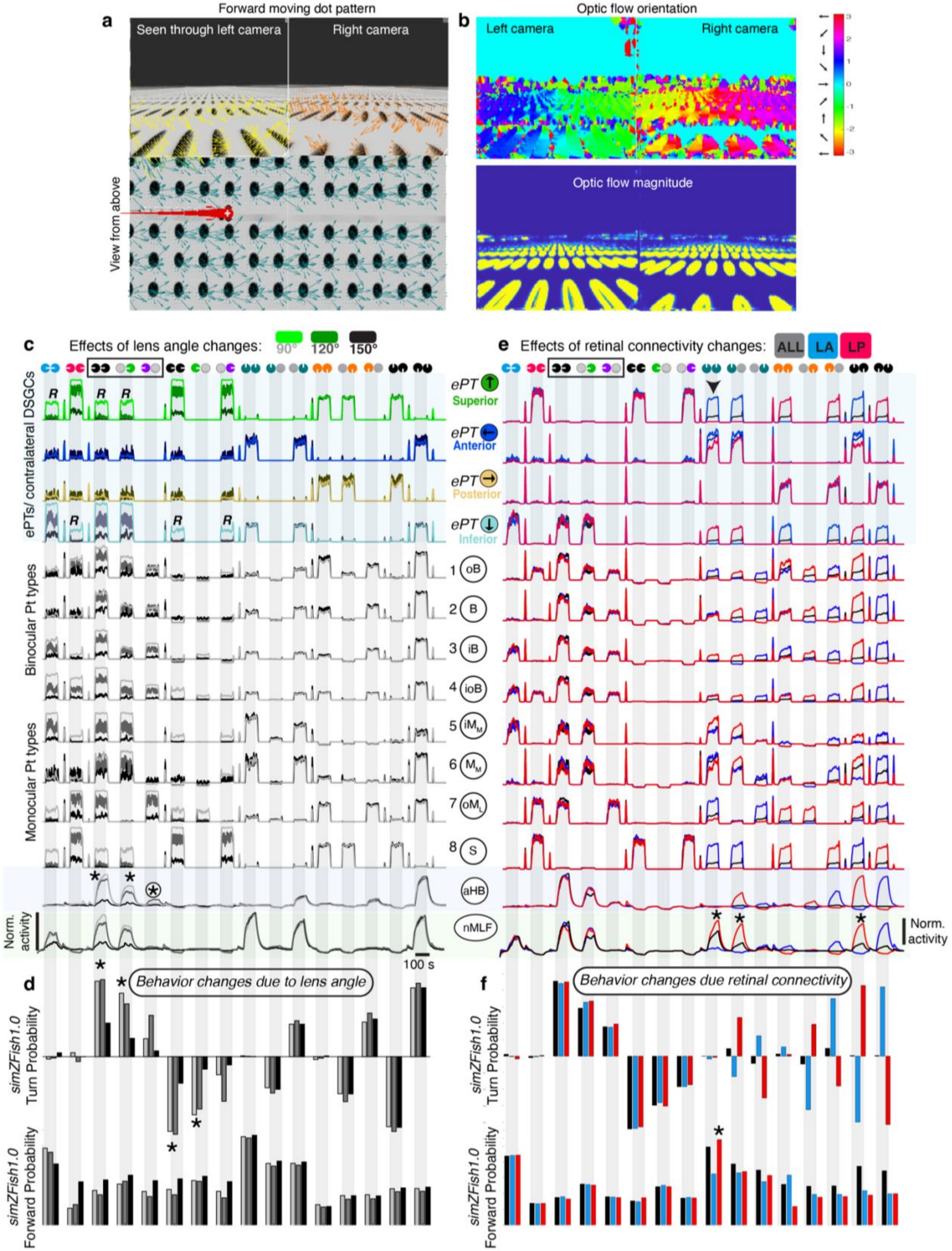

**Extended Data Fig. 6 | Effects of embodiment on *simZFish* neural activity and behavior.**

**a** Left and right camera views of *simZFish* when viewing a moving dot stimulus. Vector fields from optic flow analysis show that more complex stimuli, like dot patterns, similar to sinusoidal gratings, result in rotational optic flow when moving parallel to the body axis.

**b** Optic flow orientation and magnitude for two consecutive frames of 'dot forward motion' as in **a**.

**c** Complete left hemisphere *simZFish1.0* network output when equipped with 90°, 120°, and 150° lenses in response to 16 visual motion stimuli indicated on top (circular symbols represent the left and right eye, with white notches the direction of motion presented). The top four traces show output from left hemisphere *ePTs,* which receive direct input from right eye *DSGCs. ePTs* are colored to indicate direction selectivity of their upstream *DSGCs*: Anterior (A, dark blue), posterior (P, yellow), superior (S, green), ventral (V, light blue). Note that the brightest color traces (90°) indicate that DSGCs are activated most by stimuli viewed through the narrow-angle lens but also perceive reverse motion (R). These activations are inherited throughout the neural network, resulting in the strongest activation of *aHB* neurons during 90° viewing, marked with asterisk (*).

**d** *simZFish1.0* behavioral output, provided as the probability to generate a forward, left, or right bouts when equipped with 90°, 120°, or 150° lenses in response to 16 visual motion stimuli indicated on top. While the unnatural narrow-angle lens (90°) results in stronger turning, larger, more biorealistic 150° lenses result in higher forward bout probabilities. Note the strong turning predictions for monocular forward or backward motion.

**e** Complete left hemisphere *simZFish1.0* network output, with different *DSGCs to ePTs* connectivity in response to 16 visual motion stimuli indicated on top (circular symbols represent the left and right eye, with white notches the direction of motion presented). Traces show the normalized, artificial neural activity for when all (black), only lower posterior (*LP*, red), or lower anterior (*LA*, blue) *DSGCs* were connected to contralateral *ePTs*. Note that *iMM* neurons, the most prevalent monocular neuronal type, respond most to *LP* connectivity.

**f** *simZFish1.0* behavioral output, provided as the probability to generate forward, left, or right bouts While the unnatural narrow-angle lens (90°) results in stronger turning, larger, more biorealistic 150° lenses result in higher forward bout probabilities. Note the strong turning predictions for monocular forward or backward motion. *simZFish1.0* predicts very strong, opposing behavioral turning responses to monocular and binocular forward-backward stimulation, which prompted us to record real zebrafish behavior to these stimuli (**Fig. 4**, **Extended Data Fig. 7**).

**Extended Data Fig. 7 | Zebrafish behavioral measurements suggest *simZFish1.0* updates.**

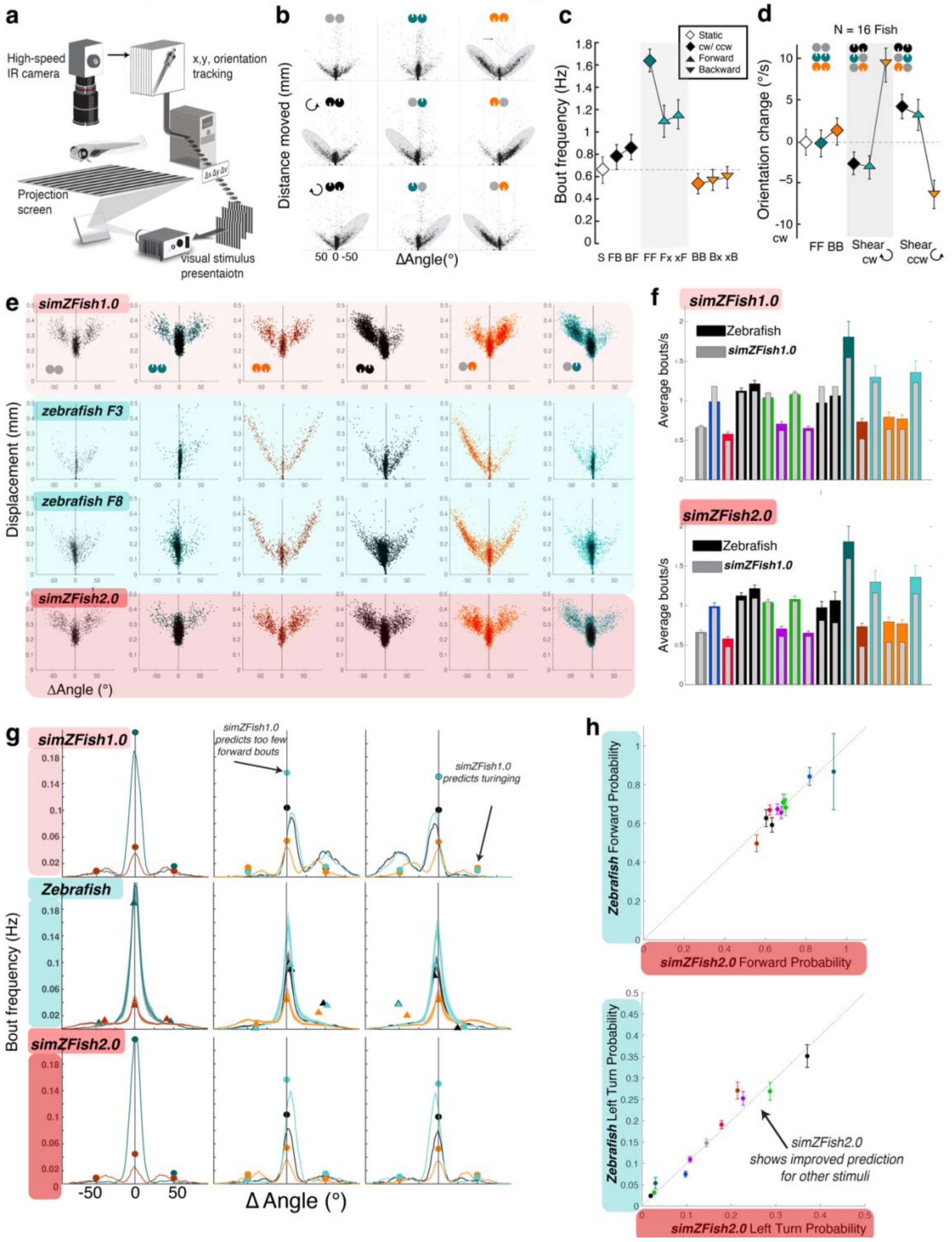

**Extended Data Fig. 7 | Zebrafish behavioral measurements suggest *simZFish1.0* updates.**

**a** Schematic of a close-loop behavioral rig used to monitor behavior in live zebrafish in response to visual motion stimuli.

**b** Scatterplots of all swimming bouts for an individual representative zebrafish, showing each bout's distance moved (y-axis) and angle (x-axis). Stimulus symbols are as in **Fig. 4b**: forward (F, teal) and backward (B, orange). Each dot represents a single locomotion event, plotting its distance and angle change. Grey outlines highlight increased routine turns to that side. Note interactions between forward and backward stimuli for shearing stimuli, indicating that backward motion information is suppressed by the simultaneous presentation of forward motion to the other eye.

**c** Average bout frequency for each stimulus. These averages show that forward motion to each eye combines to drive full, binocular forward bout frequency. N = 16 zebrafish. Error bars represent SEM across fish). On average forward motion drives, backward motion decreases bout frequency.

**d** Average angular velocity. Monocular backward causes fish to turn in the direction contralateral to the stimulated eye but is suppressed by forward to the other eye. Binocular forward (*FF*) vigorously increased forward bouts, whereas binocular backward (*BB*) elicited turning in either direction, reducing the number of bouts. Monocular forward (*xF*, *Fx*) responses approached 50% of the bout rate of *FF*, suggesting that forward is approximately linearly integrated across eyes. In contrast, *BB* suppressed the overall bout rate, with monocular backward (xB, Bx) showing slightly less bout suppression. (*xB*, *Bx*) also induced turning in the contralateral direction of the stimulated eye, suggesting that, despite containing no formal directional information, the brain interprets this as a left or rightward stimulus. Yet, when forward and backward are presented to either eye (*BF*, *FB*), resembling rotational motion, fish barely increased their bout frequency above spontaneous rates. These results indicate complex interactions across eyes and motion directions.

**e** Scatterplots for all locomotion bouts for *simZFish1.0 (top), simZFish2.0. (bottom)*, and two other example zebrafish recordings collapsed for monocular and combined forward and backward stimuli.

**f** *Top*, comparison of average bouts frequency for *simZFish1.0* (grey bars) versus zebrafish (colored bars). *Bottom*, same comparison for *simZFish2.0.* N = 16 zebrafish. Error bars represent SEM across fish.

**g** Average bout histograms, comparing. *simZFish1.0*, real zebrafish (shaded area represents SEM across fish), and *simZFish2.0.* Dots plotted on simulation data represent average zebrafish peak data for the left, forward, and right bout peaks. Triangles on the zebrafish plot (middle) are peak bout frequencies of *simZFish1.0*, showing specific mismatch, especially for predicting turning for monocular backward or forward motion.

**h** Probability plots comparing average bout probability for forward or turn bouts in *simZFish2.0* and real zebrafish (N = 32 zebrafish. Error bars represent SEM across fish) colors as in **Fig. 2**.

**Extended Data Fig. 8 | Zebrafish neural activity measurements suggest *simZFish1.0* updates.**

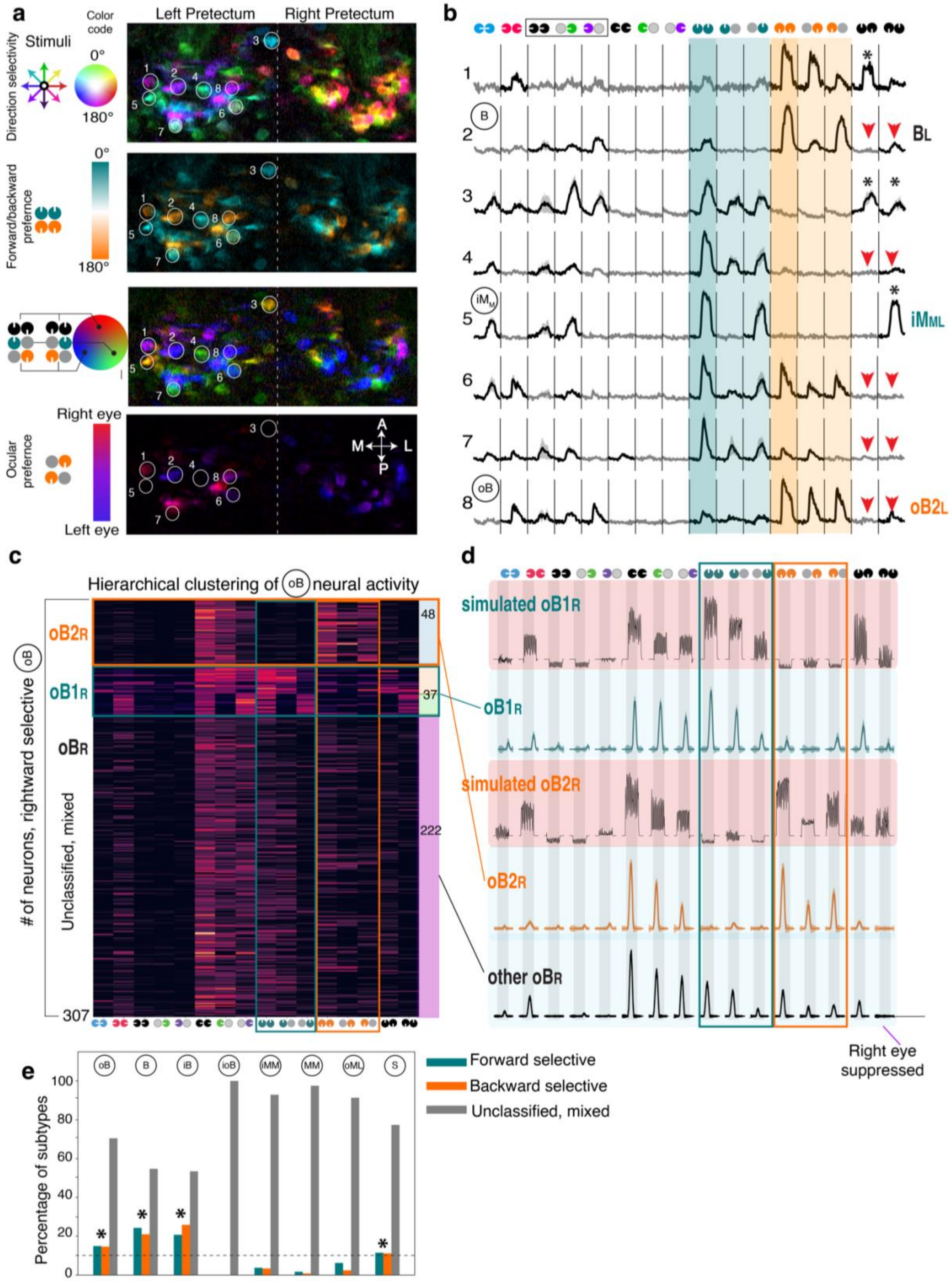

**Extended Data Fig. 8 | Zebrafish neural activity measurements suggest *simZFish1.0* updates.**

**a** Two-photon micrographs of an example plane of a 6-day-old *Tg(elavl3:H2B-GCaMP6s)* zebrafish brain showing motion-responsive neurons of the left and right pretectum. ΔF/F pixel-wise neural activity maps are color-coded for direction preference or other stimulus combinations. From top: *Direction selectivity:* directional tuning is lateralized, e.g., leftward tuned neurons are enriched in the left pretectum; spectral hue top plot indicates direction selectivity corresponding to the color wheel. *Forward-backward selectivity*: neurons are found in both hemispheres, with many neurons exhibiting a strong preference for either, indicated by many orange (backward) or teal (forward) neurons. *Binocular interactions*: Neurons exhibit different suppression levels when other stimuli are shown, e.g., ROI 4 is strongly activated by forward motion presented to either eye but is silenced by backward stimuli. *Ocular selectivity*: Backward-responsive neurons in the right pretectum tend to be activated by left eye,

**b** ΔF/F traces of the eight numbered regions of interest neurons (white circles) corresponding to **a**, showing diverse visual activation profiles. For example, neuron #4 is strongly activated by forward motion but suppressed by presenting backward motion to the other eye (red arrowheads). Neuron #5 is not suppressed (black asterisk). These binocular responses and interactions are ubiquitous, e.g., backward-selective neurons can be suppressed by forward motion to the other eye. In addition, clear preferences for forward or backward (#8, backward selective *oB*) can be observed for previously classified as *oB* neurons. Note that the *iMM* neuron (ROI 5) is an exclusively right-eye responsive neuron with strong, exclusive responses to forward and medial motion presented to the right eye, supporting the *LP* circuit hypothesis (**Fig. 3e**).

**c** Heatmap of all 307 rightward selective *oB* neurons in our data set, sorted by hierarchical clustering, also results in clear forward (middle, teal, 37) and backward selective (middle, orange) oB classifications as in **Fig. 4**.

**d** Average cluster response of all rightward selective *oB* neurons to various motion stimuli, interleaved with simulated *simZFish2.0* neural response of these subtypes.

**e** Fractions of subtypes within the eight original response classes found in the pretectum of real zebrafish. Four response types, *oB*, *B*, *ioB*, and *S* neurons, show a significant fraction (together over 20%, dashed line) of neurons exclusively responsive to either forward (teal) or backward (orange) motion in addition to mixed or unclassified responses (grey) to the monocular or binocular parallel motion stimuli. These subtypes were added to provide superior functionality of *simZFish2.0*. *: p-value <= 0.001, Wilcoxon signed-rank test.

**Extended Data Fig. 9 | Physical zebrafish-inspired *ZBot* replicates OMR neural activation and behaviors.**

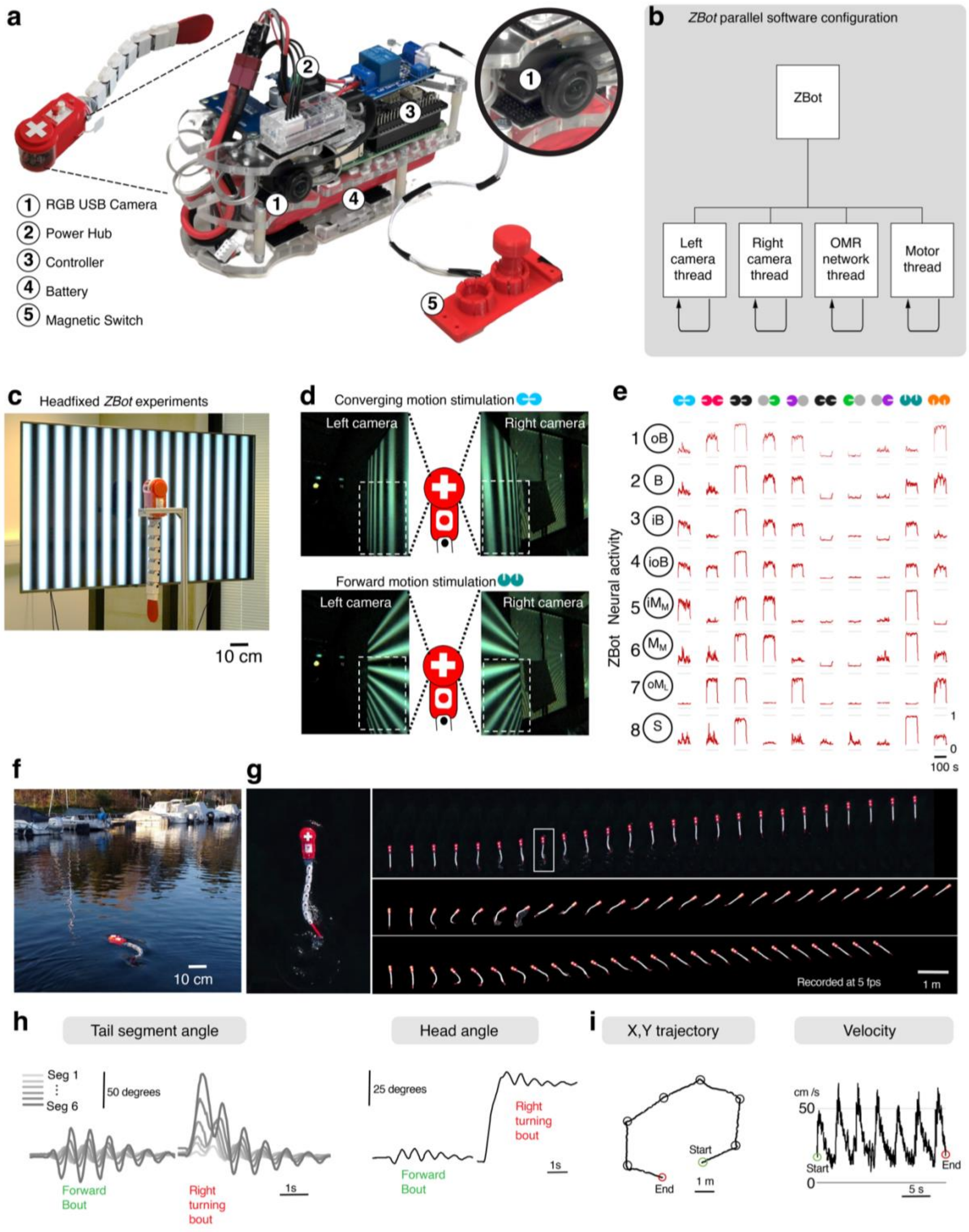

**Extended Data Fig. 9 | Physical zebrafish-inspired *ZBot* replicates OMR neural activation and behaviors.**

**a** The *ZBot* head segment contains a controller module with two RGB USB cameras positioned laterally, like real zebrafish eye positions. It also includes a power hub responsible for distributing power to both the controller and servomotors, a controller to implement the OMR program, a battery to supply power, and a magnetic switch to turn the robot on or off.

**b** Simplified diagram of the *ZBot's* software configuration, designed to enhance information processing through parallel computation. It incorporates four threads: two camera threads for camera processing, one OMR neural circuit thread for OMR implementation, and one motor control thread for controlling the robot's movements.

**c** We set up a 75-inch display placed vertically in a room. A mechanical frame holds the robot upon the display panel at about 20 cm. To reduce the robot's vibration, we manually turned off the servomotors. During the experiments, the display presented visual stimulation to the *ZBot*. To reduce the light reflection on the screen panel, we turned off the lights during the experiments.

**d** The left and right camera views of the *ZBot* in response to converging motion are shown. The dashed box indicates the visual field that drives DSGCs to drive the OMR network. The left and right camera views of the robot in response to the forward motion are shown.

**e** We recorded the activation of *lPT* neurons in response to eight motion stimuli. In each trial, we presented the stimuli for about 100 seconds. The robotic neuronal activation is comparable to the recording of zebrafish pretectum neurons in response to similar visual stimulation.

**f** Photograph of the *ZBot* performing free swimming in stationary water in Lake Leman (Switzerland).

**g** *Left*, aerial, dorsal view of zoomed-in view of *ZBot* recorded via camera drone during forward bout (white box). *Right*, projection of snapshots taken 200 ms apart of *ZBot* of the bouts from the view of a camera drone (Top: forward bout; Middle: Rightward bout; Bottom: Leftward bout.

**h** Tail segment angles of a forward bout and a leftward bout.

**i** Head-direction of a forward bout and a leftward bout. Head position (left) and velocity (right) of seven bouts in a sequence. Circles in the left figure represent the start of each bout.

**Extended Data Fig. 10| Zebrafish-inspired *ZBot* performs OMR in a naturalistic flowing river.**

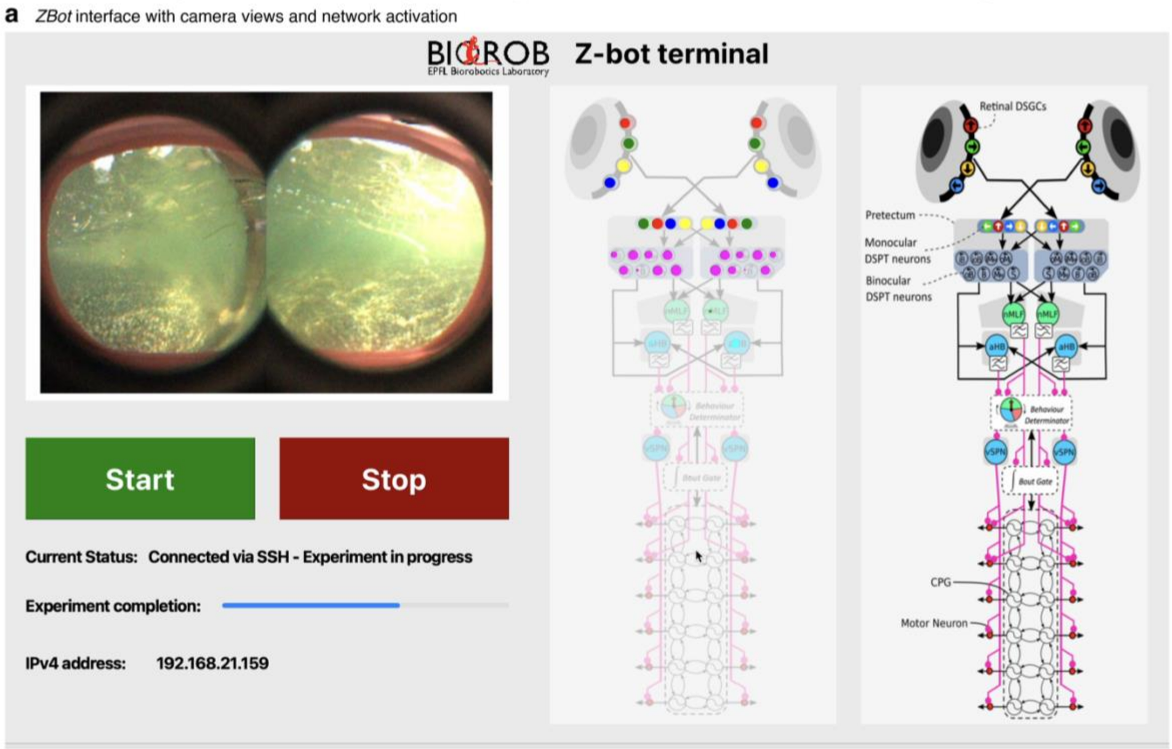


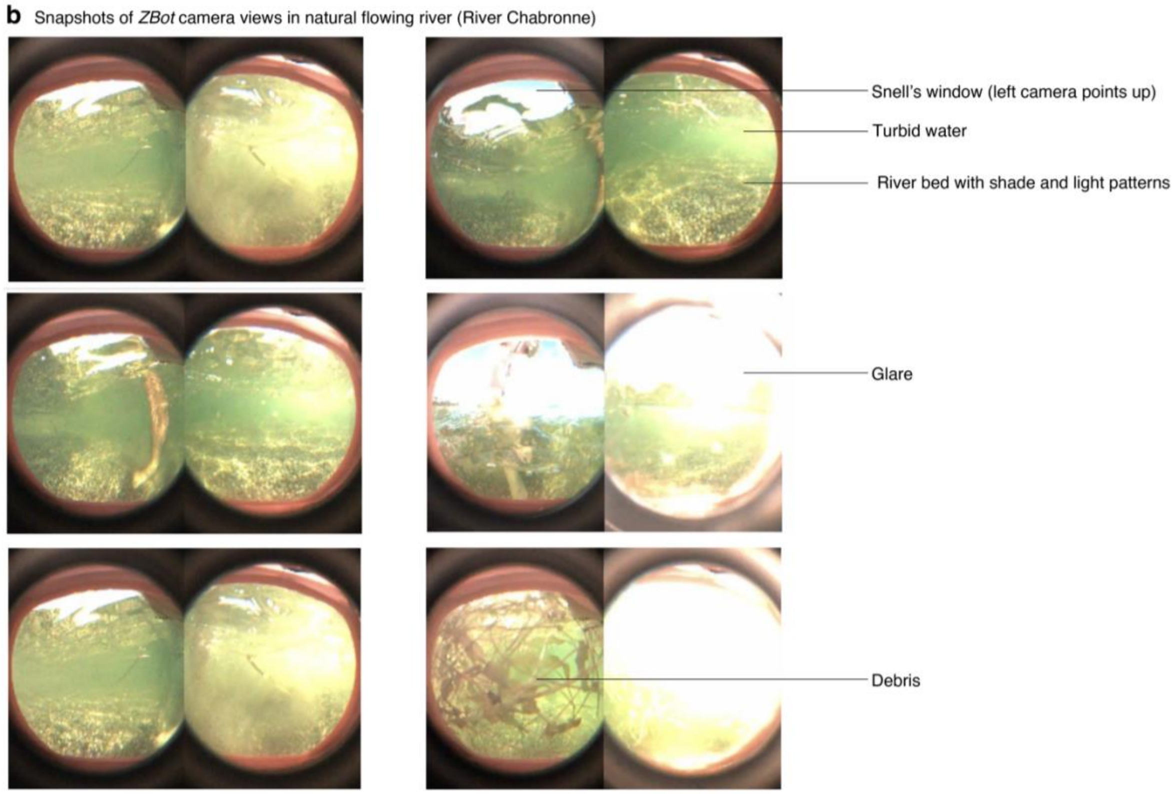

**Extended Data Fig. 10 | Zebrafish-inspired *ZBot* performs OMR in a naturalistic flowing river.**

**a** The *ZBot* interface shows left and right camera views, and neural network activation.

**b** Underwater snapshots of visual input as seen by *ZBot* during swimming in real river.

### 3. Videos

**Video 1 | *simZFish* experimental platform and graphical user interface**

Video of software graphical user interface (GUI) displaying a 45-degree dorsal view of the freely swimming *simZFish* in a simulated Petri dish, with simulated leftward moving sinusoidal grating stimuli presented from the bottom, slowed to 0.2x of the actual speed. The *simZFish* simulation allows inspection of the internal states, such as the camera views, artificial neuronal activation, and the state of the *simZFish* body, such as the servomotor positions. Visualizations can include artificial neural network activity, with each dot representing a neuron encoding activity as dot size on the graphical user interface (GUI) on the left. Leftward moving gratings activate the *DSGCs* of both eyes, neurons on the left, including the pretectal neurons, *aHB*, *NMLF*, and *vSPNs*, resulting in motor neuron activation driving locomotion. Note, s*imZFish* network activity includes self-generated visual stimulation due to its movement, activating the right pretectal neurons during a leftward bout. During the videos second part, the zoom allows a closer view of the simulated behavior. In the second part of the video, binocular forward motion is presented. Corresponds to **Fig. 2a.**

**Video 2 | *simZFish* bout and gliding swimming with recorded zebrafish kinematics**

Video of eleven consecutive bouts performed by *simZFish* using simulated hydrodynamics (red trace) plotted on the recorded behavioral trajectory of real zebrafish larva (black trace). The *simZFish* behavior was generated by applying the kinematics of a real zebrafish tail recorded as eight joint angles with a high-speed camera by Marques et al. 2018. Scale bar: 1 mm, video slowed to 0.1x of actual speed. Corresponds to **Fig. 2b.**

**Video 3 | *simZFish* swim and turn bouts**

The video shows a dorsal view of *simZFish* performing representative leftward and rightward turning bouts and a forward swim bout. The *simZFish* burst and glide swimming resembles the locomotion pattern observed in freely swimming, live zebrafish. Corresponds to **Fig. 2b.**

**Video 4 | *simZFish* maintains position in virtual water current**

Video of the *simZFish* equipped with OMR neural circuit capabilities when placed in a virtual river, with the water current dragging the *simZFish* to the right (illustrative example of the experiments in **Fig. 2g**). When the simulated water current drags the *simZFish* downstream (to the right), the OMR neural circuit is activated, enabling the *simZFish* first to orient upstream (head leftwards), aligning against the water flow direction, which is opposite of the optic flow. Like real zebrafish, *simZFish*, after initial alignment with the optic flow direction using turn bouts, *simZFish* swims straight at increased bout frequency to compensate for the perceived displacement in the virtual water, enabling *simZFish* to stay approximately in place. Corresponds to **Fig. 2g.**

**Video 5 | Optic flow analysis of *simZFish* visual input with different lens angles**

Video 5 shows right camera views of *simZFish* when equipped with lenses of different focal lengths corresponding to different viewing angles (90°, 120°, or 150°). The first video segment shows how lens angle affects local optic flow perception when viewing leftward orthogonal sinusoidal grating, with overlaid vector flow fields to visualize optic flow calculated across the visual field. When presenting such orthogonal moving patterns (e.g., right or leftward, medial or laterally moving stimuli), the lens' angle of view significantly influences the perceived optic flow with increasing magnitude for narrower angle lenses, as pattern elements closer to the eye appear to move faster, while those farther away move slower. The second video segment shows the same for a parallel moving sinusoidal pattern, forward moving gratings. Optic flow magnitude is similar for all angles of view (90°, 120°, or 150°). However, notice the rotational flow patterns for forward moving gratings. In contrast, this rotational flow pattern is not observed with orthogonal simulation, e.g., sinusoidal gratings moving left and right. *simZFish* This corresponds to the experiments shown in **Fig. 3a**.

**Video 6 | Impact of retinal connectivity on emergent neural responses**

Video 6 shows *simZFish*'s left and right camera view while presenting forward (0°) moving gratings, overlaid with vector flow field illustrating optic flow calculated across the visual field, extending from posterior (P) to anterior (A) and superior (S) to inferior (I). Visual stimuli with parallel motion, such as forward or backward-moving sinusoidal gratings, create a perspective-distorted projection of bottom-projected stimuli with apparent rotational optic flow. This causes different local optic flow directions in the lower anterior (LA, blue) and posterior (LP, magenta) visual field quadrants. As a result, direction-selective ganglion cells (*DSGCs*) in the *simZFish* and presumably the real zebrafish, as well as retinorecipient neurons located in the contralateral brain hemisphere i.e., early pretectal neurons (*ePTs*), demonstrate different activation levels depending on their *DSGCs* connections across the retina, especially to the lower anterior or posterior retina for patterns at the bottom of the field of view as viewed in a river. Corresponds to **Fig. 3e.**

**Video 7 |Neural activity in response to all visual motion stimuli combinations**

The video starts with a still image showing pixel-wise, color-coded neural responses for directional tuning, e.g., green neurons prefer forward motion, and red neurons prefer rightward motion. The video transitions into a movie of fluorescence intensity recorded with two-photon microscopy, showing calcium imaging of pretectal and hindbrain neurons in real, live larval zebrafish, expressing nuclear-targeted *GCaMP6s* under an *elval3* promoter. *Left*, mean dF/F fluorescence for five repetitions of motion stimulus presentations (symbols and colors as in **Fig. 4d**, gray circles indicate static phase before motion onset). *Right*, evolving normalized dF/F traces are shown for neurons from highlighted circular regions of interest, with the neurons belonging to forward, backward, and mixed *oB* subtypes. Beyond demonstrating the existence of forward, backward tuned *oB* neurons predicted by *simZFish*, these traces illustrate the diversity of dynamic neural responses across the pretectal area of live, real zebrafish brains. Corresponds to **Fig. 4d**.

**Video 8 | *ZBot* spontaneous swimming in stationary water**

Aerial video captured by a hovering drone of *ZBot* swimming in Lake Geneva. The video shows *ZBot* performing a forward swimming bout, a leftward, and a rightward turning bout. Corresponds to **Fig. 5c**.

**Video 9 | *ZBot* maintains position in a naturalistic flowing river using the OMR neural circuit**

This video starts with a short clip from the experimenter's perspective, showing the *ZBot* navigating against the river's flow direction using the OMR neural circuit. As the water current drags the *ZBot* downstream, the OMR neural circuit mechanism is activated by detecting optic flow in the opposite direction of the river's flow. Notice the rich visual stimuli of river rocks, floating leaves, and surrounding trees. The second video segment presents a collage of annotated aerial videos recorded by a flying drone, starting when the experimenter releases the *ZBot*. When the OMR mechanism is turned on, the *ZBot* demonstrates clear orienting bouts upstream and maintains its position within the camera's field of view for longer durations compared to when *ZBot* is blinded (with cameras off, swimming in random bouts) or when its motors are turned off. White tick marks indicate *ZBot's* location at a given corresponding time point after its release. This comparison shows how the OMR neural circuit enables the *ZBot* to react to the water current and maintain its position in a flowing natural river. Corresponds to **Fig. 5d**.
